# A quantum interface with mitochondrial bioenergetics

**DOI:** 10.64898/2026.07.03.736226

**Authors:** Parisa Aghaei, Sangjun Noh, Javier Noé Ramos-Silva, Minghao Jiang, Phang-Lang Chen, Yumay Chen, Fatemeh Goodarzinia, Robert Usselman, Philip Hemmer, Douglas C. Wallace, Peter J. Burke

## Abstract

Recent work has shown that genetically engineered proteins can serve as quantum bits in living systems. These quantum bits arise from the photochemistry of protein-bound flavins: blue-light excitation drives electron transfer to form a spin-correlated radical pair whose coherent singlet–triplet interconversion makes the protein’s fluorescence sensitive to weak magnetic fields. Because this radical-pair reaction depends on the redox state of the flavin—itself a central electron carrier in cellular metabolism—the magneto-fluorescence of a biological qubit is intrinsically coupled to the biochemistry around it. This suggests a powerful application of fundamental significance in biology, until now an unsolved problem in the field of quantum sensing. Here we show a new class of quantum sensor, mtMagLOV2, that interfaces directly to a defining feature of life itself: the bioenergetic state of the cell. We genetically engineer flavin mononucleotide (FMN)-containing, magnetic-field-sensitive fluorescent proteins (“biological qubits”) to be expressed and translocated into the key bioenergetic machinery of the cell: the mitochondrial matrix. Using confocal and super-resolution microscopy, mtMagLOV2 localizes to the mitochondrial cristae, home of the electron transport chain complexes I–V and ATP synthase—the site of oxidative phosphorylation (OXPHOS). By pharmacological manipulation of OXPHOS, we show that the sensor’s magneto-fluorescence tracks the redox (oxidation–reduction) state of the mitochondrial flavins, providing a quantum readout of redox status. The response differs between cancer cells (which rely heavily on glycolysis) and cardiomyocytes (which rely predominantly on OXPHOS), demonstrating “quantum bioenergetic profiling”. Together, these results establish biological qubits as quantum sensors capable of probing mitochondrial bioenergetics, opening a quantum window into the energetic machinery of living cells. More broadly, we anticipate that coupling quantum redox sensitivity to specific biochemical targets will extend the reach of quantum technologies across the life sciences.

## Introduction

Recently, biological qubits have been demonstrated by several groups worldwide ^1 2 3 4 5 6 7^. A biological qubit is a genetically engineered protein in which a photoexcited electron followed by an electron transfer event creates a (transient) entangled pair of spins between the donor and acceptor. The two-level system is the singlet vs. triplet state of the pair, called a radical pair (RP), whose energy level splitting depends on the static magnetic field. The occupation probability of this two-level system can be manipulated with radio frequency pulses at the spin spitting energy level divided by ℏ, typically in the GHz regime. Whether the pair is in the triplet or singlet quantum state changes the probability of the photoexcited electron relaxing to the ground state via fluorescence. Therefore, readout of the occupation of the quantum state can be achieved by fluorescence, analogous to the readout of solid-state spin qu-bits. These have all been demonstrated in cryogenically frozen cells to amplify coherent quantum state manipulation, live cells, and model organisms ^1 2 3 4 5 6^. Connecting functional biological qubits to biological processes will greatly expand the utility of quantum sensors in general, and biological qubit sensors in particular. In this paper, we demonstrate quantum sensing of oxidation-reduction (redox) reactions inside the mitochondria.

Oxidation-reduction (redox) reactions are essential to biological processes ^8 9 10 11 12 13^. A quantum interface to redox status would greatly increase the applications for quantum sensing in biology, providing a powerful platform for real-time physiological monitoring. In this paper, we report the development of a biological qubit sensor, genetically engineered to localize to the mitochondrion, mtMagLOV2. mtMagLOV2 utilizes the magnetic field dependent fluorescence of MagLOV2 ^1^ generated to detect spin-dependent singlet–triplet behavior to assess the mitochondrial flavin redox state. Using super resolution microscopy of live cells ^14^ we show the localization of mtMagLOV2 to the mitochondrial cristae the physical site of the mitochondrial OXPHOS apparatus. The mtMagLOV2 sensing on the electron transport chain redox status was confirmed by manipulating the electron transport chain electron density using specific OXPHOS inhibitors. This demonstrates the first quantum bit sensor capable of quantifying mitochondrial redox state, a fundamental parameter of classical bioenergetics.

## Results

### mtMagLOV2 design and flavins

Recent work by York, Ingaramo, and colleagues ^1 2 3^ has shown that the native protein AsLOV ^15 16 17^ can be evolved via directed evolution to display strong magnetic field dependent fluorescence, naming the new family of proteins MagLOV2 (Figure 1A). The general mechanism for this effect is believed to be the radical pair mechanism (Figure 1C), where an absorption event (shown in blue) excites a fluorophore (in this case FMN). As FMN starts in the singlet state, the excited FMN (denoted ^1^FMN*) is also in the singlet state. ^1^FMN* quickly converts to a triplet ^3^FMN* via inter system crossing due to the nuclear hyperfine interaction (ISC). Very shortly thereafter, an electron is transferred from a donor amino acid on the MagLOV protein FMN, forming a triplet spin state between the reduced FMN and the oxidized amino acid, called a radical pair. Due to the nuclear hyperfine interaction, the radical pair entangled state is split into singlet/triplet states (shown) which (coherently) evolve differently in a static magnetic field. This gives rise to magnetic field sensitive fluorescence, as the probability for the FMN* to emit a photon depends on the radical pair recombination probability, which in turn depends on whether it is in the singlet or triplet state. In MagLOV2, the non-covalently bound flavin cofactor chromophore ^18^ is the photoexcited and fluorescence species ^2 3^. Amino acids in the AsLOV protein^19^ serve as the donor in the entangled radical pair mechanism, and the FMN the acceptor. In this work, we target this self-contained quantum bit, with its own radical pair, directly into the mitochondria, and perform experiments with the mitochondria in different bioenergetic states.

**Figure 1.**
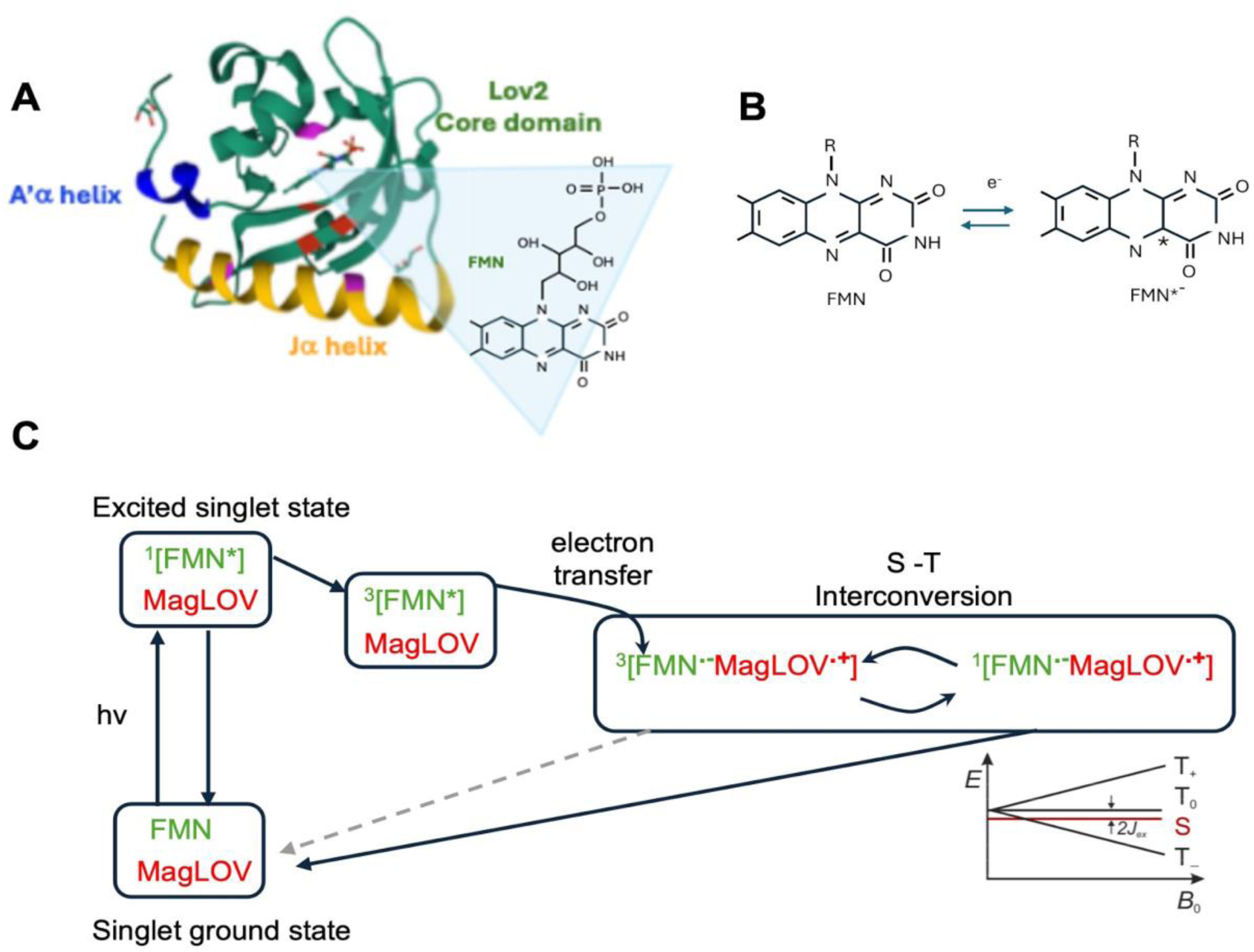
MagLOV2. (A) Structure of asLOV2 (similar to MagLOV2) protein (PDB structure 2V1A), showing non-covalently bound flavin mono-nucleotide (FMN). (The precise crystal structure of MagLOV2 is not known at this time.) (B) Redox states of FMN in solution. In the bound state, the electron donor/acceptor interaction occurs with amino acids of the MagLOV2 protein. (C) Quantum state diagram showing the spin-dependent singlet–triplet behavior, identifying the FMN as the fluorophore, and the amino acids (labeled generically as “MagLOV2”) as the donor/acceptor actors.

The magnitude of these spin-dependent magnetic field effects has been shown *in vitro* to be strictly modulated by the redox state of the flavin cofactor in multiple qubit contexts. Cohen et al ^6^ have shown the magnetic field dependent fluorescence (ΔF/F) in response to an alternating (chopped) static magnetic field of an analogous quantum bit in vitro depends on the redox state of a flavin cofactor. Similarly, solid state quantum bit sensors have also recently been demonstrated to sense cytochrome C redox status ^20^. Here, we propose that mtMagLOV2 will similarly track the real-time depletion or replenishment of this available oxidized FMN population (Figure 1B) in vivo as a function of the bioenergetic and redox status of the mitochondrial matrix and electron transport chain complexes. Thus, our hypothesis is that this qubit system, similar to those demonstrated *in vitro*, will *in vivo* demonstrate magnetic field dependent fluorescence that depends on the redox states of the soluble redox pairs such as FMN that serve as the cofactor of these particular qubits, the redox state of other soluble redox pairs in the matrix, and the redox status of the mitochondrial electron transport chain complexes, thus forming a quantum sensor of redox pairs integrated directly into the mitochondrion.

### MtMagLOV2 is localized to the mitochondria

In order to demonstrate localization of the mtMagLOV2 protein in its active, folded state inside the mitochondria, we used confocal microscopy to interrogate the location and fluorescence intensity of the mtMagLOV2 protein in live HeLa cells. MagLOV2 is known to fluoresce green, with blue excitation. Because mitochondria display endogenous cellular fluorescence in the same spectral region (due to endogenous mitochondrial flavins as well as lysosomes ^21^), we quantified the endogenous fluorescence and the fluorescence following transduction with mtMagLOV2 and showed that addition of mtMagLOV2 doubled the green fluorescence (See Supplementary Information). The second-generation mtMagLOV2-3F construct exhibited approximately 3-fold higher fluorescence intensity than the original mtMagLOV2 construct and approximately 6-fold higher fluorescence than wild type Hela cells autofluorescence under identical imaging conditions, resulting in improved signal-to-noise ratio during long-term imaging experiments.

In order to further demonstrate that folded, active mtMagLOV2 was localized in ***vital (live)*** mitochondria, we co-stained with the voltage dependent dye tetramethyl rhodamine ethyl ester perchlorate (TMRE). TMRE fluoresces in the red and so does not interfere with the green fluorescence of the mtMagLOV2 protein. Since TMRE only fluoresces if there is a sustained membrane potential ^14^, mitochondria glowing red are known to be viable. TMRE and mtMagLOV2 were found to co-localize (Figure 2), Pearson correlation coefficient of 0.87. Hence, the mtMagLOV2 quantum sensor is localized inside vital, live mitochondria.

**Figure 2.**
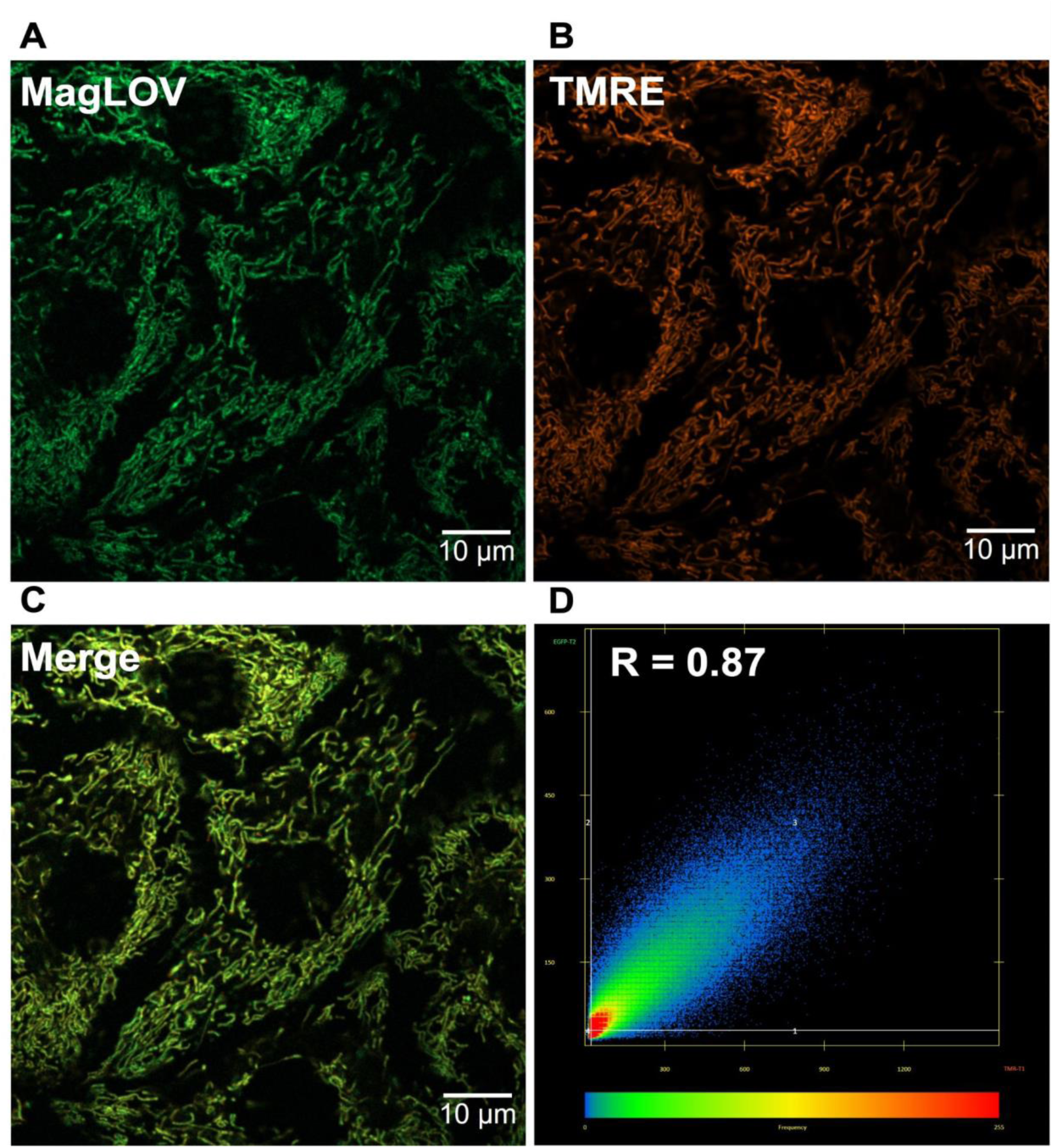
MtMagLOV2 localized to mitochondria. (A) Live-cell confocal image of mtMagLOV2 fluorescence (green channel, 488 nm excitation). (B) TMRE fluorescence (red channel, 561 nm excitation), indicating polarized mitochondria. (C) Merge of mtMagLOV2 and TMRE channels showing mitochondrial co-localization. (D) Pixel-by-pixel intensity correlation between mtMagLOV2 and TMRE signals, yielding a Pearson correlation coefficient of R = 0.87. Scale bar: 10 μm.

### MtMagLOV2 localizes to mitochondrial cristae

Using super resolution microscopy, we determined the sub-mitochondrial localization of mtMagLOV2. Using mitochondria stains TMRE (Figure 3) and Mitotracker red (Supplementary Information, <u>Fig. S4</u>), which do not overlap spectrally with the MagLOV2, we found that mtMagLOV2 localized to the cristae invaginations within the mitochondria (Figure 3 and Supplementary Information). TMRE is localized on the matrix side of the cristae invaginations, not free in the matrix ^14^. Imaging TMRE and mtMagLOV2 in a single mitochondrion (Figure 3A-C) the fluorescence profile (Figure 3D) clearly shows the mtMagLOV2 fluorescence is localized in the same region as the TMRE within the resolution of our super resolution microscope (90 nm in xy, 300 nm in z). Thus, our experimental evidence suggests the mtMagLOV2 is not freely diffusible in the matrix but is bound to the cristae invaginations. Although we cannot resolve whether the mtMagLOV2 is on the matrix or intermembrane space side (in the cristae lumen), we believe mtMagLOV2 is on the matrix side, since the TIM/TOM mechanism requires translocation into the matrix for mitochondrial localization to function properly (Figure 3E). This places the mtMagLOV2 proteins in the vicinity of the redox centers including complex I of the electron transport chain.

**Figure 3.**
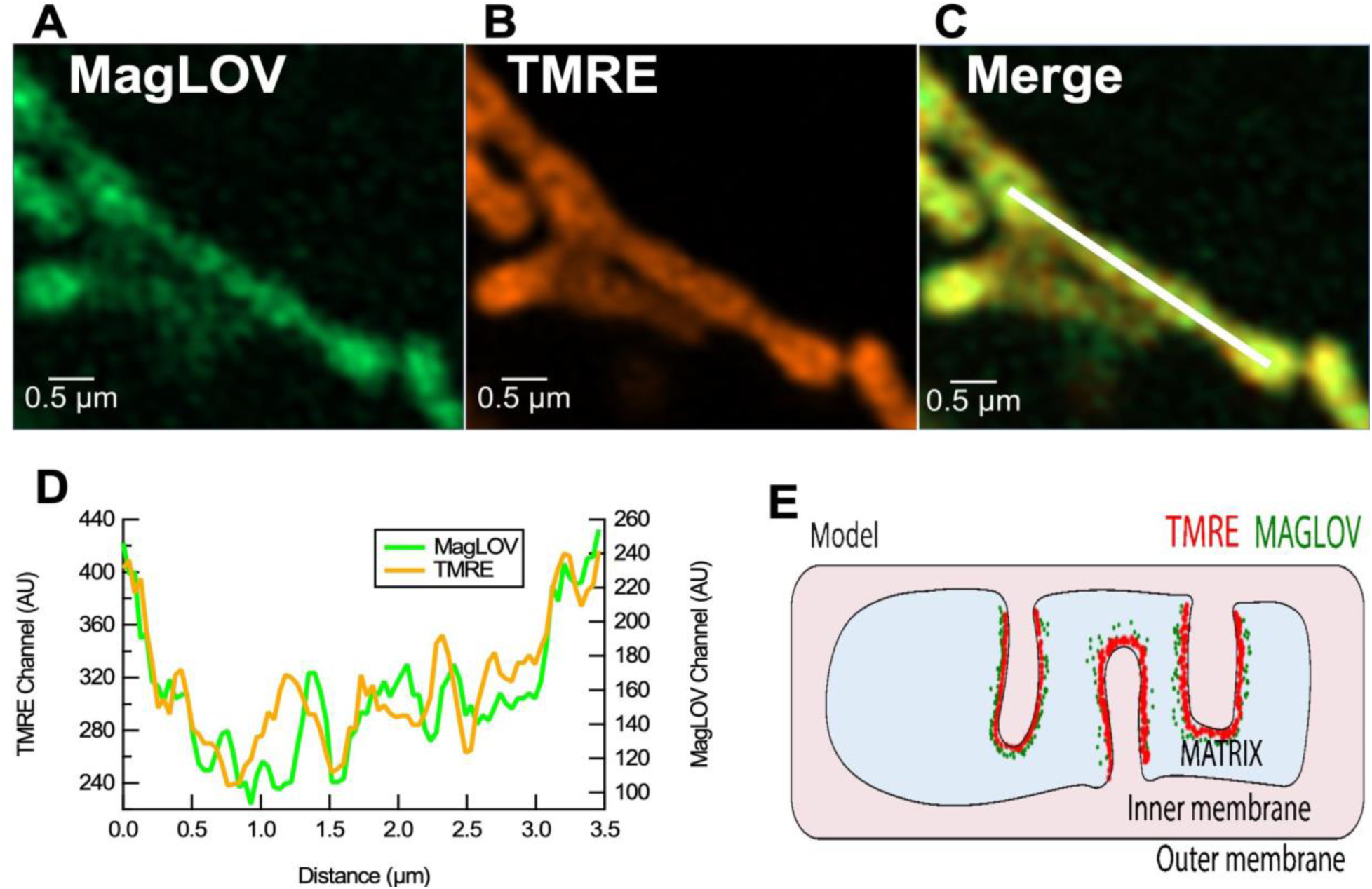
MtMagLOV2 localizes to mitochondria cristae. (A) mtMagLOV2 fluorescence (B) TMRE fluorescence (C) Merge with line-scan position (D) Fluorescence line profiles showing spatial coincidence of mtMagLOV2 and TMRE intensity peaks (E) Cartoon representation of localization to the cristae. The mtMagLOV2 is presumed to be in the matrix. Note this is not enough to resolve individual lipid bilayers of the cristae, or individual cristae if there are multiple cristae within 90 nm.

### MtMagLOV2 displays magnetic field sensitive fluorescence with confocal microscopy at the single mitochondrial level

In order to ascertain if the mtMagLOV2 fluorescence changes with magnetic field and thus can function as a biological qubit inside mitochondria, we monitored changes in mtMagLOV2 fluorescence ΔF/F when exposed to a changing magnetic field (alternating/chopped on/off with a time constant of order 11 seconds). Since the field was chopped on/off, any background drift in the fluorescence, instrumentation, focus, or even any magnetic fields generated by the mitochondria themselves cancel out, allowing us to focus purely on the mtMagLOV2 response to externally applied magnetic fields. All magnetic-field–dependent fluorescence measurements were performed prior to bleaching (Supplementary Information). Consistent with previous results *in vitro* ^1,2 2^ and in MagLOV2 expressed in Escherichia coli ^3^, we observed a magnetic field dependent fluorescence of approximately 10% in a 10 mT field (Figure 4). (The slow decay of the fluorescence is believed to be due to photobleaching.) This was true both for an entire field of view (Figure 4D), averaging over many cells, and for a single mitochondrion (field of view/region of interest, ROI) (Figure 4C). Thus, mtMagLOV2 is fully functional within individual mitochondria.

**Figure 4.**
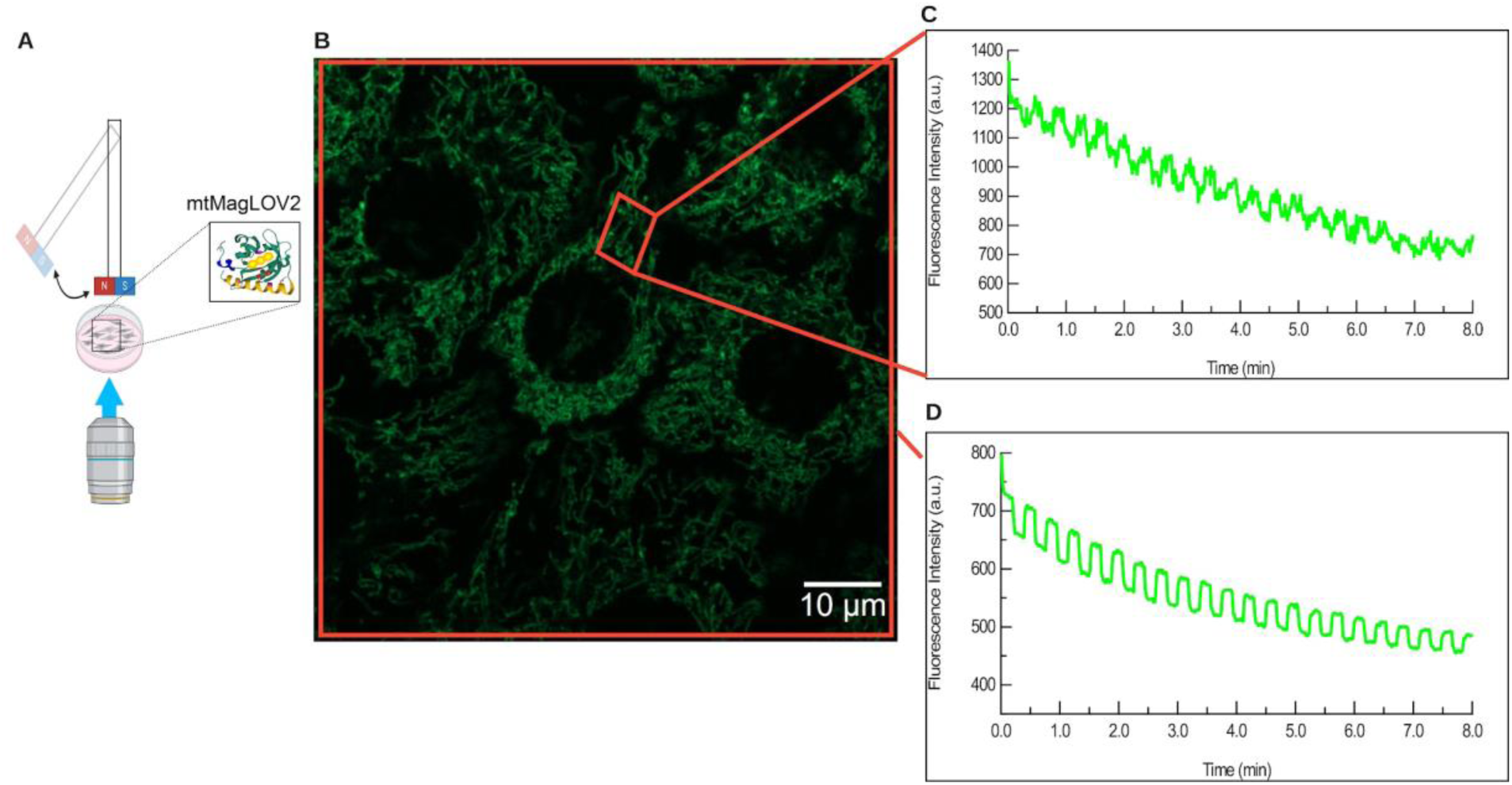
Magnetic-field–dependent fluorescence of mtMagLOV2 in live HeLa cells. (A) Schematic of the confocal imaging setup with an ON/OFF modulated magnetic field generated by a servo-positioned magnet. (B) Confocal fluorescence image of mtMagLOV2 in mitochondria. (C) Magnetic-field–dependent fluorescence trace from a small ROI corresponding to a single mitochondrion. (D) Magnetic-field–dependent fluorescence averaged over the entire field of view.

### MtMagLOV2 can serve as a quantum redox sensor of mitochondrial bioenergetic state in cell lines and cardiomyocytes

To confirm that mtMagLOV2 is sensing the bioenergetic status of the mitochondrial OXPHOS, we treated mtMagLOV2 containing Hela cells with pharmacological agents known to inhibit specific OXPHOS functions. Simultaneously, we monitored the effects on mtMagLOV2 fluorescence response in relation to an alternating (chopped) magnetic field. During treatment, we observed two effects (Figure 5): First, change in the overall fluorescence of mtMagLOV2 after treatment with the pharmacological agents. Second, a change in the response of MtMagLOV2 fluorescence (ΔF/F) to an alternating (chopped) magnetic field (on/off/on with a period of 11 seconds). We first discuss qualitative behavior, then provide quantitative analysis next.

**Figure 5.**
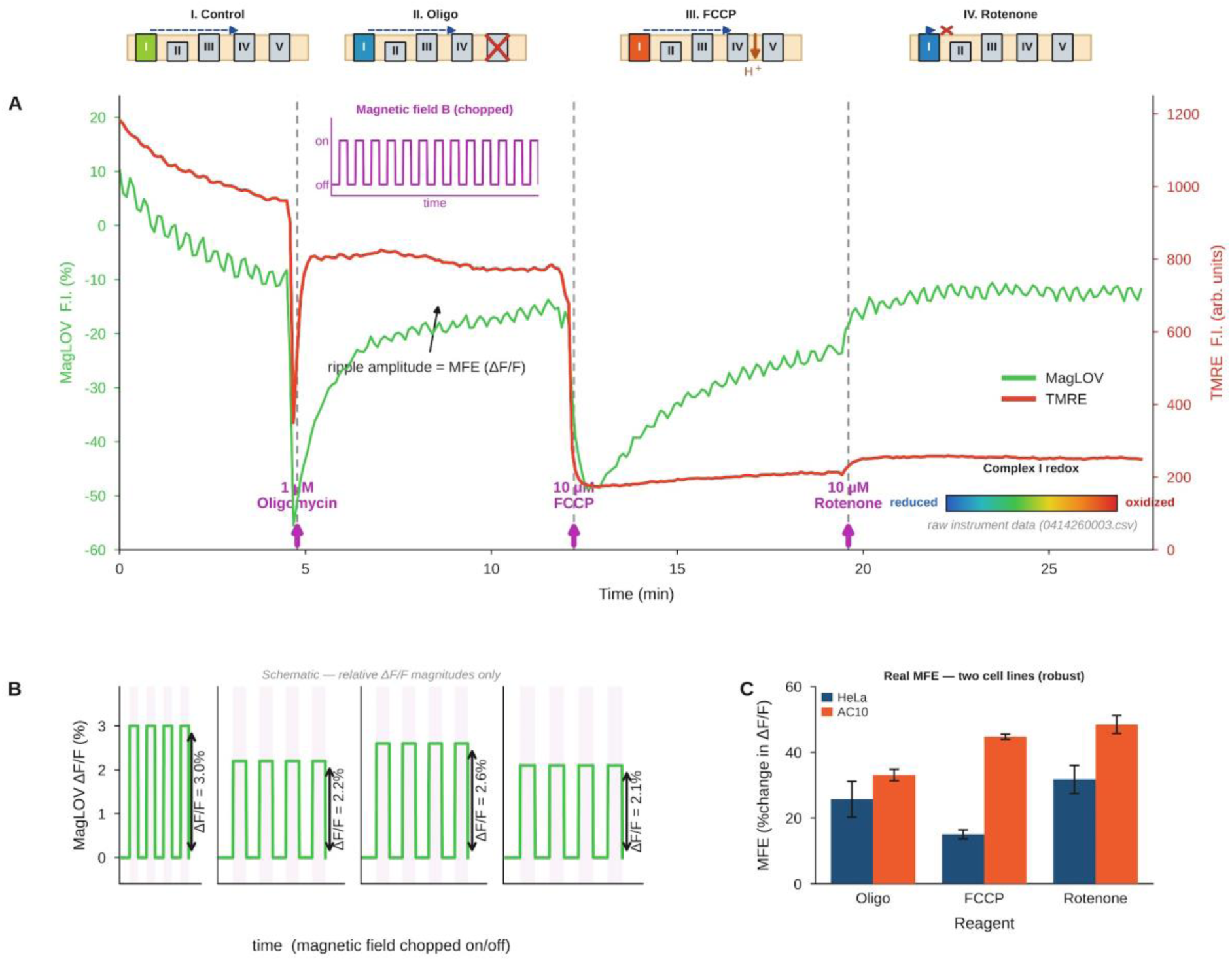
Quantum sensing of the metabolic state of mitochondria. (A) Upon pharmacological manipulation of the mitochondrial electron transport chain, the redox status of the complexes changes, including complex I. TMRE fluorescence intensity measures the mitochondrial membrane potential, providing evidence the mitochondrial are vital during the assay. Changes in TMRE membrane potential under addition of oligomycin, FCCP, and rotenone confirm the expected change in mitochondrial bioenergetic status ^23^. Simultaneously, the mtMagLOV2 fluorescence is monitored under the conditions of an alternating (chopped), externally applied magnetic field, and demonstrates clear response to the pharmacological treatment regimen, i.e. mitochondrial bioenergetic state. (B) Mathematical analysis of the change in fluorescence with magnetic field shows a unique signature for each phase of the experiment, indicated schematically as ΔF/F. (C) Predominantly glycolytic HeLa cells display a different magnetic field response (defined as the change in ΔF/F from baseline) vs. the more oxidative AC10 cardiomyocytes, demonstrating quantum bioenergetic profiling. (Error bars from n=3 independent experiments each.) Similar results are observed with both epi fluorescence (Supplementary Information) and confocal microscopy (this figure).

Changes in the mtMagLOV2 magnetic field responding fluorescence of HeLa cells, when exposed to oligomycin which inhibits the mitochondrial H^+^ transporting ATP synthase (complex V) resulting in blocked ATP synthesis and maximizes membrane potential, revealed a strong initial reduction in mtMagLOV2 fluorescence and extinction of its response to the alternating (chopped) magnetic field, though with time the fluorescence response of mtMagLOV2 recovered (Figure 5A). The recovered ΔF/F was different after treatment than before. By contrast, treatment of the mtMagLOV2 containing Hela cells with the mitochondrial uncoupled FCCP (carbonyl cyanide-p-trifluoromethoxy phenylhydrazone) collapsed the mitochondrial inner membrane potential and also initially extinguished the mtMagLOV2 fluorescence and dampened its fluorescence to the alternating (chopped) magnetic field, which again slowly recovered (Figure 5A). Finally, treatment of the mtMagLOV2-Hela cells with the complex I inhibitor, rotenone, which binds to the CoQ site downstream from the NADH-interacting FMN and the iron-sulfur cascade, had little effect on the overall mtMagLOV2 fluorescence or on the alternating (chopped) magnetic field response of mtMagLOV2 (Figure 5A). Thus, stabilizing the mitochondrial inner membrane potential, either by maximizing the potential (oligomycin) or eliminating the potential (FCCP) had a dramatic inhibitory effect on mtMagLOV2 function and magnetic field response. Since inhibiting OXPHOS (oligomycin) would fully reduce complex I FMN, while uncoupling OXPHOS (FCCP) would fully oxidize complex I FMN, the change in fluorescence of mtMagLOV2 immediately after treatment is a (quantum) sensor of the redox state of the FMN in complex I. The fact that the initial inhibitory response decayed could reflect damage to the mitochondrial inner membrane, reducing the maximum oligomycin induced membrane potential and the inhibition of the FCCP-induced maximized electron flux of the electron transport chain. These results were recapitulated in an independent laboratory at a different institution using elimination microscopy (Supplementary Information) and on cardiomyocytes (Figure 5C). The striking absence of a rotenone effect on the immediate mtMagLOV2 fluorescent response to the oscillating magnetic field localizes the interaction of mtMagLOV2 to complex I in the vicinity of FMN or the iron-sulfur centers.

While the initial transient response to pharmacological manipulation was clear, detailed analysis of the response ΔF/F to an alternating (chopped) magnetic field after stabilization reveals additional, quantitative, reproducible dependence on the bioenergetic state of the mitochondria, in particular the redox state of complex I. As ΔF/F is robust to drift, bleaching, noise, and artifacts ^22 23^, it can provide a more powerful analytical tool, as we discuss next. As Figure 5B shows, the ΔF/F after stabilization of the signal shows a dependence on the redox status of complex I: After FCCP, complex I is the most oxidized, where the largest treated ΔF/F is observed. In contrast, after oligomycin and rotenone treatment, complex I is the most reduced, where the smallest treated ΔF/F is observed. Thus, experimentally, the stabilized ΔF/F is a sensor of complex I redox status.

Both the predominantly glycolytic HeLa cells and the more oxidative AC10 cardiomyocytes responded to the alternating (chopped) field alteration induced mtMagLOV2 changes in fluorescence, when the mitochondrial were perturbed by OXPHOS active drugs. However, the size of the response of the two types of cells differed (Figure 5C). Hence, different cells will have different bioenergetic interactions with mtMagLOV2, which we will designate “quantum bioenergetic profiling. Thus, mtMagLOV2 can serve as a quantum redox sensor of mitochondrial bioenergetic state in cell lines and cardiomyocytes.

## Discussion

### Mechanism

The data clearly shows the magneto response ΔF/F is the radical pair of the MtMagLOV2/FMN cofactor, which is a standalone quantum bit, but whose magneto fluorescence depends on the redox state of nearby redox species, including complex I. This is supported by our experimental finding that ΔF/F changes with complex I redox state. complex I itself contains an FMN as the initial redox state of the electron transport chain, reduced by NADH, and therefore is the most likely pair to couple to mtMagLOV2. However, we cannot rule out the participation of other redox pairs in the sensing mechanism. The pharmacological treatments affect the redox state of the milieu of redox pairs (some of which are bound to ETC complexes, some soluble in the matrix) that make up the “redoxome” of mitochondria. Therefore, it is possible that additional redox pairs are contributing to the quantum sensing properties of mtMagLOV2. Additional genetic engineering could provide targeted redox sensors with specificity to distinct redox pairs. The overall interpretation of the data is outlined in Figure 6, showing schematically the transfection, localization, mitochondrial bioenergetic status, experimental probes and readout, and quantum state diagram. These data suggest that mtMagLOV2 is a quantum interface with mitochondrial bioenergetics.

**Figure 6.**
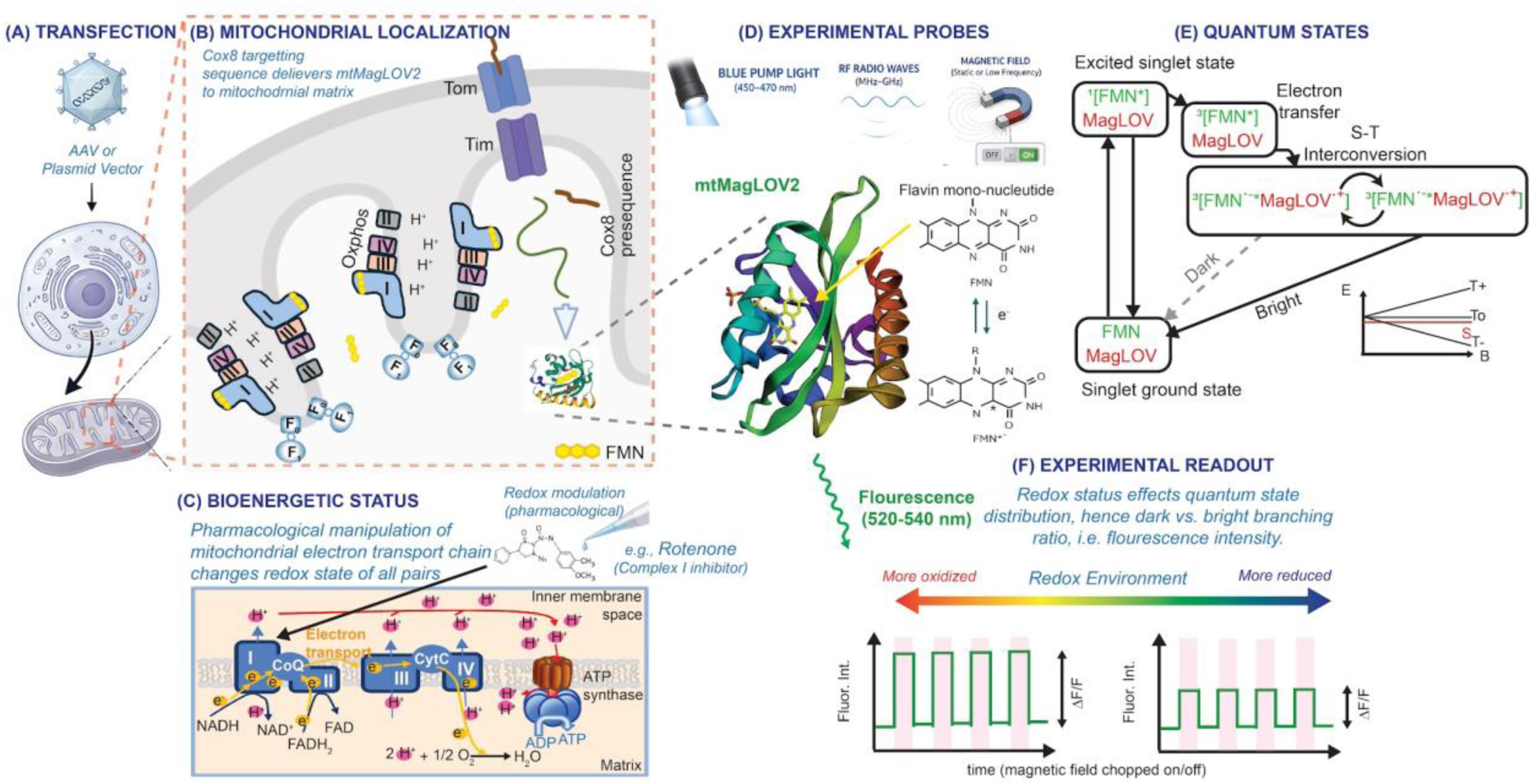
Integrated schematic of mtMagLOV2 mitochondrial targeting and quantum sensing of mitochondrial redox state. (A) Transfection: plasmid-vector transfection introduces Cox8-targeted mtMagLOV2 into cells for mitochondrial expression. (B) Mitochondrial localization: Cox8–mtMagLOV2 is imported through the TOM/TIM translocase complexes; once in the matrix, the Cox8 targeting sequence is cleaved and mtMagLOV2 folds into its mature FMN-bound form. The major components of the mitochondrial electron-transport chain (Complexes I–IV) and ATP synthase are also depicted. (C) Bioenergetic status: The redox status of the complexes of the electron transport chain (ETC) in mitochondria can be pharmacologically manipulated. (D) Experimental probes: The structure of mtMagLOV2 is not known, so the closely related structure of AsLOV (PDB structure 2V1A) is shown, with the non-covalently bound flavin mono-nucleotide (FMN). Blue-light excitation, RF/radio-wave stimulation, and static magnetic-fields are applied for quantum state manipulation. (E) Quantum states: FMN is photo-excited by blue light, then quickly transitions to a triplet state. Electron transfer from an amino acid on mtMagLOV2 reduces the flavin, creating an entangled (radical) pair. The pair, initially in the triplet state, evolves between the triplet and singlet state until it recombines, resulting in emission of a green photon, or undergoes non-radiative processes. The singlet-triplet energy level splitting depends on the static magnetic field. The transition probabilities between the various possible quantum states, and therefore the green fluorescence, also depend on the static magnetic fields. (F) Changes in the redox environment affect the quantum state distribution, resulting in a shift of the dark versus bright branching ratio of mtMagLOV2 fluorescence, producing measurable changes in ΔF/F during magnetic-field on/off cycles.

### mtMagLOV vs endogenous radical pair mechanism in mitochondria

Integrating biological quantum sensors into the electron transport chain provides new opportunities for interrogating endogenous quantum behavior of mitochondria, such as the radical pair mechanism of flavin redox reactions, magnetic field dependent respiration and reactive oxide species generation and signaling, and their downstream biological and clinical consequences. Since bioenergetics is critical for life, successful quantum state interrogation may open new vistas in engineering quantum sensors and even actuators in living systems.

While so far, we have discussed radical pairs in mtMagLOV2 only, wild-type mitochondria have their own radical pairs, independent of mtMagLOV2. Over ten years ago one of us ^24^ demonstrated magnetic field dependent respiration in mitochondria, a finding that others of us have recently reproduced and examined in more detail ^25^. Others have shown that the endogenous fluorescence of FMN is magnetic field sensitive ^26^. Taken collectively, researchers have interpreted these observations as evidence that flavins in the electron transport chain undergo a radical pair mechanism reaction. This opens the door to sensing and actuation of endogenous quantum behavior of mitochondria, for downstream biochemical and, ultimately, clinical applications. The development of mtMagLOV2 is an important tool in the toolbox of interrogation of quantum effects in mitochondrial bioenergetics.

### Time dependent quantum state populations

Previously, studies on MagLOV2 magnetic field sensitive fluorescence have used epi-fluorescence microscopy which has limited spatial resolution. We have now extended these spatial studies using super resolution microscopy to show that mtMagLOV2 is localized to the mitochondrial cristae. This is extremely important since OXPHOS redox reactions occur in the cristae membranes in association with concentrated electron and proton densities.

The quantum state population of confocal measured systems is significantly different from epi fluorescence. Pre-illumination with blue light for a similar system was required to “activate” B field sensitive fluorescence^1 2^. Although the reason is not known, it is believed to be needed to “pump” the qubits from the ground state to the excited state at a population level. After the blue “activation”, the fluorescence measurement is essentially continuous: The pump and emission are constant as a function of time, with only the B field turning on and off slowly. In prior work using epi-fluorescence ^2 3^, illumination intensities were typically 1-10 W/cm^2^ for a dwell time of order 11 seconds or more, and B field sensitivity was observed well after bleaching. In contrast, in our work, the (instantaneous) illumination intensity of a single pixel (while the beam is there) is of order 10^4^ W/cm^2^, with a dwell time of order 1 microsecond. In our confocal experiments, typically one voxel (diffraction limited volume of the point spread function of order 1 μm^3^) is illuminated for only one microsecond, before the beam moves to the next pixel. This establishes clearly the applicability of biological quantum sensors using extremely short and high intensity illumination and fluorescence.

This demonstrates that transient, pulsed, time dependent quantum state manipulation is possible in biological qubits mtMagLOV2 localized to mitochondria. Thus, our experimental design for the mtMagLOV2 protein provides a new approach for studying the temporality of mitochondria bioenergetic quantum dynamics.

The extension of these techniques to engineered pulsed experiments for quantum state manipulation ^27 28^ could enable additional detailed understanding of the lifetimes and transition rates of the quantum states, creating a detailed map of all possible transitions. Such an “almanac” would enable more targeted redox sensing, increased sensitivity, and even the possibility to perform coherent quantum state manipulation of biological qubits coupled to classical or quantum biological processes.

### Summary

In summary, we have demonstrated mitochondrial targeted biological qubit quantum sensors with 90 nm spatial resolution and microsecond time resolution. This lays the groundwork for a new class of quantum sensors in bioenergetics and biology.

## Acknowledgments

This work was supported in part by the National Science Foundation (NSF) award #2153425, and NIH grants CA259635 and AG078814 (DW). This material is based upon work supported by the Air Force Office of Scientific Research under award numbers FA9550-23-1-0436 and FA9550-23-1-0061. Any opinions, findings, conclusions, or recommendations expressed in this material are those of the author(s) and do not necessarily reflect the views of the United States Air Force. We thank Prof. Harrison Steel (Oxford) for useful discussion and providing a preliminary vector for in vitro testing of MagLOV2 (in progress). We thank Andrew York and Maria del Mar Ingaramo for interesting discussions and suggestions for vector design to target mitochondria.

## Author contributions

PA, SN, NS, MJ, FG: fluorescence experiments and cell culture. PC, YC: Transfection and vector design and implementation. PH, RU, DW: Quantum and biology interpretations. PB: Conceptualization and ideation, project supervision. All authors: Writing and revising manuscript.

## Data and code availability

The raw imaging data and MATLAB analysis code supporting the findings of this study have been deposited in Zenodo at [[Confocal Data](https://zenodo.org/records/20978740?preview=1&token=eyJhbGciOiJIUzUxMiJ9.eyJpZCI6Ijg2OTk0YmMxLWQyOGItNGQxYS1iNzg4LTk2YzU5YjQzODYxNSIsImRhdGEiOnt9LCJyYW5kb20iOiIzNjE3MDU1ZTliMDEwM2MyYzUyMjc2ZjdhNTUzNGM5MiJ9.R_FXM0B3dyKZEWmSASiNiKeAqDoOjMznnT6CvL77u63kxzgh3SfeV_W5qBxzhWAgRNffWQvF4z8XLG76GlnewQ), [Epi Data](https://zenodo.org/records/20940183?preview=1&token=eyJhbGciOiJIUzUxMiJ9.eyJpZCI6ImJlODE0MmU3LThkMDQtNDlmNC1iY2I1LWMxMjg1NTI5MDVkZCIsImRhdGEiOnt9LCJyYW5kb20iOiJmZDliN2U2ZDI5OWQ4ZjYwMTVjODNjMjRkN2M2NTlkNSJ9.sdqkRThQG19jDf6Bci64dw1gHJUeShX8Z-02C06-dfmLc-Hd35osHVvrukizFs4xFbhdNETVSR3CsIzGIAxSWg)]. The dataset will be made publicly available upon publication. The MATLAB code is also provided in the Supplementary Information.

Supplementary Information is available for this paper.

## Materials and Methods

### Cell culture and maintenance

HeLa cells (ATCC CCL-2) were cultured in high glucose Dulbecco’s Modified Eagle Medium (DMEM) supplemented with 10% (v/v) fetal bovine serum (FBS) and 1% (v/v) penicillin–streptomycin. AC10 cardiomyocytes (ATCC CRL-3569) were cultured in DMEM/F-12 medium supplemented with 12.5% (v/v) fetal bovine serum (FBS) and 1% (v/v) penicillin–streptomycin. All Cultures were maintained at 37 °C in a humidified incubator containing 5% CO₂. Cells were routinely monitored by brightfield microscopy to confirm normal morphology and free of contamination. For imaging experiments, cells at approximately 3 × 10⁴ cells/cm² were seeded onto 35 mm glass-bottom dishes (No. 1.5, MatTek). After seeding, cells were allowed to adhere and recover for 24–48 hours before treatment or staining. Cells were passaged every two days at 70–80% confluency using 0.05% trypsin–EDTA, and only early-passage cells with healthy morphology were used for imaging.

### Mitochondrial membrane potential assay and mtMagLOV2 colocalization (confocal)

For mitochondrial staining, HeLa–mtMagLOV2 and HeLa Wild Type cells were incubated with tetramethylrhodamine ethyl ester (TMRE, Invitrogen) at a final concentration of 10 nM in complete growth medium for 15 minutes at 37 °C. AC10 cardiomyocytes cells were stained under the same conditions except that the incubation time was extended to 30 minutes. After staining, cells were washed twice with warm PBS and immediately imaged in fresh phenol-red-free DMEM to minimize background fluorescence.

For colocalization analysis, mtMagLOV2 fluorescence was recorded under 488 nm excitation, and TMRE was excited at 561 nm with emissions collected using separate detection windows to prevent bleed-through. Colocalization between the mtMagLOV2 and TMRE signals was quantified using the Coloc2 plugin in Zeiss software.

During time-lapse recording, FCCP (10 µM) was added directly to the imaging medium to depolarize mitochondria. Imaging continued immediately after addition to capture the progressive loss of TMRE fluorescence and the corresponding changes in mtMagLOV2 signal distribution. The rapid decline of TMRE intensity confirmed efficient mitochondrial depolarization under these conditions.

To monitor the dynamics of the mtMagLOV2 signal, time-lapse confocal imaging was performed using a 63× oil-immersion objective (NA 1.4). Images were acquired sequentially at fixed intervals to capture changes in fluorescence intensity over time in response to magnetic field modulation.

### Microscopy and magnetic field setup (confocal)

A Zeiss LSM 900 confocal microscope equipped with an Airyscan detector was used for fluorescence imaging. All experiments used a 63× oil-immersion objective with a numerical aperture (NA) of 1.4. mtMagLOV2 fluorescence was excited using a 488 nm laser at 1% laser power, and emission was collected between 500 and 550 nm. TMRE was excited at 561 nm (0.3% laser power), with emission detected between 565 and 630 nm. The photomultiplier gain was set to 1000 V for the 488 nm channel and 700 V for the 561 nm channel. All acquisition settings were kept the same throughout each experiment.

We recorded time-lapse sequences with a frame time of 1.67 seconds and an image size of 917 by 917 pixels, without signal averaging. Recording continued for 30 minutes with no intervals between frames, allowing for high temporal resolution in monitoring fluorescence changes. The imaging conditions were optimized to minimize photobleaching while maintaining a strong signal.

Magnetic field stimulation was achieved using a custom-built servo-controlled system designed to modulate a small permanent magnet near the imaging plane. To ensure stable alignment during imaging, we fabricated a 3D-printed mounting frame that securely fixed the servo motor close to the microscope stage. We connected the motor arm to a 3D-printed hex-shaped rod (Figure 4A) that held a neodymium magnet at its distal end. At the position of the cells, the magnetic field amplitude measured approximately 10 mT (±0.2 mT), as assessed with a calibrated Gaussmeter using a Hall probe sensor.

Field modulation was controlled by an Arduino-based system programmed to alternate the magnetic field in precise ON/OFF cycles. From minutes 0 to 6, the magnet was toggled every 11 seconds (11 seconds ON / 11 seconds OFF). To ensure that variations in fluorescence were dependent on the magnetic field and not due to mechanical or optical artifacts, the servo was turned off for 2 minutes, during which no periodic modulation occurred. After this pause, the field cycling resumed for an additional 7 minutes with longer intervals of 22 seconds (22 seconds ON / 22 seconds OFF). This setup enabled real-time recording of mtMagLOV2 fluorescence dynamics under controlled magnetic field modulation.

### Image analysis and statistics (confocal)

For quantitative comparison of wild type HeLa cells to mtMagLOV2 fluorescence, image processing and fluorescence quantification were performed using ImageJ (Fiji, version 2.3). Background fluorescence was subtracted from each frame using a rolling-ball radius of 50 pixels, and the mean fluorescence intensity was measured within manually defined regions of interest (ROIs) corresponding to individual cells or mitochondrial clusters.

To better quantify the relative fluorescence strengths of mtMagLOV2 and wild type HeLa cells, we performed two complementary analyses: an initial whole-image comparison and a more rigorous cell-by-cell ROI-based measurement.

In the whole-field analysis, the mean pixel intensity was calculated over the entire image area, which includes not only mitochondria but also large regions of cytosol, nucleus, and background. Because wild type HeLa cells exhibit diffuse, low-level autofluorescence throughout the cytoplasm, this global averaging underestimates the true contrast between mtMagLOV2-positive mitochondria and the weak mitochondrial autofluorescence in wild type cells. Using this coarse metric, mtMagLOV2 appeared only slightly brighter than wild type HeLa cells.

To obtain a more accurate and biologically meaningful estimate, we next performed a targeted ROI-based analysis on individual cells. For each image, multiple freehand ROIs were drawn directly over mitochondrial regions displaying true signal (excluding nuclei and avoiding saturated regions). For every cell, the mean fluorescence intensity within each ROI was extracted. Importantly, to correct for imaging noise and camera offset, a separate background ROI was selected in an area devoid of cellular structures, and this value was subtracted from each measurement. This background-corrected intensity provides a more faithful representation of the actual protein-derived fluorescence.

When the analysis was restricted to mitochondrial ROIs and corrected for background, the contrast between mtMagLOV2 and wild type fluorescence increased substantially: across six cells per condition, the average mtMagLOV2 signal was approximately 2-fold higher than the autofluorescent signal observed in wild type HeLa mitochondria. This refined estimate reflects the true molecular brightness difference, whereas the whole-image analysis is biased downward by the inclusion of large non-mitochondrial regions in wild type cells.

Colocalizatio between mtMagLOV2 and TMRE signals was analyzed using the Coloc2 plugin in the Zeiss software.

For time-lapse datasets, raw fluorescence intensity traces for the TMRE and mtMagLOV2 channels were analyzed using Igor Pro (WaveMetrics) and MATLAB. The TMRE signal was plotted as absolute fluorescence intensity in arbitrary units, while mtMagLOV2 fluorescence was normalized to the mean intensity across field-OFF frames and expressed as percentage change according to:

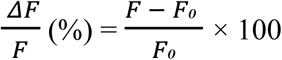

Where *F* is the fluorescence intensity at each time point, and *F₀* was defined as the average mtMagLOV2 fluorescence intensity across field-OFF frames.

The Magnetic Fluorescence Effect (MFE) was defined as the relative percent change in ΔF/F following pharmacological perturbation:

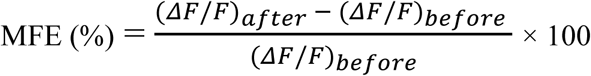

Where ((*ΔF*/*F*)*_before_*) represents the average magnetic-field-induced fluorescence modulation before drug addition, and ((*ΔF*/*F*)*_after_*) represents the modulation measured after the signal reached a new steady state following treatment. For this calculation, MATLAB code was used to define pre-treatment, post-treatment, and drug-addition transient windows manually from each time-lapse trace. The transient period immediately following drug addition was excluded from analysis to avoid injection-related artifacts and signal stabilization effects.

### Microscopy and magnetic field setup (epi)

We collected the mtMagLOV2 data on a homebuilt widefield fluorescence microscope constructed on a Thorlabs optical cage system. Excitation was provided by a Thorlabs Chrolis six-channel LED engine, excitation at 475 nm. The illumination power was measured with a Volumetric intensity meter of 4.6 mW/cm^2^. Fluorescence was collected through a Nikon 20× 0.4 NA infinity-corrected objective and detected by an Andor Zyla 4.2 sCMOS camera (2048 × 2048 pixels, 6.5 µm pixel size) mounted on an ASI FTP stage. Magnetic stimulation was delivered via a custom hand-wound Helmholtz coil, 3D-printed to fit directly onto the ASI stage adjacent to the objective producing a calibrated field (GM-2 gaussmeter, AlphaLabs) of 14 mT at the sample plane. Frame-synchronized acquisition was achieved through a hardware trigger chain: an Arduino microcontroller governed all experimental timing, synchronizing field switching with image acquisition, and issued TTL pulses to a Stanford Research Systems DG535 delay generator, which in turn triggered the Zyla for each frame. Frame period was set to the camera exposure time plus a 50 ms guard interval to ensure complete sensor readout between acquisitions. A pulsed magnetic field was applied in an “on/off” cycle with a periodicity of 5 s over a total stimulation duration of 60 s, with no pre-bleaching interval to capture the initial baseline fluorescence. Time-lapse imaging was performed at a temporal resolution of 1.8 frames per second, yielding 109 frames per experimental run.

To establish a baseline for magnetic responsiveness, untreated HeLa–mtMagLOV2 cells were imaged in pure FluoroBrite™ DMEM (Figure S13).

### Image analysis (epi)

MtMagLOV2 fluorescence time-series data were analyzed using a homebuilt Python analysis tool. Briefly, individual TIFF frames acquired by the qudi instrument control software were loaded sequentially and assembled into a stack. The magnetic field ON/OFF state of each frame was reconstructed from the acquisition timing parameters described above. Mean projection images were used for cell segmentation, which was performed using the Cellpose cyto3 deep-learning model ^29^, with cells filtered by area to exclude debris and out-of-focus objects. For each segmented cell, mean fluorescence intensity was extracted per frame. Normalized fluorescence responses (ΔF/F) were computed as (F − F₀)/F₀, where F₀ is the mean intensity across field-OFF frames, serving as a normalization to account for cell-to-cell variability in baseline expression. Frames within a two-frame guard window around each field transition were excluded from F₀ calculation and statistical comparisons to avoid contamination by transient switching artifacts. Cells were classified as responsive, inverse-responsive, or non-responsive based on the sign consistency of ΔF/F across field-ON cycles and the signal-to-noise ratio relative to field-OFF variance.

### Plasmid Construction and Lentiviral Production

The critical elements of our design are as follows: 1) to localize MagLOV2 to mitochondria, the Cox8 presequence was fused in-frame to the N-terminus of MagLOV2, thereby directing mitochondrial import via the TIM–TOM import complexes; 2) the expression cassette was incorporated into a lentiviral expression vector containing an IRES–puromycin resistance module to enable enrichment of cells retaining expression cassette. Lentiviral particles were produced in Lenti-X 293T cells and used to infect HeLa or AC10 cells. Puromycin-resistant clones were isolated and expanded for further analysis.

The plasmid pLV-EF1α-IRES-Puro was generated using pLV-EF1α-IRES-Hygro and pQCXIP as templates. Gene segments were synthesized by Integrated DNA Technologies (IDT), amplified using the primer pair LHCox8 and LHHPAR, and assembled into BamHI/HpaI-digested pLV-EF1α-IRES-Puro to create the construct designated pLPMtMagLOV2 (R10). Similarly, pLPMtMagLOV2-3Flag were created with the 3 Flag tags at the C-terminus allowing monitoring using M2 antibody. The resulting plasmid was verified by Sanger sequencing using the EF1α primer to confirm the integrity of the mtMagLOV2 or the mtMagLOV2-3Flag sequence.

Recombinant lentivirus was produced using the Lenti-X 293T cell line (Takara Bio USA, Inc.). The packaging mix included the VSV-G envelope plasmid and the second-generation packaging plasmid pCMV-dR8.91, which supplies the *gag* and *pol* proteins required for lentiviral assembly. Transfection was performed using the calcium phosphate precipitation method. At 48–72 hours post-transfection, the culture supernatant containing lentiviral particles was collected, clarified by filtration through a 0.45 µm PES membrane filter, and used to infect HeLa or AC10 cells in the presence of 4 µg/mL polybrene. Following a 24-hour incubation period, the medium was replaced with fresh growth medium supplemented with 10 µg/mL puromycin for selection. After approximately 10 days, puromycin-resistant colonies of stable integrants became visible. Clones exhibiting mitochondrial fluorescence above background autofluorescence were isolated and expanded for further analysis.

In addition to the original mtMagLOV2 (R10) construct, a C-terminal triple FLAG-tagged variant (mtMagLOV2-3F) was also generated and expressed in HeLa or AC10 cells. The 3FLAG construct was used in later experiments to confirm reproducibility of the magnetic fluorescence response and to improve signal stability during long-term culture. The mtMagLOV2-3F cells exhibited qualitatively similar magnetic-field–dependent fluorescence behavior and mitochondrial perturbation responses compared to the original mtMagLOV2 line, while providing higher fluorescence intensity than the original mtMagLOV2 construct.

### Optically detected magnetic resonance (ODMR)

HeLa cells stably expressing mtMagLOV2 were imaged in time-lapse mode using a Zeiss LSM 900 Airyscan confocal microscope (Carl Zeiss) with 488 nm excitation (1% laser power, ∼60 μW at the sample plane). MtMagLOV2 fluorescence was collected in the 500–550 nm emission range, corresponding to the intrinsic flavin (FMN) emission of the protein. Images were acquired with a frame interval of 100 ms, consisting of 70 ms acquisition and 30 ms delay ^30^.

An RF signal was generated using an Agilent N5181A signal generator and swept from 100 MHz to 1 GHz in 1 MHz increments. The signal generator was programmed to advance by 1 MHz every 100 ms, providing frame-by-frame correspondence between each acquired image and the programmed RF frequency.

The RF signal was amplified using a Mini-Circuits ZHL-10M2G0005 + amplifier and delivered to a 1 mm-diameter omega-loop antenna. The RF power after amplification was measured to be +25 dBm (316 mW) at the amplifier output. Assuming negligible cable loss and an approximately 50 Ω matched load, this corresponds to an estimated loop-center B₁ field of ∼0.14 mT peak. Because the actual RF power coupled into the loop depends on impedance matching and reflected power, this B₁ value should be regarded as a nominal estimate unless calibrated directly at the antenna or sample position.

A static magnetic field B₀ ≈ 19.6 mT was applied approximately perpendicular to B₁. This places the expected electron spin resonance condition at f_res_ = γₑB₀ ≈ 28.025 MHz mT⁻¹ × 19.6 mT ≈ 549 MHz ^3^, corresponding to t ≈ 45 s within each frequency sweep.

