## Supplementary Information for "A quantum interface with mitochondrial bioenergetics"

#### Expanded discussion of the radical pair mechanism in Figure 1

A RP born in the triplet state initializes its spin population distributed across $T_+$, $T_0$, and $T_-$. At low magnetic fields (~50 μT), hyperfine interactions drive coherent mixing between these triplet sublevels and the singlet state S, continuously converting triplet character into singlet character over the RP lifetime. The accumulated singlet character across the spin population over the RP lifetime determines the probability of ground state recovery and therefore fluorescence yield. At high magnetic field (>1 mT), Zeeman splitting energetically isolates $T_+$ and $T_-$ from S. Two thirds of the initial spin population can no longer mix into singlet character, while the $T_0$ fraction remains active. The total singlet character that develops before RP collapse is therefore reduced, and fluorescence decreases correspondingly. This framework makes no assumptions about the relative rates of subsequent chemical steps. Regardless of the downstream fate of singlet and triplet radical pairs, whether recombination, protonation, or slow redox return, the magnetic field effect is determined entirely at the level of spin population dynamics. The observed decrease in fluorescence at high field, and its recovery upon removal of the field or application of a resonant RF field, is a direct consequence of this spin population budget.

#### Mitochondrial targeted biological qubit: mtMagLOV2

To generate mtMagLOV2, we added the N-terminal mitochondrial targeting peptide from Cox8 to the R10 variant reported by Oxford ("MagLOV2") [1] (Figure S1A) such that the unfolded protein will be imported through the TOM/TIM complexes to the matrix where it will fold and bind to mitochondrial matrix FMN [2] [3] [4]. Delivery of the polypeptide to target cells by Lenti-virus transduction resulted in mtMagLOV2 localization into the mitochondrial matrix where the N-terminal targeting peptide is cleaved off and the resulting polypeptide folds into the MagLOV2 conformation (Figure S1B).

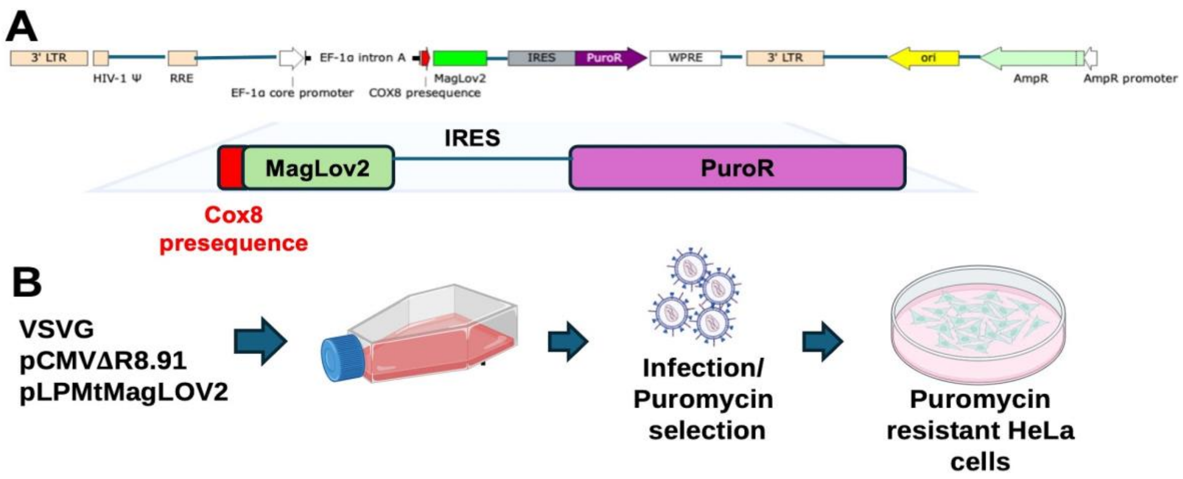


**Figure S1. Schematic of mtMagLOV2 construct design and stable HeLa cell generation.** (A) Map of the pLPMtMagLOV2 plasmid. The construct contains a bicistronic expression cassette in which the Cox8 presequence is fused in-frame to MagLOV2, followed by an IRES–puromycin resistance module. (B) Generation of mitochondria-localized mtMagLOV2 in HeLa cells. Lentiviral particles were produced in Lenti-X 293T cells using the calcium-phosphate precipitation method, harvested, filtered, and used to infect HeLa cells.

#### Confocal and Epi

MtMagLOV2 fluorescence MFEs were observed under two experimentally distinct illumination conditions: confocal microscopy with pulsed excitation at 1% laser power and a 1 μs pixel dwell time, and continuous epifluorescence illumination under steady-state conditions. Despite the fundamental difference in these measurement regimes, both yield the same qualitative MFE dependence, which we argue reflects the intrinsic single-event nature of the underlying spin physics.

In the confocal case, the 1 μs dwell time is comparable to or encompasses the RP lifetime, such that each pixel visit constitutes an essentially single-pass measurement of one cohort of transiently generated RPs. The spin dynamics occur within that experimental timeframe, and the fluorescence collected reflects the singlet character accumulated within a single RP generation. In the epifluorescence case, continuous illumination drives steady-state cycling of the FMN population through repeated RP formation and collapse events. The observed fluorescence represents a time-averaged singlet yield per RP generation.

The consistent MFE observation across both instruments provides important mechanistic insight, demonstrating that the MFE is determined at the level of the individual RP events, not through cumulative ensemble effects requiring many photocycle iterations. In both cases, the applied magnetic field controls the same quantity: the fraction of the initial triplet spin population accessible for singlet-triplet mixing over the RP lifetime. At high field, Zeeman splitting isolates $T_+$ and $T_-$, removing two thirds of the initial spin population from the mixing manifold and reducing the total singlet character that develops before RP collapse. This reduction in accumulated singlet character directly reduces fluorescence yield, independent of the illumination modality or the specific rate constants governing subsequent chemical steps. The convergence of pulsed and continuous measurements on the same MFE therefore constitutes evidence that the spin population budget, established at RP formation and governed by the applied field, is the primary and sufficient determinant of the observed magnetic field response.

A notable distinction between the two regimes is the relationship of the measurement to the initial photobleaching transient. In the confocal case, the brief 1 μs dwell time samples the FMN population prior to any significant photobleaching, capturing the MFE from the fully intact ground state population. In the epifluorescence case, continuous illumination drives an initial photobleaching transient over the first several seconds, after which the measurement is made on the resulting photostable steady-state population. The observation of consistent MFE character across both conditions, before and after this transient photobleaching process, further supports the conclusion that the MFE is an intrinsic property of the spin dynamics at the individual RP level, rather than an artifact of population heterogeneity or photodegradation state.

**mtMagLOV2 colocalization with MitoTracker Red**

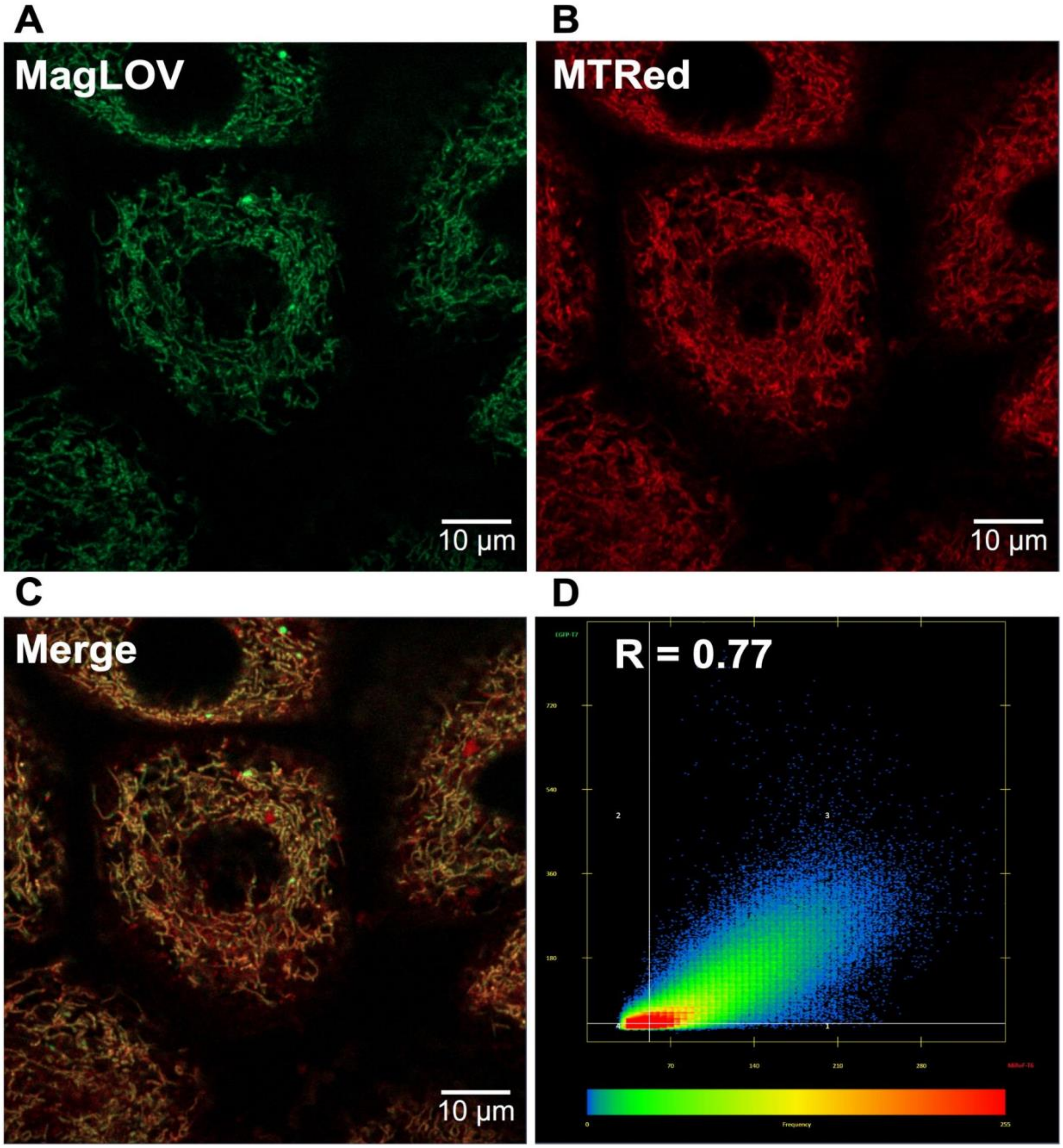


**Figure S2. MtMagLOV2 colocalization with MitoTracker Red in live mtMagLOV2 HeLa cells.** (A) Live-cell confocal image of mtMagLOV2 fluorescence (green channel, 488 nm excitation). (B) MitoTracker Red fluorescence (red channel, 561 nm excitation), labeling the mitochondrial network independently of mitochondrial membrane potential. (C) Merge of mtMagLOV2 and MitoTracker Red channels showing strong spatial overlap along mitochondrial structures. (D) Pixel-by-pixel intensity correlation analysis between mtMagLOV2 and MitoTracker Red signals, yielding a Pearson correlation coefficient of $R = 0.77$. Scale bar: 10 μm.

#### Dyes and spectra

To enable simultaneous monitoring of mitochondrial membrane potential (TMRE, red channel) and MagLOV2 fluorescence (green channel), we selected dyes with minimal spectral overlap. As shown in Figure S3, the excitation (488 nm) and emission (500–550 nm) window used for MagLOV2 imaging remains spectrally distinct from TMRE excitation (561 nm) and emission (565–630 nm). This separation ensures that the observed magnetic-field–dependent changes in MagLOV2 fluorescence cannot arise from TMRE bleed-through or detector cross-talk.

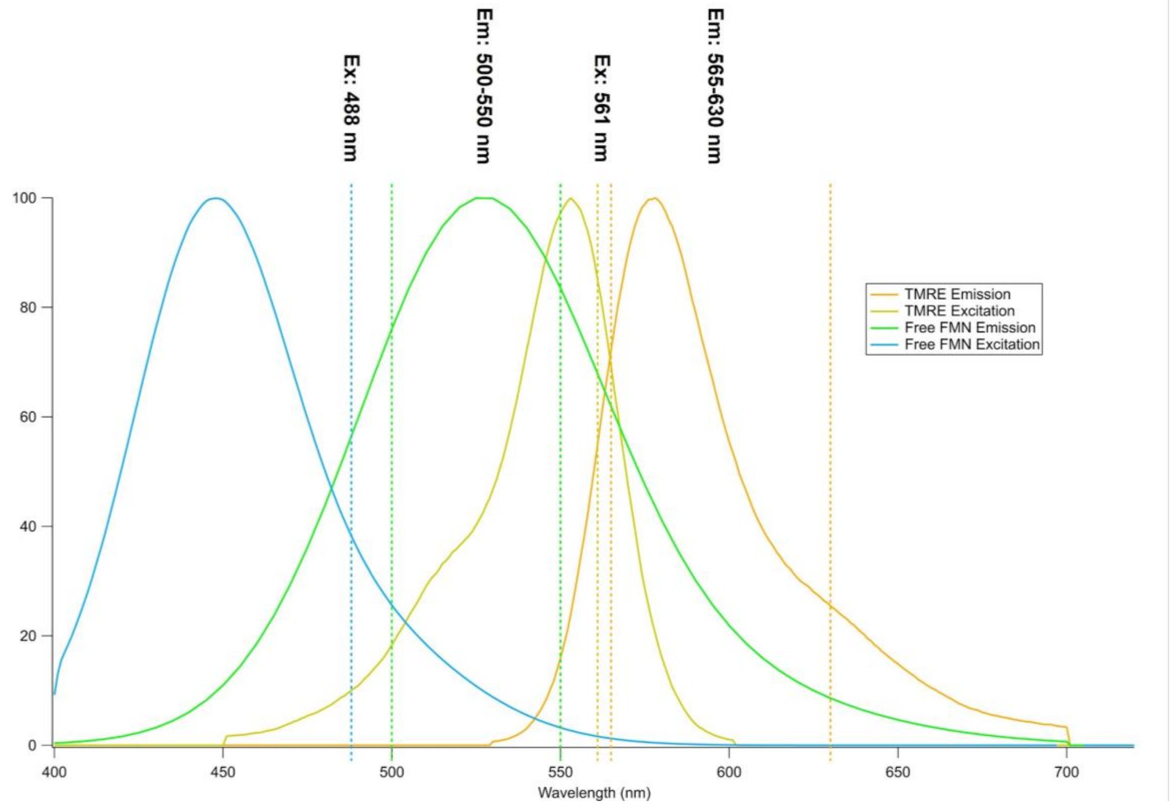


**Figure S3. Excitation and emission spectra of Free FMN and TMRE.** TMRE shows a peak excitation near ~550 nm and emission around ~575 nm, consistent with its use under 561 nm excitation and red-channel detection. Free FMN absorbs in the blue region compatible with 488 nm excitation and emits in the green range, supporting detection in the 500–550 nm window used for LOV/FMN-based MagLOV2 fluorescence imaging. FMN spectra were adapted from [5] [6]; TMRE spectra were adapted from the manufacturer/database.

#### Green mitochondrial fluorescence in mtMagLOV2-expressing and wild type cells

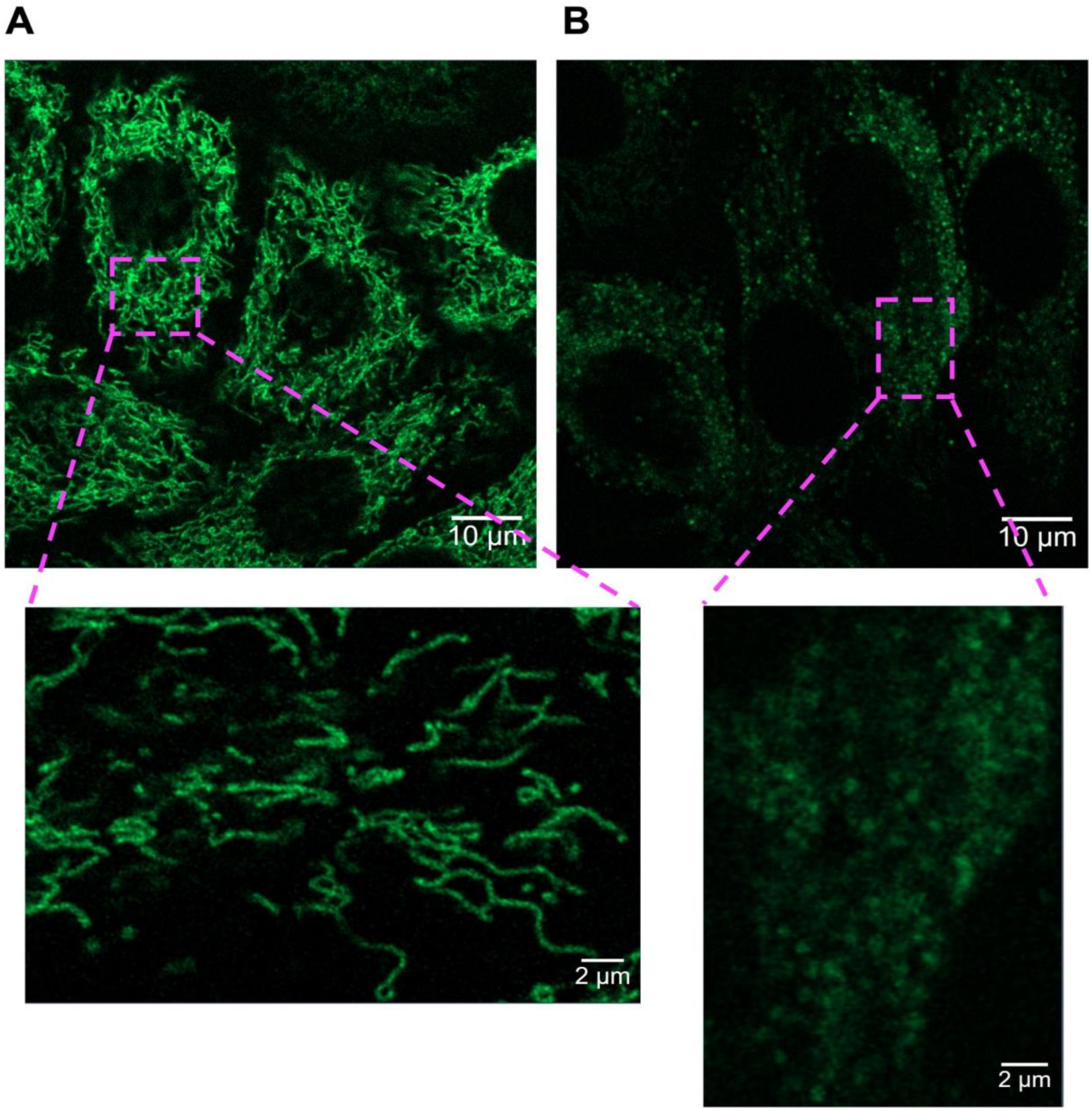


**Figure S4. Green mitochondrial fluorescence in mtMagLOV2-expressing and wild type cells.** (A) HeLa cells expressing mtMagLOV2. (B) Wild type HeLa cells showing endogenous green autofluorescence. The mean green fluorescence intensity in mtMagLOV2-expressing cells is ~2-fold higher than the wild type endogenous fluorescence.

#### Control Experiments for background and artifact detection

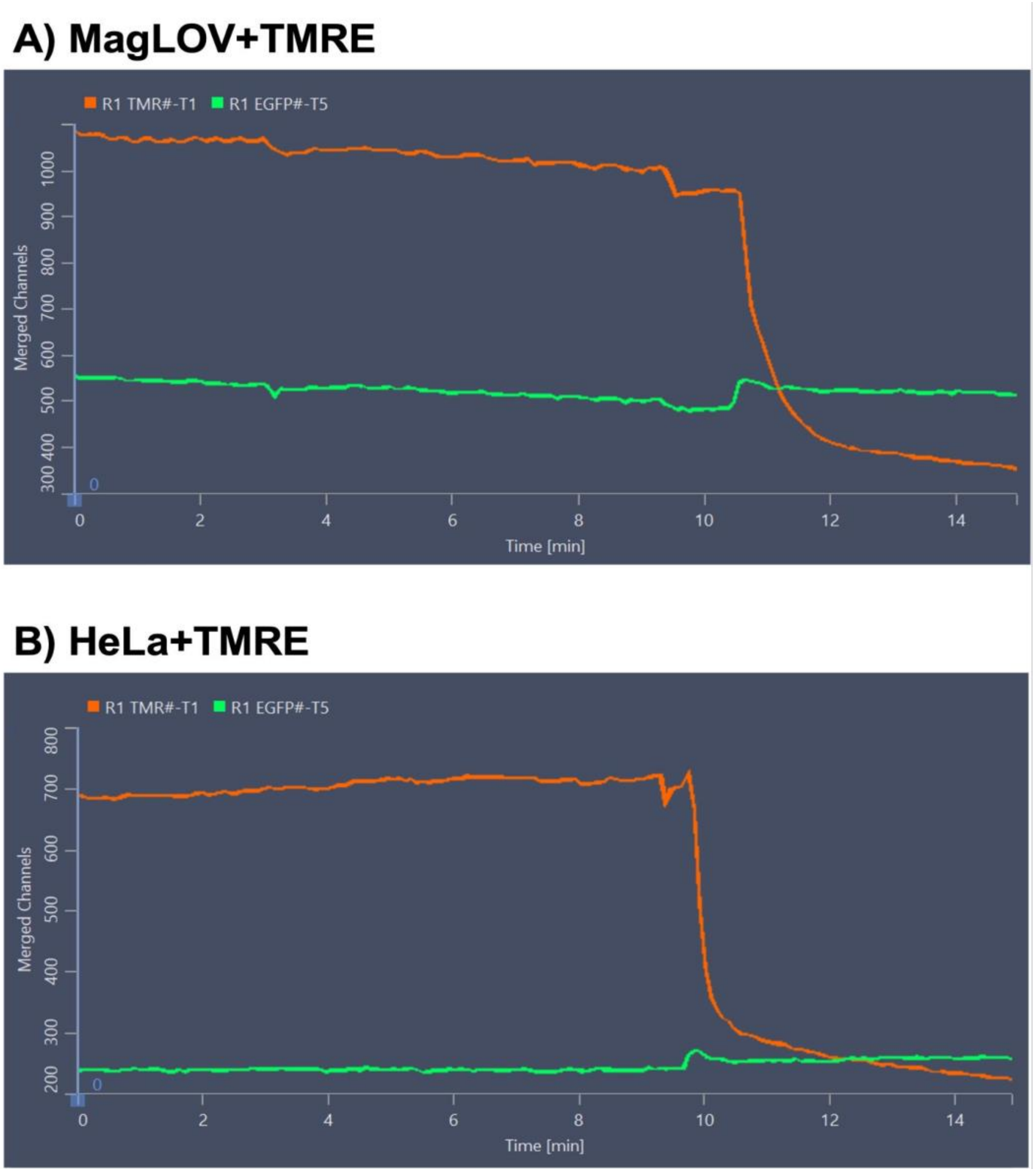


**Figure S5. Mechanical control experiment. As a mechanical control, the servo motor was operated with the same motion profile while the magnet was removed.** Under (A) mtMagLOV2+TMRE and (B) HeLa+TMRE, no periodic modulation of mtMagLOV2 fluorescence was observed, confirming that the detected oscillations were not caused by vibrations or stage instability.

To ensure that the magnetic field modulation did not introduce optical or mechanical artifacts in the imaging system, control experiments were performed in the absence of cells. Solutions containing either PBS or cell culture medium (DMEM + 10% FBS), with and without TMRE (10 nM), were imaged under the same confocal conditions used for live-cell experiments.

Time-lapse recordings were acquired for 10–15 minutes at a frame time of 1.67 s using 1% (488 nm) and 0.3% (561 nm) laser powers. Magnetic field modulation was applied in 11-second ON/OFF cycles, followed by FCCP addition as a procedural control.

No periodic or correlated changes in fluorescence were detected under any of these conditions, confirming that the observed magnetic-field-dependent oscillations in live-cell experiments were not due to optical noise, dye instability, or mechanical vibration of the setup.

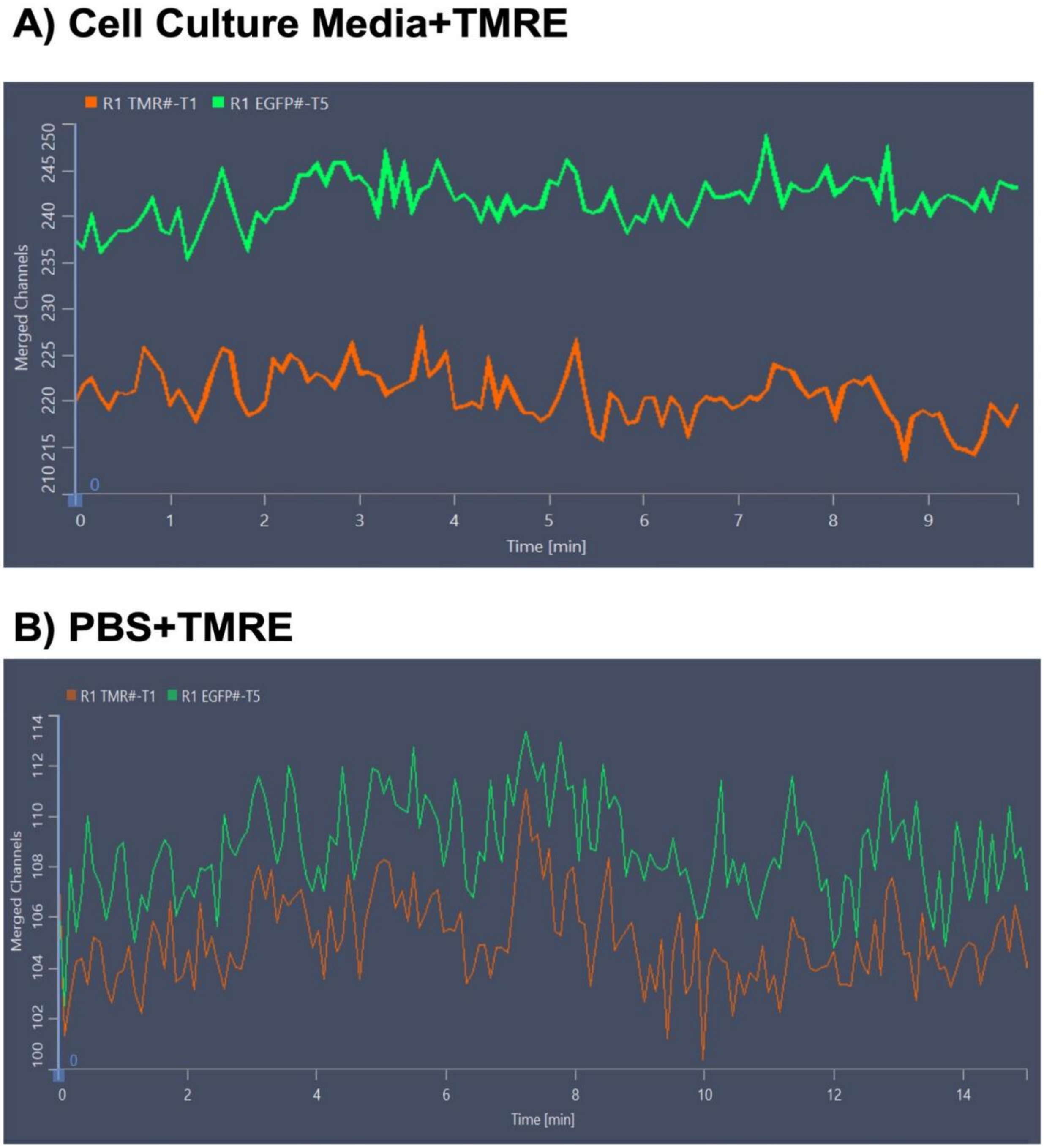


**Figure S6. Solutions containing either** (A) cell culture medium (DMEM + 10% FBS) or (B) PBS, with TMRE (10 nM), were imaged under the same confocal conditions used for live-cell experiments.

#### Photobleaching controls

To verify that fluorescence changes were specifically related to magnetic field modulation and not due to photobleaching or cellular stress, several control experiments were performed under identical imaging conditions. Photobleaching tests were performed on both HeLa wild type and HeLa–mtMagLOV2 cells in the absence of TMRE or FCCP to evaluate the stability of fluorescence under continuous illumination. Cells were imaged for 30 minutes using identical confocal settings (frame time 835 ms, image size 917 × 917 pixels, no averaging, interval 0 s) at three different laser powers (0.2%, 1%, and 10%).

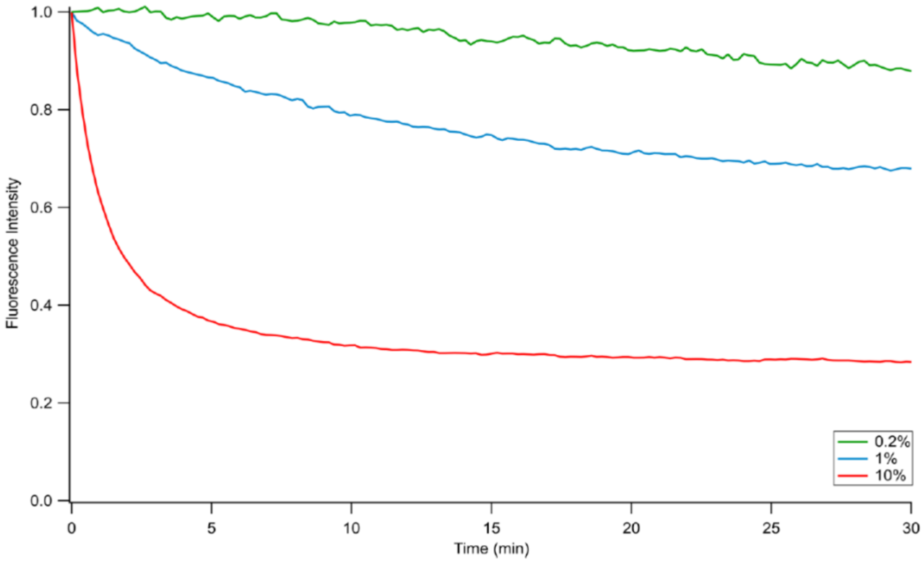


**Figure S7. Results above show the bleaching of mtMagLOV2 under differing illumination intensities.** Based on this data, all of our magnetic field data experiments were performed at or below 1% to avoid photobleaching effects.

#### ODMR Experiment

Three consecutive frequency-swept ODMR measurements (sweeps 1–3) were performed on the same HeLa-mtMagLOV2 field of view. Across all sweeps, a progressive decrease in overall fluorescence intensity was observed throughout each frequency sweep, attributable to cell toxicity due to RF induced heating under continuous 488 nm illumination during confocal acquisition.

In sweeps 1 and 2, fluorescence intensity was sufficient for analysis. After detrending the photobleaching baseline, an ODMR resonance feature was identified at approximately 549 MHz, consistent with the expected ESR condition for a free radical with $g \approx 2.00$ under the applied static field of $B_0 \approx 19.6$ mT (predicted f_res = 28.025 MHz $mT^{-1} \times 19.6$ mT ≈ 549 MHz). The small deviation between the observed resonance frequency and the predicted value likely reflects residual uncertainty in the calibrated field strength at the sample position.

By sweep 3, accumulated cell toxicity due to RF induced heating had reduced the fluorescence intensity to a level comparable to background, and no frequency-dependent modulation was discernible. This sweep therefore serves as an internal low-signal reference, confirming that the resonance feature observed in sweeps 1 and 2 is dependent on sufficient fluorescent signal rather than being an artifact of the RF delivery system.

The ODMR setup is illustrated in Figure S8. Cells used for ODMR measurements are shown in Figure S9. Representative ODMR spectra obtained from three consecutive frequency sweeps are shown in Figure S10.

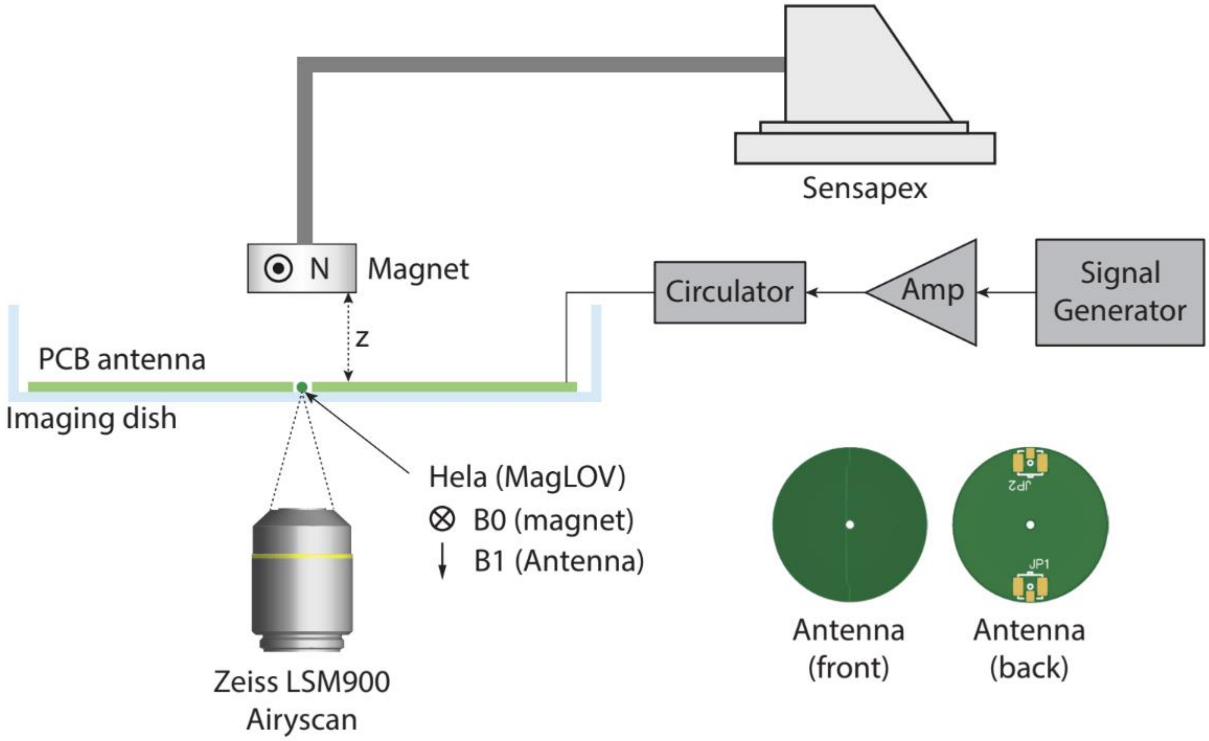
Sensapex
N
Magnet
Circulator
Amp
Signal Generator
z
PCB antenna
Imaging dish
Hela (MagLOV)
B0 (magnet)
B1 (Antenna)
JP2
JP1
Antenna (front)
Antenna (back)
Zeiss LSM900 Airyscan

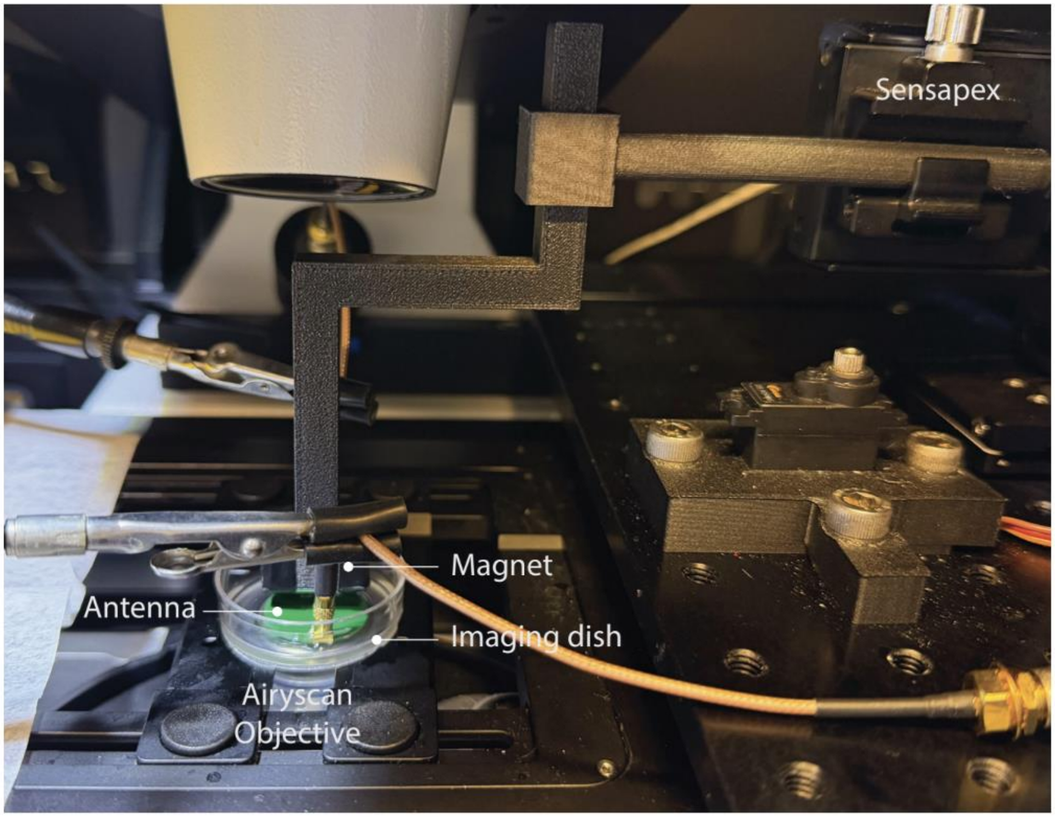
Sensapex
Magnet
Antenna
Imaging dish
Airyscan Objective

**Figure S8. Schematic of the ODMR experimental setup.** A PCB loop antenna positioned beneath the imaging dish delivered the RF field ($B_1$), while a permanent magnet generated the static magnetic field ($B_0$). HeLa cells expressing mtMagLOV2 were imaged using a Zeiss LSM900 Airyscan confocal microscope. The RF signal was generated by a signal generator, amplified, and routed through a circulator before delivery to the antenna. A Sensapex micromanipulator was used for precise positioning of the magnet relative to the sample.

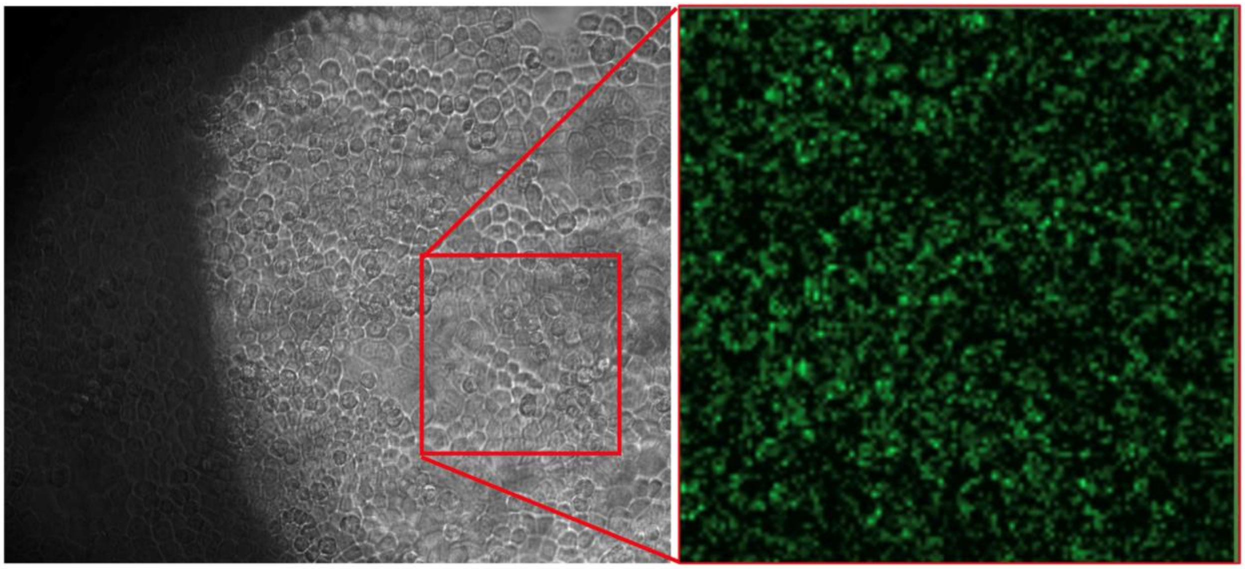

**Figure S9.** Confocal image of HeLa-mtMagLOV2 cells used in ODMR measurements, showing mitochondrial fluorescence prior to RF sweep.

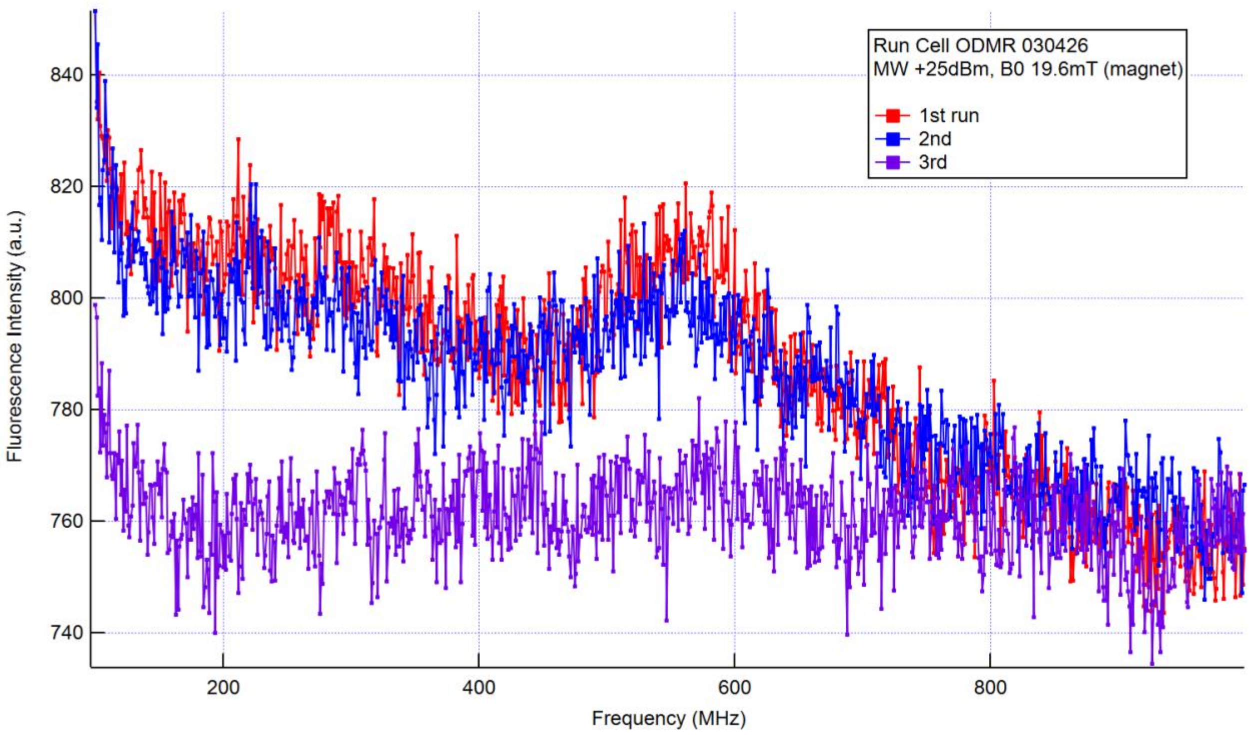


**Figure S10.** ODMR spectra from three consecutive frequency sweeps (100 MHz to 1 GHz). A resonance feature is visible at ~562 MHz in sweeps 1 and 2, consistent with the expected ESR condition. Sweep 3 shows near-background fluorescence due to accumulated cell toxicity.

#### DNA sequence and construct

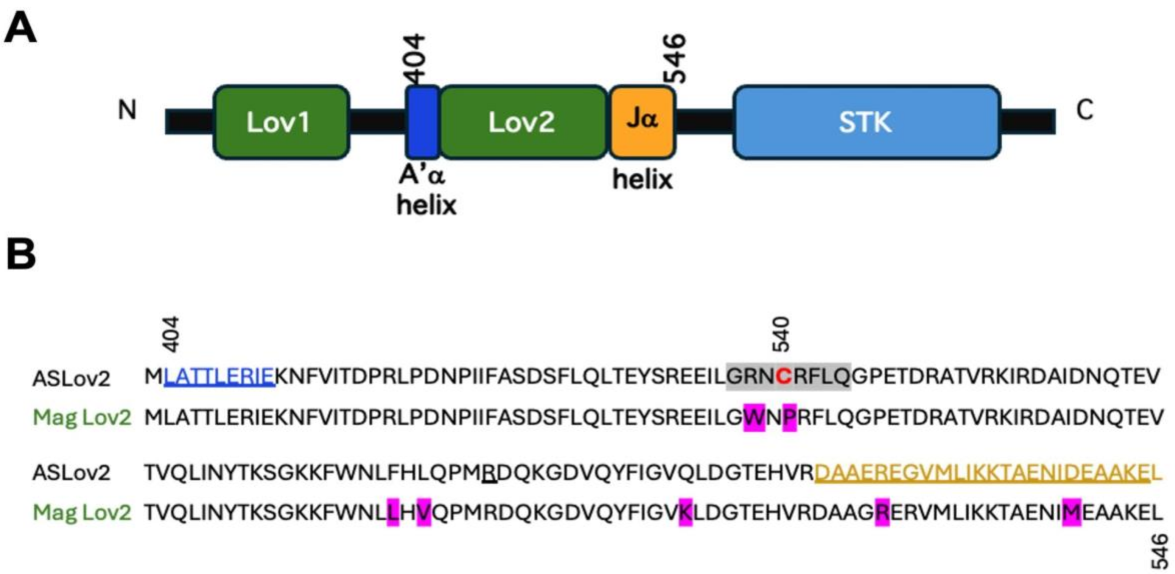


**Figure S11. Design and sequence features of the mitochondrial-targeted MagLOV2 construct.** (A) Schematic representation of the MagLOV2 fusion protein. The construct consists of tandem LOV domains (LOV1 and LOV2), followed by a Jα helix and a C-terminal STK domain. A mitochondrial targeting sequence (Cox8) was fused in-frame to the N-terminus to direct MagLOV2 to mitochondria via the TOM–TIM import pathway. (B) Amino acid sequence alignment comparing the native AsLOV2 domain with the engineered MagLOV2 variant. Residue substitutions introduced in MagLOV2 are highlighted, illustrating sequence modifications relative to the parental LOV2 domain.

MtMag: from IDT

CATTTCAGGTGTCGTGAGGATCGCCACC**ATGTCCGTCCTGACGCCGCTGCTG**
**CGGGGCTTGACAGGCTCGGCCCGGCGGCTCCCAGTGCCGCGCGCCAA**GGGTTC
TGGAAGTGGATCCGGAAGCGGAGCTAGC**ATGCTTGCTACTACTTTAGAACGTATA**
**GAGAAAAACTTCGTGATCACGGACCCGAGACTACCTGACAACCCTATAATTTTT**
**GCAAGTGACTCATTCCTTCAGTTGACTGAGTATTCTAGGGAAGAGATTCTAGGG**

**AGAAATCCTAGATTCTTGCAAGGACCAGAAACTGACCGTGCCACTGTGAGGAAAATCAGGGATGCGATCGACAACCAAACCGAGGTGACAGTGCAGCTAATAAATTACACTAAATCTGGCAAGAAGTTCTGGAACCTATTTCATGTGCAACCCATGAGAGACCAAAAAGGAGACGTACAGTACTTCATAGGGGTAAAGTTGGATGGTACTGAGCATGTTAGAGACGCGGCAGAACGTGAAAAGGTTATGTTAATAAAAAAGACCGCTGAAAACATAATGGAAGCGGCAAAGGAGTTGTTAGGTTAA**CTAACTTAAGCTAGCAACGGTTTC

Primers:

LHCox8: CCATTTCAGGTGTCGTGAGGATCGCCACCATGTCCGTCCTGACGC

LHHPAR: AACCGTTGCTAGCTTAAGTTAGTT

EF1A: CTCAAGCCTCAGACAGTGGTTC

pLPMtMagLOV2 (R10)

TGGAAGGGCTAATTCACTCCCAAAGAAGACAAGATATCCTTGATCTGTGGATCTACCACACACAAGGCTACTTCCCTGATTAGCAGAACTACACACCAGGGCCAGGGGTCAGATATCCACTGACCTTTGGATGGTGCTACAAGCTAGTACCAGTTGAGCCAGATAAGGTAGAAGAGGCCAATAAAGGAGAGAACACCAGCTTGTTACACCCTGTGAGCCTGCATGGGATGGATGACCCGGAGAGAGAAGTGTTAGAGTGGAGGTTTGACAGCCGCCTAGCATTTCATCACGTGGCCCGAGAGCTGCATCCGGAGTACTTCAAGAACTGCTGATATCGAGCTTGCTACAAGGGACTTTCCGCTGGGGACTTTCCAGGGAGGCGTGGCCTGGGCGGGACTGGGGAGTGGCGAGCCCTCAGATCCTGCATATAAGCAGCTGCTTTTTGCCTGTACTGGGTCTCTCTGGTTAGACCAGATCTGAGCCTGGGAGCTCTCTGGCTAACTAGGGAA

CCCACTGCTTAAGCCTCAATAAAGCTTGCCTTGAGTGCTTCAAGTAGTGTGTGCCCG
TCTGTTGTGTGACTCTGGTAACTAGAGATCCCTCAGACCCTTTTAGTCAGTGTGGAA
AATCTCTAGCAGTGGCGCCCGAACAGGGACTTGAAAGCGAAAGGGAAACCAGAGG
AGCTCTCTCGACGCAGGACTCGGCTTGCTGAAGCGCGCACGGCAAGAGGCGAGGGG
CGGCGACTGGTGAGTACGCCAAAAATTTTGACTAGCGGAGGCTAGAAGGAGAGAGA
TGGGTGCGAGAGCGTCAGTATTAAGCGGGGGAGAATTAGATCGCGATGGGAAAAAA
TTCGGTTAAGGCCAGGGGGAAAGAAAAAATATAAATTAAAACATATAGTATGGGCA
AGCAGGGAGCTAGAACGATTCGCAGTTAATCCTGGCCTGTTAGAAACATCAGAAGG
CTGTAGACAAATACTGGGACAGCTACAACCATCCCTTCAGACAGGATCAGAAGAAC
TTAGATCATTATATAATACAGTAGCAACCCTCTATTGTGTGCATCAAAGGATAGAGA
TAAAAGACACCAAGGAAGCTTTAGACAAGATAGAGGAAGAGCAAAACAAAAGTAA
GACCACCGCACAGCAAGCGGCCGGCCGCTGATCTTCAGACCTGGAGGAGGAGATAT
GAGGGACAATTGGAGAAGTGAATTATATAAATATAAAGTAGTAAAAATTGAACCAT
TAGGAGTAGCACCCACCAAGGCAAAGAGAAGAGTGGTGCAGAGAGAAAAAAGAGC
AGTGGGAATAGGAGCTTTGTTCCTTGGGTTCTTGGGAGCAGCAGGAAGCACTATGG
GCGCAGCGTCAATGACGCTGACGGTACAGGCCAGACAATTATTGTCTGGTATAGTGC
AGCAGCAGAACAATTTGCTGAGGGCTATTGAGGCGCAACAGCATCTGTTGCAACTC
ACAGTCTGGGGCATCAAGCAGCTCCAGGCAAGAATCCTGGCTGTGGAAAGATACCT
AAAGGATCAACAGCTCCTGGGGATTTGGGGTTGCTCTGGAAAACTCATTTGCACCAC
TGCTGTGCCTTGGAATGCTAGTTGGAGTAATAAATCTCTGGAACAGATTTGGAATCA
CACGACCTGGATGGAGTGGGACAGAGAAATTAACAATTACACAAGCTTAATACACT
CCTTAATTGAAGAATCGCAAAACCAGCAAGAAAAGAATGAACAAGAATTATTGGAA
TTAGATAAATGGGCAAGTTTGTGGAATTGGTTTAACATAACAAATTGGCTGTGGTAT

ATAAAATTATTCATAATGATAGTAGGAGGCTTGGTAGGTTTAAGAATAGTTTTTGCT
GTACTTTCTATAGTGAATAGAGTTAGGCAGGGATATTCACCATTATCGTTTCAGACC
CACCTCCCAACCCCGAGGGGACCCGACAGGCCCGAAGGAATAGAAGAAGAAGGTG
GAGAGAGAGACAGAGACAGATCCATTCGATTAGTGAACGGATCTCGACGGTATCGC
CTTTAAAAGAAAAGGGGGGATTGGGGGGTACAGTGCAGGGGAAAGAATAGTAGAC
ATAATAGCAACAGACATACAAACTAAAGAATTACAAAAACAAATTACAAAAATTCA
AAATTTTCGGGTTTATTACAGGGACAGCAGAGATCCAGTTTATCGATGAGGCCCTTT
CGTCTTCACTCGAGGTGCCCGTCAGTGGGCAGAGCGCACATCGCCCACAGTCCCCGA
GAAGTTGGGGGGAGGGGTCGGCAATTGAACCGGTGCCTAGAGAAGGTGGCGCGGG
GTAAACTGGGAAAGTGATGTCGTGTACTGGCTCCGCCTTTTTCCCGAGGGTGGGGGA
GAACCGTATATAAGTGCAGTAGTCGCCGTGAACGTTCTTTTTCGCAACGGGTTTGCC
GCCAGAACACAGGTAAGTGCCGTGTGTGGTTCCCGCGGGCCTGGCCTCTTTACGGGT
TATGGCCCTTGCGTGCCTTGAATTACTTCCACCTGGCTGCAGTACGTGATTCTTGATC
CCGAGCTTCGGGTTGGAAGTGGGTGGGAGAGTTCGAGGCCTTGCGCTTAAGGAGCC
CCTTCGCCTCGTGCTTGAGTTGAGGCCTGGCCTGGGCGCTGGGGCCGCCGCGTGCGA
ATCTGGTGGCACCTTCGCGCCTGTCTCGCTGCTTTCGATAAGTCTCTAGCCATTTAAA
ATTTTTGATGACCTGCTGCGACGCTTTTTTTCTGGCAAGATAGTCTTGTAAATGCGGG
CCAAGATCTGCACACTGGTATTTCGGTTTTTGGGGCCGCGGGCGGCGACGGGGCCCG
TGCGTCCCAGCGCACATGTTCGGCGAGGCGGGGCCTGCGAGCGCGGCCACCGAGAA
TCGGACGGGGGTAGTCTCAAGCTGGCCGGCCTGCTCTGGTGCCTGGCCTCGCGCCGC
CGTGTATCGCCCCGCCCTGGGCGGCAAGGCTGGCCCGGTCGGCACCAGTTGCGTGA
GCGGAAAGATGGCCGCTTCCCGGCCCTGCTGCAGGGAGCTCAAAATGGAGGACGCG
GCGCTCGGGAGAGCGGGCGGGTGAGTCACCCACACAAAGGAAAAGGGCCTTTCCGT

CCTCAGCCGTCGCTTCATGTGACTCCACGGAGTACCGGGCGCCGTCCAGGCACCTCGATTAGTTCTCGAGCTTTTGGAGTACGTCGTCTTTAGGTTGGGGGGAGGGGTTTTATGCGATGGAGTTTCCCCACACTGAGTGGGTGGAGACTGAAGTTAGGCCAGCTTGGCACTTGATGTAATTCTCCTTGGAATTTGCCCTTTTTGAGTTTGGATCTTGGTTCATTCTCAAGCCTCAGACAGTGGTTCAAAGTTTTTTTCTTC**CATTTCAGGTGTCGTGAGGATCGCCACCATGTCCGTCCTGACGCCGCTGCTGCTGCGGGGCTTGACAGGCTCGGCCCGGCGGCTCCCAGTGCCGCGCGCCAAGGGTTCTGGAAGTGGATCCGGAAGCGGAGCTAGCATGCTTGCTACTACTTTAGAACGTATAGAGAAAAACTTCGTGATCACGGACCCGAGACTACCTGACAACCCTATAATTTTTGCAAGTGACTCATTCCTTCAGTTGACTGAGTATTCTAGGGAAGAGATTCTAGGGAGAAATCCTAGATTCTTGCAAGGACCAGAAACTGACCGTGCCACTGTGAGGAAAATCAGGGATGCGATCGACAACCAAACCGAGGTGACAGTGCAGCTAATAAATTACACTAAATCTGGCAAGAAGTTCTGGAACCTATTTCATGTGCAACCCATGAGAGACCAAAAAGGAGACGTACAGTACTTCATAGGGGTAAAGTTGGATGGTACTGAGCATGTTAGAGACGCGGCAGAACGTGAAAAGGTTATGTTAATAAAAAAGACCGCTGAAAACATAATGGAAGCGGCAAAGGAGTTGTTAGGTTAACTAACTTAAGCTAGCAACGGTTTC**CCTCTAGCGGGATCAATTCCGCCCCCCCCCCTAACGTTACTGGCCGAAGCCGCTTGGAATAAGGCCGGTGTGCGTTTGTCTATATGTTATTTTCCACCATATTGCCGTCTTTTGGCAATGTGAGGGCCCGGAAACCTGGCCCTGTCTTCTTGACGAGCATTCCTAGGGGTCTTTCCCCTCTCGCCAAAGGAATGCAAGGTCTGTTGAATGTCGTGAAGGAAGCAGTTCCTCTGGAAGCTTCTTGAAGACAAACAACGTCTGTAGCGACCCTTTGCAGGCAGCGGAACCCCCCACCTGGCGACAGGTGCCTCTGCGGCCAAAAGCCACGTGTATAAGATACACCTGCAAAGGCGGCACAACCCCAGTGCCACGTTGTGAGTTGGATAGTTGTGGAAAGAGTCAAA

292 TGGCTCTCCTCAAGCGTATTCAACAAGGGGCTGAAGGATGCCCAGAAGGTACCCCA

293 TTGTATGGGATCTGATCTGGGGCCTCGGTGCACATGCTTTACATGTGTTTAGTCGAG

294 GTTAAAAAACGTCTAGGCCCCCCGAACCACGGGGACGTGGTTTTCCTTTGAAAAAC

295 ACGATAATACCATGAAAAAGCCTGAACTCACCGCGACGTCTGTCGAGAAGTTTCTG

296 ATCGAAAAGTTCGACAGCGTCTCCGACCTGATGCAGCTCTCGGAGGGCGAAGAATC

297 TCGTGCTTTCAGCTTCGATGTAGGAGGGCGTGGATATGTCCTGCGGGTAAATAGCTG

298 CGCCGATGGTTTCTACAAAGATCGTTATGTTTATCGGCACTTTGCATCGGCCGCGCTC

299 CCGATTCCGGAAGTGCTTGACATTGGGGAATTTAGCGAGAGCCTGACCTATTGCATC

300 TCCCGCCGTGCACAGGGTGTCACGTTGCAAGACCTGCCTGAAACCGAACTGCCCGCT

301 GTTCTGCAGCCGGTCGCGGAGGCCATGGATGCGATCGCTGCGGCCGATCTTAGCCA

302 GACGAGCGGGTTCGGCCCATTCGGACCGCAAGGAATCGGTCAATACACTACATGGC

303 GTGATTTCATATGCGCGATTGCTGATCCCCATGTGTATCACTGGCAAACTGTGATGG

304 ACGACACCGTCAGTGCGTCCGTCGCGCAGGCTCTCGATGAGCTGATGCTTTGGGCCG

305 AGGACTGCCCCGAAGTCCGGCACCTCGTGCACGCGGATTTCGGCTCCAACAATGTCC

306 TGACGGACAATGGCCGCATAACAGCGGTCATTGACTGGAGCGAGGCGATGTTCGGG

307 GATTCCCAATACGAGGTCGCCAACATCTTCTTCTGGAGGCCGTGGTTGGCTTGTATG

308 GAGCAGCAGACGCGCTACTTCGAGCGGAGGCATCCGGAGCTTGCAGGATCGCCGCG

309 GCTCCGGGCGTATATGCTCCGCATTGGTCTTGACCAACTCTATCAGAGCTTGGTTGA

310 CGGCAATTTCGATGATGCAGCTTGGGCGCAGGGTCGATGCGACGCAATCGTCCGATC

311 CGGAGCCGGGACTGTCGGGCGTACACAAATCGCCCGCAGAAGCGCGGCCGTCTGGA

312 CCGATGGCTGTGTAGAAGTACTCGCCGATAGTGGAAACCGACGCCCCAGCACTCGT

313 CCGAGGGCAAAGGAATAGCCCGGGCGGGGCGCGTCTGGAACAATCAACCTCTGGAT

314 TACAAAATTTGTGAAAGATTGACTGGTATTCTTAACTATGTTGCTCCTTTTACGCTAT

GTGGATACGCTGCTTTAATGCCTTTGTATCATGCTATTGCTTCCCGTATGGCTTTCAT
TTTCTCCTCCTTGTATAAATCCTGGTTGCTGTCTCTTTATGAGGAGTTGTGGCCCGTT
GTCAGGCAACGTGGCGTGGTGTGCACTGTGTTTGCTGACGCAACCCCCACTGGTTGG
GGCATTGCCACCACCTGTCAGCTCCTTTCCGGGACTTTCGCTTTCCCCCTCCCTATTG
CCACGGCGGAACTCATCGCCGCCTGCCTTGCCCGCTGCTGGACAGGGGCTCGGCTGT
TGGGCACTGACAATTCCGTGGTGTTGTCGGGGAAGCTGACGTCCTTTCCATGGCTGC
TCGCCTGTGTTGCCACCTGGATTCTGCGCGGGACGTCCTTCTGCTACGTCCCTTCGGC
CCTCAATCCAGCGGACCTTCCTTCCCGCGGCCTGCTGCCGGCTCTGCGGCCTCTTCCG
CGTCTTCGCCTTCGCCCTCAGACGAGTCGGATCTCCCTTTGGGCCGCCTCCCCGCCTG
GAATTAATTCTGCAGTCGAGACCTAGAAAAACATGGAGCAATCACAAGTAGCAATA
CAGCAGCTACCAATGCTGATTGTGCCTGGCTAGAAGCACAAGAGGAGGAGGAGGTG
GGTTTTCCAGTCACACCTCAGGTACCTTTAAGACCAATGACTTACAAGGCAGCTGTA
GATCTTAGCCACTTTTTAAAAGAAAAGAGGGGACTGGAAGGGCTAATTCACTCCCA
ACGAAGACAAGATATCCTTGATCTGTGGATCTACCACACACAAGGCTACTTCCCTGA
TTAGCAGAACTACACACCAGGGCCAGGGGTCAGATATCCACTGACCTTTGGATGGT
GCTACAAGCTAGTACCAGTTGAGCCAGATAAGGTAGAAGAGGCCAATAAAGGAGA
GAACACCAGCTTGTTACACCCTGTGAGCCTGCATGGGATGGATGACCCGGAGAGAG
AAGTGTTAGAGTGGAGGTTTGACAGCCGCCTAGCATTTCATCACGTGGCCCGAGAGC
TGCATCCGGAGTACTTCAAGAACTGCTGATATCGAGCTTGCTACAAGGGACTTTCCG
CTGGGGACTTTCCAGGGAGGCGTGGCCTGGGCGGGACTGGGGAGTGGCGAGCCCTC
AGATCCTGCATATAAGCAGCTGCTTTTTGCCTGTACTGGGTCTCTCTGGTTAGACCAG
ATCTGAGCCTGGGAGCTCTCTGGCTAACTAGGGAACCCACTGCTTAAGCCTCAATAA
AGCTTGCCTTGAGTGCTTCAAGTAGTGTGTGCCCGTCTGTTGTGTGACTCTGGTAACT

AGAGATCCCTCAGACCCTTTTAGTCAGTGTGGAAAATCTCTAGCAGTAGTAGTTCAT
GTCATCTTATTATTCAGTATTTATAACTTGCAAAGAAATGAATATCAGAGAGTGAGA
GGCCTTGACATTGCTAGCGTTTACCGTCGACCTCTAGCTAGAGCTTGGCGTAATCAT
GGTCATAGCTGTTTCCTGTGTGAAATTGTTATCCGCTCACAATTCCACACAACATAC
GAGCCGGAAGCATAAAGTGTAAAGCCTGGGGTGCCTAATGAGTGAGCTAACTCACA
TTAATTGCGTTGCGCTCACTGCCCGCTTTCCAGTCGGGAAACCTGTCGTGCCAGCTG
CATTAATGAATCGGCCAACGCGCGGGGAGAGGCGGTTTGCGTATTGGGCGCTCTTCC
GCTTCCTCGCTCACTGACTCGCTGCGCTCGGTCGTTCGGCTGCGGCGAGCGGTATCA
GCTCACTCAAAGGCGGTAATACGGTTATCCACAGAATCAGGGGATAACGCAGGAAA
GAACATGTGAGCAAAAGGCCAGCAAAAGGCCAGGAACCGTAAAAAGGCCGCGTTG
CTGGCGTTTTTCCATAGGCTCCGCCCCCCTGACGAGCATCACAAAAATCGACGCTCA
AGTCAGAGGTGGCGAAACCCGACAGGACTATAAAGATACCAGGCGTTTCCCCCTGG
AAGCTCCCTCGTGCGCTCTCCTGTTCCGACCCTGCCGCTTACCGGATACCTGTCCGCC
TTTCTCCCTTCGGGAAGCGTGGCGCTTTCTCATAGCTCACGCTGTAGGTATCTCAGTT
CGGTGTAGGTCGTTCGCTCCAAGCTGGGCTGTGTGCACGAACCCCCCGTTCAGCCCG
ACCGCTGCGCCTTATCCGGTAACTATCGTCTTGAGTCCAACCCGGTAAGACACGACT
TATCGCCACTGGCAGCAGCCACTGGTAACAGGATTAGCAGAGCGAGGTATGTAGGC
GGTGCTACAGAGTTCTTGAAGTGGTGGCCTAACTACGGCTACACTAGAAGAACAGT
ATTTGGTATCTGCGCTCTGCTGAAGCCAGTTACCTTCGGAAAAAGAGTTGGTAGCTC
TTGATCCGGCAAACAAACCACCGCTGGTAGCGGTGGTTTTTTTGTTTGCAAGCAGCA
GATTACGCGCAGAAAAAAAGGATCTCAAGAAGATCCTTTGATCTTTTCTACGGGGTC
TGACGCTCAGTGGAACGAAAACTCACGTTAAGGGATTTTGGTCATGAGATTATCAAA
AAGGATCTTCACCTAGATCCTTTTAAATTAAAAATGAAGTTTTAAATCAATCTAAAG

361 TATATATGAGTAAACTTGGTCTGACAGTTACCAATGCTTAATCAGTGAGGCACCTAT
362 CTCAGCGATCTGTCTATTTCGTTCATCCATAGTTGCCTGACTCCCCGTCGTGTAGATA
363 ACTACGATACGGGAGGGCTTACCATCTGGCCCCAGTGCTGCAATGATACCGCGAGA
364 CCCACGCTCACCGGCTCCAGATTTATCAGCAATAAACCAGCCAGCCGGAAGGGCCG
365 AGCGCAGAAGTGGTCCTGCAACTTTATCCGCCTCCATCCAGTCTATTAATTGTTGCC
366 GGGAAGCTAGAGTAAGTAGTTCGCCAGTTAATAGTTTGCGCAACGTTGTTGCCATTG
367 CTACAGGCATCGTGGTGTCACGCTCGTCGTTTGGTATGGCTTCATTCAGCTCCGGTTC
368 CCAACGATCAAGGCGAGTTACATGATCCCCCATGTTGTGCAAAAAAGCGGTTAGCTC
369 CTTCGGTCCTCCGATCGTTGTCAGAAGTAAGTTGGCCGCAGTGTTATCACTCATGGTT
370 ATGGCAGCACTGCATAATTCTCTTACTGTCATGCCATCCGTAAGATGCTTTTCTGTGA
371 CTGGTGAGTACTCAACCAAGTCATTCTGAGAATAGTGTATGCGGCGACCGAGTTGCT
372 CTTGCCCGGCGTCAATACGGGATAATACCGCGCCACATAGCAGAACTTTAAAAGTG
373 CTCATCATTGGAAAACGTTCTTCGGGGCGAAAACTCTCAAGGATCTTACCGCTGTTG
374 AGATCCAGTTCGATGTAACCCACTCGTGCACCCAACTGATCTTCAGCATCTTTTACTT
375 TCACCAGCGTTTCTGGGTGAGCAAAAACAGGAAGGCAAAATGCCGCAAAAAAGGG
376 AATAAGGGCGACACGGAAATGTTGAATACTCATACTCTTCCTTTTTCAATATTATTG
377 AAGCATTTATCAGGGTTATTGTCTCATGAGCGGATACATATTTGAATGTATTTAGAA
378 AAATAAACAAATAGGGGTTCCGCGCACATTTCCCCGAAAAGTGCCACCTGACGTCG
379 ACGGATCGGGAGATCAACTTGTTTATTGCAGCTTATAATGGTTACAAATAAAGCAAT
380 AGCATCACAAATTTCACAAATAAAGCATTTTTTTCACTGCATTCTAGTTGTGGTTTGT
381 CCAAACTCATCAATGTATCTTATCATGTCTGGATCAACTGGATAACTCAAGCTAACC
382 AAAATCATCCCAAACTTCCCACCCCATACCCTATTACCACTGCCAATTACCTGTGGT

TTCATTTACTCTAAACCTGTGATTCCTCTGAATTATTTCATTTAAAGAAATTGTATT
TGTTAAATATGTACTACAAACTTAGTAGT

pLPMtMagLOV2-3Flag (R10-3F)

TGGAAGGGCTAATTCACTCCCAAAGAAGACAAGATATCCTTGATCTGTGGATCTACC
ACACACAAGGCTACTTCCCTGATTAGCAGAACTACACACCAGGGCCAGGGGTCAGA
TATCCACTGACCTTTGGATGGTGCTACAAGCTAGTACCAGTTGAGCCAGATAAGGTA
GAAGAGGCCAATAAAGGAGAGAACACCAGCTTGTTACACCCTGTGAGCCTGCATGG
GATGGATGACCCGGAGAGAGAAGTGTTAGAGTGGAGGTTTGACAGCCGCCTAGCAT
TTCATCACGTGGCCCGAGAGCTGCATCCGGAGTACTTCAAGAACTGCTGATATCGAG
CTTGCTACAAGGGACTTTCCGCTGGGGACTTTCCAGGGAGGCGTGGCCTGGGCGGG
ACTGGGGAGTGGCGAGCCCTCAGATCCTGCATATAAGCAGCTGCTTTTTGCCTGTAC
TGGGTCTCTCTGGTTAGACCAGATCTGAGCCTGGGAGCTCTCTGGCTAACTAGGGAA
CCCACTGCTTAAGCCTCAATAAAGCTTGCCTTGAGTGCTTCAAGTAGTGTGTGCCCG
TCTGTTGTGTGACTCTGGTAACTAGAGATCCCTCAGACCCTTTTAGTCAGTGTGGAA
AATCTCTAGCAGTGGCGCCCGAACAGGGACTTGAAAGCGAAAGGGAAACCAGAGG
AGCTCTCTCGACGCAGGACTCGGCTTGCTGAAGCGCGCACGGCAAGAGGCGAGGGG
CGGCGACTGGTGAGTACGCCAAAAATTTTGACTAGCGGAGGCTAGAAGGAGAGAGA
TGGGTGCGAGAGCGTCAGTATTAAGCGGGGGAGAATTAGATCGCGATGGGAAAAAA
TTCGGTTAAGGCCAGGGGGAAAGAAAAAATATAAATTAAAACATATAGTATGGGCA
AGCAGGGAGCTAGAACGATTCGCAGTTAATCCTGGCCTGTTAGAAACATCAGAAGG
CTGTAGACAAATACTGGGACAGCTACAACCATCCCTTCAGACAGGATCAGAAGAAC
TTAGATCATTATATAATACAGTAGCAACCCTCTATTGTGTGCATCAAGGATAGAGA
TAAAAGACACCAAGGAAGCTTTAGACAAGATAGAGGAAGAGCAAAACAAAAGTAA

GACCACCGCACAGCAAGCGGCCGGCCGCTGATCTTCAGACCTGGAGGAGGAGATAT
GAGGGACAATTGGAGAAGTGAATTATATAAATATAAAGTAGTAAAAATTGAACCAT
TAGGAGTAGCACCCACCAAGGCAAAGAGAAGAGTGGTGCAGAGAGAAAAAAGAGC
AGTGGGAATAGGAGCTTTGTTCCTTGGGTTCTTGGGAGCAGCAGGAAGCACTATGG
GCGCAGCGTCAATGACGCTGACGGTACAGGCCAGACAATTATTGTCTGGTATAGTGC
AGCAGCAGAACAATTTGCTGAGGGCTATTGAGGCGCAACAGCATCTGTTGCAACTC
ACAGTCTGGGGCATCAAGCAGCTCCAGGCAAGAATCCTGGCTGTGGAAAGATACCT
AAAGGATCAACAGCTCCTGGGGATTTGGGGTTGCTCTGGAAAACTCATTTGCACCAC
TGCTGTGCCTTGGAATGCTAGTTGGAGTAATAAATCTCTGGAACAGATTTGGAATCA
CACGACCTGGATGGAGTGGGACAGAGAAATTAACAATTACACAAGCTTAATACACT
CCTTAATTGAAGAATCGCAAAACCAGCAAGAAAAGAATGAACAAGAATTATTGGAA
TTAGATAAATGGGCAAGTTTGTGGAATTGGTTTAACATAACAAATTGGCTGTGGTAT
ATAAAATTATTCATAATGATAGTAGGAGGCTTGGTAGGTTTAAGAATAGTTTTTGCT
GTACTTTCTATAGTGAATAGAGTTAGGCAGGGATATTCACCATTATCGTTTCAGACC
CACCTCCCAACCCCGAGGGGACCCGACAGGCCCGAAGGAATAGAAGAAGAAGGTG
GAGAGAGAGACAGAGACAGATCCATTCGATTAGTGAACGGATCTCGACGGTATCGC
CTTTAAAAGAAAAGGGGGGATTGGGGGGTACAGTGCAGGGGAAAGAATAGTAGAC
ATAATAGCAACAGACATACAAACTAAAGAATTACAAAAACAAATTACAAAAATTCA
AAATTTTCGGGTTTATTACAGGGACAGCAGAGATCCAGTTTATCGATGAGGCCCTTT
CGTCTTCACTCGAGGTGCCCGTCAGTGGGCAGAGCGCACATCGCCCACAGTCCCCGA
GAAGTTGGGGGGAGGGGTCGGCAATTGAACCGGTGCCTAGAGAAGGTGGCGCGGG
GTAAACTGGGAAAGTGATGTCGTGTACTGGCTCCGCCTTTTTCCCGAGGGTGGGGGA
GAACCGTATATAAGTGCAGTAGTCGCCGTGAACGTTCTTTTTCGCAACGGGTTTGCC

GCCAGAACACAGGTAAGTGCCGTGTGTGGTTCCCGCGGGCCTGGCCTCTTTACGGGT
TATGGCCCTTGCGTGCCTTGAATTACTTCCACCTGGCTGCAGTACGTGATTCTTGATC
CCGAGCTTCGGGTTGGAAGTGGGTGGGAGAGTTCGAGGCCTTGCGCTTAAGGAGCC
CCTTCGCCTCGTGCTTGAGTTGAGGCCTGGCCTGGGCGCTGGGGCCGCCGCGTGCGA
ATCTGGTGGCACCTTCGCGCCTGTCTCGCTGCTTTCGATAAGTCTCTAGCCATTTAAA
ATTTTTGATGACCTGCTGCGACGCTTTTTTTCTGGCAAGATAGTCTTGTAAATGCGGG
CCAAGATCTGCACACTGGTATTTCGGTTTTTGGGGCCGCGGGCGGCGACGGGGCCCG
TGCGTCCCAGCGCACATGTTCGGCGAGGCGGGGCCTGCGAGCGCGGCCACCGAGAA
TCGGACGGGGGTAGTCTCAAGCTGGCCGGCCTGCTCTGGTGCCTGGCCTCGCGCCGC
CGTGTATCGCCCCGCCCTGGGCGGCAAGGCTGGCCCGGTCGGCACCAGTTGCGTGA
GCGGAAAGATGGCCGCTTCCCGGCCCTGCTGCAGGGAGCTCAAAATGGAGGACGCG
GCGCTCGGGAGAGCGGGCGGGTGAGTCACCCACACAAAGGAAAAGGGCCTTTCCGT
CCTCAGCCGTCGCTTCATGTGACTCCACGGAGTACCGGGCGCCGTCCAGGCACCTCG
ATTAGTTCTCGAGCTTTTGGAGTACGTCGTCTTTAGGTTGGGGGGAGGGGTTTTATG
CGATGGAGTTTCCCCACACTGAGTGGGTGGAGACTGAAGTTAGGCCAGCTTGGCACT
TGATGTAATTCTCCTTGGAATTTGCCCTTTTTGAGTTTGGATCTTGGTTCATTCTCAA
GCCTCAGACAGTGGTTCAAAGTTTTTTTCTTCCATTTCAGGTGTCGTGAGGATCTCTA
GAGTTTAAACGCTAGCATGTCCGTCCTGACGCCGCTGCTGCTGCGGGGCTTGACAGG
CTCGGCCCGGCGGCTCCCAGTGCCGCGCGCCAAGATCCATTCGTTGGGGGATCCCAT
GCTTGCTACTACTTTAGAACGTATAGAGAAAAACTTCGTGATCACGGACCCGAGACT
ACCTGACAACCCTATAATTTTTGCAAGTGACTCATTCCTTCAGTTGACTGAGTATTCT
AGGGAAGAGATTCTAGGGAGAAATCCTAGATTCTTGCAAGGACCAGAAACTGACCG
TGCCACTGTGAGGAAAATCAGGGATGCGATCGACAACCAAACCGAGGTGACAGTGC

AGCTAATAAATTACACTAAATCTGGCAAGAAGTTCTGGAACCTATTTCATGTGCAAC
CCATGAGAGACCAAAAAGGAGACGTACAGTACTTCATAGGGGTAAAGTTGGATGGT
ACTGAGCATGTTAGAGACGCGGCAGAACGTGAAAAGGTTATGTTAATAAAAAAGAC
CGCTGAAAACATAATGGAAGCGGCAAAGGAGTTGACTAGTGACTACAAAGACCATG
ACGGTGATTATAAAGATCATGACATCGACTACAAGGATGACGATGACAAGTAGAAG
CTTGGTTAACTAACTTAAGCTAGCAACGGTTTCCCTCTAGCGGGATCAATTCCGCCC
CCCCCCCTAACGTTACTGGCCGAAGCCGCTTGGAATAAGGCCGGTGTGCGTTTGTC
TATATGTTATTTTCCACCATATTGCCGTCTTTTGGCAATGTGAGGGCCCGGAAACCTG
GCCCTGTCTTCTTGACGAGCATTCCTAGGGGTCTTTCCCCTCTCGCCAAAGGAATGC
AAGGTCTGTTGAATGTCGTGAAGGAAGCAGTTCCTCTGGAAGCTTCTTGAAGACAAA
CAACGTCTGTAGCGACCCTTTGCAGGCAGCGGAACCCCCCACCTGGCGACAGGTGC
CTCTGCGGCCAAAAGCCACGTGTATAAGATACACCTGCAAAGGCGGCACAACCCCA
GTGCCACGTTGTGAGTTGGATAGTTGTGGAAAGAGTCAAATGGCTCTCCTCAAGCGT
ATTCAACAAGGGGCTGAAGGATGCCCAGAAGGTACCCCATTGTATGGGATCTGATC
TGGGGCCTCGGTGCACATGCTTTACATGTGTTTAGTCGAGGTTAAAAAACGTCTAG
GCCCCCCGAACCACGGGGACGTGGTTTTCCTTTGAAAAACACGATAATACCATGGCC
ACCGAGTACAAGCCCACGGTGCGCCTCGCCACCCGCGACGACGTCCCCCGGGCCGT
ACGCACCCTCGCCGCCGCGTTCGCCGACTACCCCGCCACGCGCCACACCGTCGACCC
GGACCGCCACATCGAGCGGGTCACCGAGCTGCAAGAACTCTTCCTCACGCGCGTCG
GGCTCGACATCGGCAAGGTGTGGGTCGCGGACGACGGCGCCGCGGTGGCGGTCTGG
ACCACGCCGGAGAGCGTCGAAGCGGGGGCGGTGTTCGCCGAGATCGGCTCGCGCAT
GGCCGAGTTGAGCGGTTCCCGGCTGGCCGCGCAGCAACAGATGGAAGGCCTCCTGG
CGCCGCACCGGCCCAAGGAGCCCGCGTGGTTCCTGGCCACCGTCGGCGTCTCGCCCG

ACCACCAGGGCAAGGGTCTGGGCAGCGCCGTCGTGCTCCCCGGAGTGGAGGCGGCCGAGCGCGCTGGGGTGCCCGCCTTCCTGGAGACCTCCGCGCCCCGCAACCTCCCCTTCTACGAGCGGCTCGGCTTCACCGTCACCGCCGACGTCGAGGTGCCCGAAGGACCGCGCACCTGGTGCATGACCCGCAAGCCCGGTGCCTGAGTTCGCGTCTGGAACAATCAACCTCTGGATTACAAAATTTGTGAAAGATTGACTGGTATTCTTAACTATGTTGCTCCTTTTACGCTATGTGGATACGCTGCTTTAATGCCTTTGTATCATGCTATTGCTTCCCGTATGGCTTTCATTTTCTCCTCCTTGTATAAATCCTGGTTGCTGTCTCTTTATGAGGAGTTGTGGCCCGTTGTCAGGCAACGTGGCGTGGTGTGCACTGTGTTTGCTGACGCAACCCCCACTGGTTGGGGCATTGCCACCACCTGTCAGCTCCTTTCCGGGACTTTCGCTTTCCCCCTCCCTATTGCCACGGCGGAACTCATCGCCGCCTGCCTTGCCCGCTGCTGGACAGGGGCTCGGCTGTTGGGCACTGACAATTCCGTGGTGTTGTCGGGGAAGCTGACGTCCTTTCCATGGCTGCTCGCCTGTGTTGCCACCTGGATTCTGCGCGGGACGTCCTTCTGCTACGTCCCTTCGGCCCTCAATCCAGCGGACCTTCCTTCCCGCGGCCTGCTGCCGGCTCTGCGGCCTCTTCCGCGTCTTCGCCTTCGCCCTCAGACGAGTCGGATCTCCCTTTGGGCCGCCTCCCCGCCTGGAATTAATTCTGCAGTCGAGACCTAGAAAAACATGGAGCAATCACAAGTAGCAATACAGCAGCTACCAATGCTGATTGTGCCTGGCTAGAAGCACAAGAGGAGGAGGAGGTGGGTTTTCCAGTCACACCTCAGGTACCTTTAAGACCAATGACTTACAAGGCAGCTGTAGATCTTAGCCACTTTTTAAAAGAAAAGAGGGGACTGGAAGGGCTAATTCACTCCCAACGAAGACAAGATATCCTTGATCTGTGGATCTACCACACACAAGGCTACTTCCCTGATTAGCAGAACTACACACCAGGGCCAGGGGTCAGATATCCACTGACCTTTGGATGGTGCTACAAGCTAGTACCAGTTGAGCCAGATAAGGTAGAAGAGGCCAATAAAGGAGAGAACACCAGCTTGTTACACCCTGTGAGCCTGCATGGGATGGATGACCCGGAGAGAGAAGTGTTAGAGTGGAGGTTTGACAGCCGCCTAGCATTTCATCACGTGGCCC

498 GAGAGCTGCATCCGGAGTACTTCAAGAACTGCTGATATCGAGCTTGCTACAAGGGA

499 CTTTCCGCTGGGGACTTTCCAGGGAGGCGTGGCCTGGGCGGGACTGGGGAGTGGCG

500 AGCCCTCAGATCCTGCATATAAGCAGCTGCTTTTTGCCTGTACTGGGTCTCTCTGGTT

501 AGACCAGATCTGAGCCTGGGAGCTCTCTGGCTAACTAGGGAACCCACTGCTTAAGC

502 CTCAATAAAGCTTGCCTTGAGTGCTTCAAGTAGTGTGTGCCCGTCTGTTGTGTGACTC

503 TGGTAACTAGAGATCCCTCAGACCCTTTTAGTCAGTGTGGAAAATCTCTAGCAGTAG

504 TAGTTCATGTCATCTTATTATTCAGTATTTATAACTTGCAAAGAAATGAATATCAGA

505 GAGTGAGAGGCCTTGACATTGCTAGCGTTTACCGTCGACCTCTAGCTAGAGCTTGGC

506 GTAATCATGGTCATAGCTGTTTCCTGTGTGAAATTGTTATCCGCTCACAATTCCACAC

507 AACATACGAGCCGGAAGCATAAAGTGTAAAGCCTGGGGTGCCTAATGAGTGAGCTA

508 ACTCACATTAATTGCGTTGCGCTCACTGCCCGCTTTCCAGTCGGGAAACCTGTCGTG

509 CCAGCTGCATTAATGAATCGGCCAACGCGCGGGGAGAGGCGGTTTGCGTATTGGGC

510 GCTCTTCCGCTTCCTCGCTCACTGACTCGCTGCGCTCGGTCGTTCGGCTGCGGCGAGC

511 GGTATCAGCTCACTCAAAGGCGGTAATACGGTTATCCACAGAATCAGGGGATAACG

512 CAGGAAAGAACATGTGAGCAAAAGGCCAGCAAAAGGCCAGGAACCGTAAAAAGGC

513 CGCGTTGCTGGCGTTTTTCCATAGGCTCCGCCCCCCTGACGAGCATCACAAAAATCG

514 ACGCTCAAGTCAGAGGTGGCGAAACCCGACAGGACTATAAAGATACCAGGCGTTTC

515 CCCCTGGAAGCTCCCTCGTGCGCTCTCCTGTTCCGACCCTGCCGCTTACCGGATACCT

516 GTCCGCCTTTCTCCCTTCGGGAAGCGTGGCGCTTTCTCATAGCTCACGCTGTAGGTAT

517 CTCAGTTCGGTGTAGGTCGTTCGCTCCAAGCTGGGCTGTGTGCACGAACCCCCCGTT

518 CAGCCCGACCGCTGCGCCTTATCCGGTAACTATCGTCTTGAGTCCAACCCGGTAAGA

519 CACGACTTATCGCCACTGGCAGCAGCCACTGGTAACAGGATTAGCAGAGCGAGGTA

520 TGTAGGCGGTGCTACAGAGTTCTTGAAGTGGTGGCCTAACTACGGCTACACTAGAAG

AACAGTATTTGGTATCTGCGCTCTGCTGAAGCCAGTTACCTTCGGAAAAAGAGTTGG
TAGCTCTTGATCCGGCAAACAAACCACCGCTGGTAGCGGTGGTTTTTTTGTTTGCAA
GCAGCAGATTACGCGCAGAAAAAAAGGATCTCAAGAAGATCCTTTGATCTTTTCTAC
GGGGTCTGACGCTCAGTGGAACGAAAACTCACGTTAAGGGATTTTGGTCATGAGATT
ATCAAAAAGGATCTTCACCTAGATCCTTTTAAATTAAAAATGAAGTTTTAAATCAAT
CTAAAGTATATATGAGTAAACTTGGTCTGACAGTTACCAATGCTTAATCAGTGAGGC
ACCTATCTCAGCGATCTGTCTATTTCGTTCATCCATAGTTGCCTGACTCCCCGTCGTG
TAGATAACTACGATACGGGAGGGCTTACCATCTGGCCCCAGTGCTGCAATGATACCG
CGAGACCCACGCTCACCGGCTCCAGATTTATCAGCAATAAACCAGCCAGCCGGAAG
GGCCGAGCGCAGAAGTGGTCCTGCAACTTTATCCGCCTCCATCCAGTCTATTAATTG
TTGCCGGGAAGCTAGAGTAAGTAGTTCGCCAGTTAATAGTTTGCGCAACGTTGTTGC
CATTGCTACAGGCATCGTGGTGTCACGCTCGTCGTTTGGTATGGCTTCATTCAGCTCC
GGTTCCCAACGATCAAGGCGAGTTACATGATCCCCCATGTTGTGCAAAAAAGCGGTT
AGCTCCTTCGGTCCTCCGATCGTTGTCAGAAGTAAGTTGGCCGCAGTGTTATCACTC
ATGGTTATGGCAGCACTGCATAATTCTCTTACTGTCATGCCATCCGTAAGATGCTTTT
CTGTGACTGGTGAGTACTCAACCAAGTCATTCTGAGAATAGTGTATGCGGCGACCGA
GTTGCTCTTGCCCGGCGTCAATACGGGATAATACCGCGCCACATAGCAGAACTTTAA
AAGTGCTCATCATTGGAAAACGTTCTTCGGGGCGAAAACTCTCAAGGATCTTACCGC
TGTTGAGATCCAGTTCGATGTAACCCACTCGTGCACCCAACTGATCTTCAGCATCTTT
TACTTTCACCAGCGTTTCTGGGTGAGCAAAAACAGGAAGGCAAAATGCCGCAAAAA
AGGGAATAAGGGCGACACGGAAATGTTGAATACTCATACTCTTCCTTTTTCAATATT
ATTGAAGCATTTATCAGGGTTATTGTCTCATGAGCGGATACATATTTGAATGTATTTA
GAAAAATAAACAAATAGGGGTTCCGCGCACATTTCCCCGAAAAGTGCCACCTGACG

TCGACGGATCGGGAGATCAACTTGTTTATTGCAGCTTATAATGGTTACAAATAAAGC
AATAGCATCACAAATTTCACAAATAAAGCATTTTTTTCACTGCATTCTAGTTGTGGTT
TGTCCAAACTCATCAATGTATCTTATCATGTCTGGATCAACTGGATAACTCAAGCTA
ACCAAAATCATCCCAAACTTCCCACCCCATACCCTATTACCACTGCCAATTACCTGT
GGTTTCATTTACTCTAAACCTGTGATTCCTCTGAATTATTTTCATTTTAAAGAAATTG
TATTTGTTAAATATGTACTACAAACTTAGTAGT

#### Overview of Experimental Dataset and Categorization

A total of 83 live-cell confocal microscopy experiments (Sep 2025 – May 2026) were performed to characterize the magnetic fluorescence response of mtMagLOV2 across multiple biological conditions. Given the scope of the dataset, experiments were organized into seven categories, each addressing a distinct experimental question related to magnetic field response, mitochondrial membrane potential, fluorescence stability, construct validation, or artifact exclusion. The complete list of experiments is provided in Table S1.

**Category 1: Control Experiments**

The first category comprised cell-free control experiments in which no cells and no mtMagLOV2 construct were present. These experiments were performed using PBS or cell culture medium supplemented with mitochondrial membrane potential dyes (TMRE or MitoTracker Red, 10 nM each) under identical imaging and magnetic field conditions as the experimental groups. The purpose of these controls was to determine whether any observed fluorescence changes could originate from the imaging setup, magnetic field modulation, or the dyes themselves, rather than from a biological response of mtMagLOV2-expressing cells.

Across all control experiments, magnetic field ON/OFF cycling produced no reproducible fluorescence modulation. Only stochastic fluorescence fluctuations consistent with photon shot noise were detected. Addition of FCCP (10 μM) during control experiments did not alter this baseline behavior. These results confirm that the magnetic fluorescence response observed in mtMagLOV2-expressing cells is not attributable to dye photophysics, electromagnetic interference, or instrument artifacts.

**Category 2: HeLa Wild Type Cells**

The second category consisted of experiments using parental HeLa wild type cells, which do not express mtMagLOV2. These experiments served as a cellular negative control to establish that the magnetic fluorescence effect is specifically dependent on mtMagLOV2 expression and is not an intrinsic property of HeLa cells or their mitochondria.

HeLa wild type cells were stained with TMRE (10 nM) and imaged under magnetic field cycling conditions, including ON/OFF intervals of 11 s and 33 s. In selected experiments, FCCP (10 μM) was added to collapse the mitochondrial membrane potential; the expected decrease in TMRE fluorescence was observed in all cases, confirming dye functionality. However, no magnetic fluorescence modulation was detected in HeLa wild type cells under any condition. These results support the conclusion that the magnetic fluorescence effect observed in subsequent categories is attributable specifically to mtMagLOV2.

**Category 3: mtMagLOV2 Characterization in HeLa Cells**

The third category comprised characterization experiments using HeLa mtMagLOV2 cells, a stable HeLa line expressing mtMagLOV2. The primary objective was to establish whether mtMagLOV2 produces a reproducible, field-dependent fluorescence response.

Across a large number of experiments performed under varying magnetic cycling conditions, mtMagLOV2 fluorescence intensity increased reproducibly when the magnetic field was active and decreased upon field removal, tracking the ON/OFF duty cycle of the magnet. To assess reversibility, the magnetic field was switched off for an extended period (approximately 2 min) during ongoing imaging; upon reapplication of the field, the fluorescence response returned to its prior amplitude. This reversible, repeatable behavior confirms that the observed signal is a genuine magnetic response of mtMagLOV2 and not attributable to fluorescence drift or photobleaching.

**Category 4: Drug Perturbation Experiments**

The fourth category investigated the sensitivity of the mtMagLOV2 magnetic fluorescence response to mitochondrial perturbation. Cells were treated with established modulators of the electron transport chain and mitochondrial membrane potential — oligomycin, FCCP, and rotenone — and mtMagLOV2 fluorescence was monitored in parallel with TMRE as an independent indicator of membrane potential.

***Oligomycin (1 μM)***

Oligomycin, an inhibitor of mitochondrial ATP synthase, was tested at multiple concentrations. A concentration of 1 μM was identified as appropriate based on the observed TMRE response, which showed the expected increase in fluorescence consistent with mitochondrial hyperpolarization following ATP synthase inhibition. Parallel mtMagLOV2 measurements showed an initial reduction in mtMagLOV2 fluorescence and magnetic-field modulation following oligomycin treatment, followed by partial recovery toward a stabilized post-treatment response.

***FCCP (10 μM)***

FCCP was used as a mitochondrial uncoupler. Following FCCP addition, TMRE fluorescence decreased markedly in all experiments, confirming mitochondrial membrane potential collapse. mtMagLOV2 fluorescence and magnetic-field modulation were also altered following FCCP treatment. In most traces, the magnetic-field modulation amplitude decreased immediately after FCCP addition and subsequently recovered toward a stabilized post-treatment state. In some experiments, a transient increase in mtMagLOV2 fluorescence was observed immediately after drug addition before the signal returned toward baseline. This variability was likely attributable in part to mechanical perturbation associated with manual drug addition, including brief opening of the microscope enclosure, which may introduce focus instability or temperature fluctuations during acquisition.

***Rotenone (10 µM)***

Rotenone treatment had little effect on the overall mtMagLOV2 fluorescence or on the magnetic field modulation response, consistent with its site of action downstream of the FMN-interacting region of complex I.

***Sequential Drug Experiments***

To recapitulate a Seahorse-style mitochondrial stress test paradigm, experiments were performed in which oligomycin, FCCP, and rotenone were added sequentially during a single imaging session. Drugs were added in the order oligomycin → FCCP → rotenone, with an interval of approximately 7 min between additions to allow signal stabilization before the next perturbation. TMRE fluorescence behaved in accordance with expected mitochondrial physiology: a modest or stable response after oligomycin, a large decrease after FCCP, and sustained low fluorescence after rotenone. Although the precise mtMagLOV2 response pattern varied between experiments — likely due to the manual drug addition artifact described above — one finding was consistent across nearly all drug perturbation experiments regardless of drug identity or protocol: mitochondrial perturbation altered the amplitude of the mtMagLOV2 magnetic fluorescence modulation. This indicates that mtMagLOV2 is not solely a magnetic field reporter but is also sensitive to the functional state of the mitochondria and the integrity of the electron transport chain.

**Category 5: HeLa mtMagLOV2-3F Cells**

The fifth category comprised experiments using HeLa mtMagLOV2-3F cells, a variant of HeLa mtMagLOV2 cells expressing a triple FLAG-tagged mtMagLOV2 construct. These experiments were initiated following a substantial and progressive decline in mtMagLOV2 fluorescence expression observed in the original HeLa mtMagLOV2 cell population over prolonged culture. HeLa mtMagLOV2-3F cells, recovered from cryopreserved stocks, were used as an alternative expressing line.

HeLa mtMagLOV2-3F cells exhibited qualitatively identical magnetic field responses and drug-induced changes in mtMagLOV2 fluorescence as HeLa mtMagLOV2 cells. Notably, the absolute fluorescence intensity of HeLa mtMagLOV2-3F cells was substantially higher than that of HeLa mtMagLOV2, providing improved signal-to-noise for subsequent experiments. These results indicate that the triple FLAG tag does not alter the functional properties of mtMagLOV2 and that the construct remains fully responsive to magnetic stimulation and mitochondrial perturbation.

**Category 6: Puromycin-Selected Cells**

The sixth category consisted of experiments performed following puromycin selection of HeLa mtMagLOV2 and HeLa mtMagLOV2-3F cultures. Puromycin selection was applied to re-enrich for mtMagLOV2-expressing cells after a decline in fluorescence expression was observed in a fraction of the cell population. Following selection, experiments demonstrated magnetic fluorescence responses consistent with those observed in the unselected mtMagLOV2-expressing lines, confirming that puromycin selection successfully maintained the expressing subpopulation.

**Category 7: AC10 Cardiomyocytes**

The seventh category comprised experiments performed in AC10 cardiomyocytes expressing mtMagLOV2-3F. These experiments were conducted to determine whether the mitochondrial magnetic fluorescence response observed in HeLa cells could also be reproduced in a metabolically active cardiac cell model. Confocal imaging demonstrated mitochondrial localization of mtMagLOV2-3F and strong colocalization with TMRE fluorescence in AC10 cardiomyocytes. Under magnetic field cycling conditions, AC10 cells exhibited magnetic-field–dependent fluorescence modulation qualitatively similar to that observed in HeLa cells. Pharmacological perturbation with oligomycin, FCCP, or rotenone altered mtMagLOV2 magnetic-field modulation (ΔF/F)(\Delta F/F)(ΔF/F), with FCCP inducing the expected collapse of TMRE fluorescence and confirming mitochondrial depolarization. Quantitative comparison of the MFE response showed that the magnitude of drug-induced changes differed between AC10 cardiomyocytes and HeLa cells. These findings suggest that the mtMagLOV2 magnetic-field-dependent fluorescence response and its sensitivity to mitochondrial perturbation are not restricted to HeLa cells and can also be observed in cardiomyocytes.

Table S1 summarizes the imaging parameters and experimental conditions used across all live-cell fluorescence measurements in this study. Magnetic field stimulation was applied in alternating ON/OFF cycles and FCCP (10 μM) was added at the indicated time point during continued imaging. All experiments were performed under identical optical settings unless otherwise noted.

**Table S1.** Summary of imaging parameters and experimental conditions used in live-cell fluorescence experiments

| # | Date | Cell used | Label | Staining Condition | Laser Settings | Objective | Detection (nm) | Pixel Time (μs) | Pixel size (nm) | FOV (μm) | Duration (min) | MF, Drug |
|---|---|---|---|---|---|---|---|---|---|---|---|---|
| 1 | 092625(0002) | PBS | TMRE | 10nM; 15 min; DMEM | λex561nm (0.3%)<br>λex488nm (1%) | 63x | 565-630<br>500-550 | 0.85 | 85 | 917×917 | 15 | 22/22 s, OFF, 22/22 s, FCCP |
| 2 | 092625(0010) | HeLa Wild Type | - | - | λex561nm (0.3%)<br>λex488nm (1%) | 63x | 565-630<br>500-550 | 0.85 | 85 | 917×917 | 10 | 22/22 s, OFF |
| 3 | 092725(0002) | HeLa MagLOV R10 | - | - | λex488nm (1%) | 63x | 500-550 | 0.85 | 85 | 917×917 | 10 | Servo is ON without magnet |
| 4 | 092725(0007) | HeLa MagLOV R10 | - | - | λex488nm (1%) | 63x | 565-630<br>500-550 | 0.85 | 85 | 917×917 | 10 | Servo is ON without magnet |
| 5 | 093025(0006) | Cell Culture Media | - | - | λex561nm (0.3%)<br>λex488nm (1%) | 63x | 565-630<br>500-550 | 0.85 | 85 | 917×917 | 10 | 22/22 s, FCCP |
| 6 | 093025(0007) | Cell Culture Media | TMRE | 10nM; 15 min; DMEM | λex561nm (0.3%)<br>λex488nm (1%) | 63x | 565-630<br>500-550 | 0.85 | 85 | 917×917 | 10 | 22/22 s, FCCP |
| 7 | 100125(0015) | HeLa Wild Type | TMRE | 10nM; 15 min; DMEM | λex561nm (0.3%)<br>λex488nm (1%) | 63x | 565-630<br>500-550 | 0.85 | 85 | 917×917 | 15 | Servo is ON without magnet, OFF, FCCP |
| 8 | 100325(0001) | Cell Culture Media | - | - | λex561nm (0.3%)<br>λex488nm (1%) | 63x | 565-630<br>500-550 | 0.85 | 85 | 917×917 | 10 | 11/11 s, OFF, 11/11 s, FCCP |
| 9 | 100325(0002) | Cell Culture Media | MTRed | 10nM; 15 min; DMEM | λex561nm (0.3%)<br>λex488nm (1%) | 63x | 565-630<br>500-550 | 0.85 | 85 | 917×917 | 10 | 11/11 s, OFF, 11/11 s, FCCP |
| 10 | 100725(0006) | HeLa MagLOV R10 | MTRed | 10nM; 15 min; DMEM | λex561nm (0.3%)<br>λex488nm (1%) | 63x | 565-630<br>500-550 | 0.85 | 85 | 917×917 | 30 | 11/11 s, 33/33 s, FCCP |
| 11 | 100725(0009) | HeLa MagLOV R10 | TMRE | 10nM; 15 min; DMEM | λex561nm (0.3%)<br>λex488nm (1%) | 63x | 565-630<br>500-550 | 0.85 | 85 | 917×917 | 30 | 11/11 s, 33/33 s, FCCP |
| 12 | 100725(0012) | HeLa Wild Type | MTRed | 10nM; 15 min; DMEM | λex561nm (0.3%)<br>λex488nm (1%) | 63x | 565-630<br>500-550 | 0.85 | 85 | 917×917 | 30 | 11/11 s, 33/33 s, FCCP |
| 13 | 100725(0015) | HeLa Wild Type | TMRE | 10nM; 15 min; DMEM | λex561nm (0.3%)<br>λex488nm (1%) | 63x | 565-630<br>500-550 | 0.85 | 85 | 917×917 | 30 | 11/11 s, 33/33 s, FCCP |
| 14 | 100725(0018) | HeLa Wild Type | TMRE | 10nM; 15 min; DMEM | λex561nm (0.3%)<br>λex488nm (1%) | 63x | 565-630<br>500-550 | 0.85 | 85 | 917×917 | 30 | 11/11 s, 33/33 s, FCCP |
| 15 | 101625(0003) | HeLa Wild Type | TMRE | 10nM; 15 min; DMEM | λex561nm (0.3%)<br>λex488nm (1%) | 63x | 565-630<br>500-550 | 0.85 | 85 | 917×917 | 30 | 11/11 s, OFF, 33/33 s, 60/60 s |
| 16 | 101625(0018) | HeLa MagLOV R10 | TMRE | 10nM; 15 min; DMEM | λex561nm (0.3%)<br>λex488nm (1%) | 63x | 565-630<br>500-550 | 0.85 | 85 | 917×917 | 15 | 11/11 s |
| 17 | 101625(0021) | HeLa Wild Type | TMRE | 10nM; 15 min; DMEM | λex561nm (0.3%) | 63x | 565-630 | 0.85 | 85 | 917×917 | 30 | 11/11 s |

| | | | | | | | | | | | | |
|---|---|---|---|---|---|---|---|---|---|---|---|---|
| | | | | | λex488nm (1%) | | 500-550 | | | | | |
| 18 | 101725(0003) | HeLa Wild Type | - | - | λex488nm (1%) | 63x | 500-550 | 0.85 | 85 | 917×917 | 15 | 11/11 s, OFF, 11/11 s |
| 19 | 101725(0006) | HeLa Wild Type | - | - | λex488nm (1%) | 63x | 500-550 | 0.85 | 85 | 917×917 | 15 | 11/11 s, OFF, 11/11 s |
| 20 | 101725(0012) | HeLa MagLOV R10 | - | - | λex488nm (1%) | 63x | 500-550 | 0.85 | 85 | 917×917 | 15 | 11/11 s, OFF, 11/11 s |
| 21 | 101725(0015) | HeLa MagLOV R10 | - | - | λex488nm (1%) | 63x | 500-550 | 0.85 | 85 | 917×917 | 15 | 11/11 s, OFF, 11/11 s |
| 22 | 101725(0021) | HeLa MagLOV R10 | TMRE | 10nM; 15 min; DMEM | λex561nm (0.3%) λex488nm (1%) | 63x | 565-630 500-550 | 0.85 | 85 | 917×917 | 20 | 11/11 s, OFF, 33/33 s, 67/67 s |
| 23 | 102125(0044) | HeLa MagLOV R10 | TMRE | 10nM; 15 min; DMEM | λex561nm (0.3%) λex488nm (1%) | 63x | 565-630 500-550 | 0.85 | 85 | 917×917 | 30 | 11/11 s, OFF, 22/22 s, FCCP |
| 24 | 102225(0003) | HeLa MagLOV R10 | TMRE | 10nM; 15 min; DMEM | λex561nm (0.3%) λex488nm (1%) | 63x | 565-630 500-550 | 0.85 | 85 | 917×917 | 30 | 11/11 s, OFF, 22/22 s, FCCP |
| 25 | 102225(0022) | HeLa Wild Type | TMRE | 10nM; 15 min; DMEM | λex561nm (0.3%) λex488nm (1%) | 63x | 565-630 500-550 | 0.85 | 85 | 917×917 | 30 | 11/11 s, OFF, 22/22 s, FCCP |
| 26 | 102225(0025) | HeLa MagLOV R10 | TMRE | 10nM; 15 min; DMEM | λex561nm (0.3%) λex488nm (1%) | 63x | 565-630 500-550 | 0.85 | 85 | 917×917 | 30 | 11/11 s, OFF, 22/22 s, FCCP |
| 27 | 102325(0003) | HeLa MagLOV R10 | TMRE | 10nM; 15 min; DMEM | λex561nm (0.3%) λex488nm (1%) | 63x | 565-630 500-550 | 0.85 | 85 | 917×917 | 30 | 11/11 s, OFF, 22/22 s, FCCP |
| 28 | 102325(0006) | HeLa Wild Type | TMRE | 10nM; 15 min; DMEM | λex561nm (0.3%) λex488nm (1%) | 63x | 565-630 500-550 | 0.85 | 85 | 917×917 | 30 | 11/11 s, OFF, 22/22 s, FCCP |
| 29 | 020326(0003) | HeLa MagLOV R10 | TMRE | 10nM; 15 min; DMEM | λex561nm (0.3%) λex488nm (1%) | 63x | 565-630 500-550 | 0.85 | 85 | 917×917 | 5 | 11/11 s |
| 30 | 020326(0006) | HeLa MagLOV R10 | TMRE | 10nM; 15 min; DMEM | λex561nm (0.3%) λex488nm (1%) | 63x | 565-630 500-550 | 0.85 | 85 | 917×917 | 15 | 22/22 s, OFF, 22/22 s, FCCP |
| 31 | 020426(0004) | HeLa MagLOV R10 | TMRE | 10nM; 15 min; DMEM | λex561nm (0.3%) λex488nm (1%) | 63x | 565-630 500-550 | 0.85 | 85 | 917×917 | 15 | 11/11 s |
| 32 | 020626(0009) | HeLa MagLOV R10 | TMRE | 10nM; 15 min; DMEM | λex561nm (0.3%) λex488nm (1%) | 63x | 565-630 500-550 | 0.85 | 85 | 917×917 | 30 | 11/11 s, OFF, 22/22 s |
| 33 | 020626(0014) | HeLa MagLOV R10 | TMRE | 10nM; 15 min; DMEM | λex561nm (0.3%) λex488nm (1%) | 63x | 565-630 500-550 | 1.17 | 85 | 662×639 | 30 | 11/11 s, OFF, 22/22 s, FCCP |
| 34 | 020926(0007) | HeLa MagLOV R10 | TMRE | 10nM; 15 min; DMEM | λex561nm (0.3%) λex488nm (1%) | 63x | 565-630 500-550 | 1.17 | 85 | 662×639 | 30 | 11/11 s, OFF, 11/11 s, FCCP |
| 35 | 021026(0011) | HeLa MagLOV R10 | TMRE | 10nM; 15 min; DMEM | λex561nm (0.3%) λex488nm (1%) | 63x | 565-630 500-550 | 1.04 | 85 | 749×630 | 30 | 11/11 s, OFF, |

| | | | | | | | | | | | | |
|---|---|---|---|---|---|---|---|---|---|---|---|---|
| | | | | | | | | | | | | 22/22 s, FCCP |
| 36 | 0211260007 | HeLa Wild Type | TMRE | 10nM; 15 min; DMEM | λex561nm (0.3%) λex488nm (1%) | 63x | 565-630 500-550 | 0.85 | 85 | 917×917 | 30 | 11/11 s, OFF, 22/22 s, FCCP |
| 37 | 021226(0003) | HeLa Wild Type | TMRE | 10nM; 15 min; DMEM | λex561nm (0.3%) λex488nm (1%) | 63x | 565-630 500-550 | 0.85 | 85 | 917×917 | 30 | 11/11 s, OFF, 22/22 s, FCCP |
| 38 | 021226(0006) | HeLa MagLOV R10 | TMRE | 10nM; 15 min; DMEM | λex561nm (0.3%) λex488nm (1%) | 63x | 565-630 500-550 | 0.85 | 85 | 917×917 | 30 | 11/11 s, OFF, 22/22 s, FCCP |
| 39 | 021326(0012) | HeLa MagLOV R10-3F | TMRE | 10nM; 15 min; DMEM | λex561nm (0.3%) λex488nm (1%) | 63x | 565-630 500-550 | 0.85 | 85 | 917×917 | 30 | 11/11 s, OFF, 22/22 s, FCCP |
| 40 | 021726(0009) | HeLa MagLOV R10-3F | TMRE | 10nM; 15 min; DMEM | λex561nm (0.3%) λex488nm (1%) | 63x | 565-630 500-550 | 0.85 | 85 | 917×917 | 30 | 11/11 s, OFF, 22/22 s |
| 41 | 021726(0032) | HeLa MagLOV R10-3F | TMRE | 10nM; 15 min; DMEM | λex561nm (0.3%) λex488nm (1%) | 63x | 565-630 500-550 | 0.85 | 85 | 917×917 | 30 | 11/11 s, OFF, 22/22 s, FCCP |
| 42 | 021726(0037) | HeLa MagLOV R10 No cells show MagLov signal | TMRE | 10nM; 15 min; DMEM | λex561nm (0.3%) λex488nm (1%) | 63x | 565-630 500-550 | 0.85 | 85 | 917×917 | 30 | 11/11 s, OFF, 22/22 s, FCCP |
| 43 | 021926(0007) | HeLa MagLOV R10-3F | TMRE | 10nM; 15 min; DMEM | λex561nm (0.3%) λex488nm (1%) | 63x | 565-630 500-550 | 0.85 | 85 | 917×917 | 30 | 11/11 s, OFF, 22/22 s, FCCP |
| 44 | 022026(0003) | HeLa MagLOV R10-3F No cells show MagLov signal | TMRE | 10nM; 15 min; DMEM | λex561nm (0.3%) λex488nm (1%) | 63x | 565-630 500-550 | 0.85 | 85 | 917×917 | 30 | 11/11 s, OFF, 22/22 s, FCCP |
| 45 | 022026(0009) | HeLa MagLOV R10-3F | TMRE | 10nM; 15 min; DMEM | λex561nm (0.3%) λex488nm (1%) | 63x | 565-630 500-550 | 0.85 | 85 | 917×917 | 8 | 11/11 s, OFF |
| 46 | 022026(0020) | HeLa MagLOV R10-3F | TMRE | 10nM; 15 min; DMEM | λex561nm (0.3%) λex488nm (1%) | 63x | 565-630 500-550 | 0.85 | 85 | 917×917 | 30 | 11/11 s, OFF, 22/22 s, FCCP |
| 47 | 022326(0004) | HeLa MagLOV R10+Puromycin | TMRE | 10nM; 15 min; DMEM | λex561nm (0.3%) λex488nm (1%) | 63x | 565-630 500-550 | 0.85 | 85 | 917×917 | 30 | 11/11 s, OFF, 22/22 s, FCCP |
| 48 | 022426(0006) | HeLa MagLOV R10+Puromycin | TMRE | 10nM; 15 min; DMEM | λex561nm (0.3%) λex488nm (1%) | 63x | 565-630 500-550 | 0.85 | 85 | 917×917 | 30 | 11/11 s, OFF, 22/22 s |
| 49 | 022426(0019) | HeLa MagLOV R10-3F+Puromycin | TMRE | 10nM; 15 min; DMEM | λex561nm (0.3%) λex488nm (1%) | 63x | 565-630 500-550 | 0.85 | 85 | 917×917 | 30 | 11/11 s, OFF, 22/22 s, FCCP |
| 50 | 022426(0026) | HeLa MagLOV R10-3F | TMRE | 10nM; 15 min; DMEM | λex561nm (0.3%) λex488nm (1%) | 63x | 565-630 500-550 | 0.85 | 85 | 917×917 | 30 | 11/11 s, OFF, 22/22 s, FCCP |
| 51 | 022626(0003) | HeLa MagLOV R10+Puromycin | TMRE | 10nM; 15 min; DMEM | λex561nm (0.3%) λex488nm (1%) | 63x | 565-630 500-550 | 0.85 | 85 | 917×917 | 30 | 11/11 s, OFF, 11/11 s |
| 52 | 022626(0006) | HeLa MagLOV R10+Puromycin | TMRE | 10nM; 15 min; DMEM | λex561nm (0.3%) λex488nm (1%) | 63x | 565-630 500-550 | 0.85 | 85 | 917×917 | 30 | 11/11 s, OFF, |

| | | | | | | | | | | | | |
|---|---|---|---|---|---|---|---|---|---|---|---|---|
| | | | | | | | | | | | | 11/11 s, FCCP |
| 53 | 022726(0003) | HeLa MagLOV R10+Puromycin | TMRE | 10nM; 15 min; DMEM | λex561nm (0.3%) λex488nm (1%) | 63x | 565-630 500-550 | 0.85 | 85 | 917×917 | 30 | 11/11 s, OFF, 22/22 s, Rotenone 10uM |
| 54 | 030426(0005) | HeLa MagLOV R10-3F | TMRE | 10nM; 15 min; DMEM | λex561nm (0.3%) λex488nm (1%) | 63x | 565-630 500-550 | 0.85 | 85 | 917×917 | 5 | 11/11 s, 20mT magnet |
| 55 | 030426(0012) | HeLa MagLOV R10-3F | TMRE | 10nM; 15 min; DMEM | λex561nm (0.3%) λex488nm (1%) | 63x | 565-630 500-550 | 0.85 | 85 | 917×917 | 5 | 11/11 s |
| 56 | 031126(0008) | HeLa MagLOV R10-3F | TMRE | 10nM; 15 min; DMEM | λex561nm (0.3%) λex488nm (1%) | 63x | 565-630 500-550 | 0.85 | 85 | 917×917 | 30 | 11/11 s, OFF, 22/22 s, Oligomyci n 1uM |
| 57 | 031326(0011) | HeLa MagLOV R10-3F | TMRE | 10nM; 15 min; DMEM | λex561nm (0.3%) λex488nm (1%) | 63x | 565-630 500-550 | 0.85 | 85 | 917×917 | 30 | 11/11 s, OFF, 22/22 s, Oligomyci n 10uM |
| 58 | 031326(0016) | HeLa MagLOV R10-3F | TMRE | 10nM; 15 min; DMEM | λex561nm (0.3%) λex488nm (1%) | 63x | 565-630 500-550 | 0.85 | 85 | 917×917 | 30 | 11/11 s, OFF, 22/22 s, FCCP |
| 59 | 031326(0022) | HeLa MagLOV R10-3F | TMRE | 10nM; 15 min; DMEM | λex561nm (0.3%) λex488nm (1%) | 63x | 565-630 500-550 | 0.85 | 85 | 917×917 | 30 | 11/11 s, OFF, 22/22 s, FCCP |
| 60 | 041326(0007) | HeLa MagLOV R10-3F | TMRE | 10nM; 15 min; DMEM | λex561nm (0.3%) λex488nm (1%) | 63x | 565-630 500-550 | 0.85 | 85 | 917×917 | 10 | 11/11 s, FCCP |
| 61 | 041326(0010) | HeLa MagLOV R10-3F | TMRE | 10nM; 15 min; DMEM | λex561nm (0.3%) λex488nm (1%) | 63x | 565-630 500-550 | 0.85 | 85 | 917×917 | 10 | 11/11 s, FCCP |
| 62 | 041326(0013) | HeLa MagLOV R10+Puromycin | TMRE | 10nM; 15 min; DMEM | λex561nm (0.3%) λex488nm (1%) | 63x | 565-630 500-550 | 0.85 | 85 | 917×917 | 10 | 11/11 s, FCCP |
| 63 | 041426(0003) | HeLa MagLOV R10+Puromycin | TMRE | 10nM; 15 min; DMEM | λex561nm (0.3%) λex488nm (1%) | 63x | 565-630 500-550 | 0.85 | 85 | 917×917 | 27 | 11/11 s, Oligomyci n1uM, FCCP10u M, Rotenone 10uM |
| 64 | 041726(0003) | HeLa MagLOV R10+Puromycin | TMRE | 10nM; 15 min; DMEM | λex561nm (0.3%) λex488nm (1%) | 63x | 565-630 500-550 | 0.85 | 85 | 917×917 | 25 | 11/11 s, FCCP |
| 65 | 041726(0007) | HeLa MagLOV R10+Puromycin | TMRE | 10nM; 15 min; DMEM | λex561nm (0.3%) λex488nm (1%) | 63x | 565-630 500-550 | 0.85 | 85 | 917×917 | 25 | 11/11 s, FCCP 5uM |
| 66 | 041726(0010) | HeLa MagLOV R10+Puromycin | TMRE | 10nM; 15 min; DMEM | λex561nm (0.3%) λex488nm (1%) | 63x | 565-630 500-550 | 0.85 | 85 | 917×917 | 25 | 11/11 s, FCCP |
| 67 | 041726(0016) | HeLa MagLOV R10+Puromycin | TMRE | 10nM; 15 min; DMEM | λex561nm (0.3%) λex488nm (1%) | 63x | 565-630 500-550 | 0.85 | 85 | 917×917 | 25 | 11/11 s, FCCP 5uM |
| 68 | 042126(0003) | HeLa MagLOV R10+Puromycin | - | - | λex488nm (1%) | 63x | 500-550 | 0.85 | 85 | 917×917 | 20 | 11/11 s |
| 69 | 042126(0006) | HeLa MagLOV R10+Puromycin | - | - | λex561nm (0.3%) λex488nm (1%) | 63x | 565-630 500-550 | 0.85 | 85 | 917×917 | 20 | 11/11 s |
| 70 | 042126(0009) | HeLa MagLOV R10+Puromycin | - | - | λex488nm (1%) | 63x | 500-550 | 0.8 | 59 | 1319×13 19 | 20 | 11/11 s |
| 71 | 042126(0012) | HeLa MagLOV R10+Puromycin | - | - | λex488nm (1%) | 63x | 500-550 | 0.85 | 85 | 917×917 | 20 | FCCP, 11/11 s |

| | | | | | | | | | | | | |
|---|---|---|---|---|---|---|---|---|---|---|---|---|
| 72 | 042126(0015) | HeLa MagLOV R10+Puromycin | - | - | λex488nm (1%) | 63x | 500-550 | 0.85 | 85 | 917×917 | 25 | 11/11 s, FCCP |
| 73 | 042826(0007) | HeLa MagLOV R10+Puromycin | TMRE | 10nM; 15 min; DMEM | λex561nm (0.3%) λex488nm (1%) | 63x | 565-630 500-550 | 0.85 | 85 | 917×917 | 30 | 11/11 s, Oligomyci n1uM, FCCP 10uM, Rotenone 10uM |
| 74 | 042826(0015) | HeLa MagLOV R10+Puromycin | TMRE | 10nM; 15 min; DMEM | λex561nm (0.3%) λex488nm (1%) | 63x | 565-630 500-550 | 0.85 | 85 | 917×917 | 30 | 11/11 s, Oligomyci n1uM, FCCP 10uM, Rotenone 10uM |
| 75 | 050426(0009) | AC10-R10-3F | TMRE | 10nM;30 min; DMEM | λex561nm (0.3%) λex488nm (1%) | 63x | 565-630 500-550 | 0.85 | 85 | 917×917 | 30 | 11/11 s, FCCP 10uM |
| 76 | 050426(0012) | HeLa MagLOV R10+Puromycin | TMRE | 10nM; 15 min; DMEM | λex561nm (0.3%) λex488nm (1%) | 63x | 565-630 500-550 | 0.85 | 85 | 917×917 | 16 | 11/11 s, FCCP 3uM |
| 77 | 050426(0015) | HeLa MagLOV R10+Puromycin | TMRE | 10nM; 15 min; DMEM | λex561nm (0.3%) λex488nm (1%) | 63x | 565-630 500-550 | 0.85 | 85 | 917×917 | 16 | 11/11 s, media |
| 78 | 052526(0005) | HeLa MagLOV R10+Puromycin | TMRE | 10nM; 15 min; DMEM | λex561nm (0.3%) λex488nm (1%) | 63x | 565-630 500-550 | 0.85 | 85 | 917×917 | 5 | 11/11 s |
| 79 | 052526(0007) | HeLa MagLOV R10+Puromycin | TMRE | 10nM; 15 min; DMEM | λex561nm (0.3%) λex488nm (1%) | 63x | 565-630 500-550 | 0.85 | 85 | 917×917 | 5 | 11/11 s |
| 80 | 052526(0015) | AC10-R10-3F | TMRE | 10nM;30 min; DMEM | λex561nm (0.3%) λex488nm (1%) | 63x | 565-630 500-550 | 0.85 | 85 | 917×917 | 35 | 11/11 s, Oligomyci n1uM, FCCP 10uM, Rotenone 10uM |
| 81 | 052726(0011) | AC10-R10-3F | TMRE | 10nM;30 min; DMEM | λex561nm (0.3%) λex488nm (1%) | 63x | 565-630 500-550 | 0.85 | 85 | 917×917 | 30 | 11/11 s, Oligomyci n1uM, FCCP 10uM, Rotenone 10uM |
| 82 | 052726(0030) | AC10-R10-3F | TMRE | 10nM;30 min; DMEM | λex561nm (0.3%) λex488nm (1%) | 63x | 565-630 500-550 | 0.85 | 85 | 917×917 | 30 | 11/11 s, Oligomyci n1uM, FCCP 10uM, Rotenone 10uM |
| 83 | 052826(0003) | AC10-R10-3F | TMRE | 10nM;30 min; DMEM | λex561nm (0.3%) λex488nm (1%) | 63x | 565-630 500-550 | 0.85 | 85 | 917×917 | 30 | 11/11 s, Oligomyci n1uM, FCCP 10uM, Rotenone 10uM |

Table S2 lists all fluorescence imaging datasets acquired in this study, including HeLa wild type and MagLOV-expressing cells stained with TMRE or MitoTracker Red. All images were acquired under comparable microscope settings to ensure consistency across experiments.

**Table S2.** List of fluorescence imaging datasets and acquisition parameters.

| # | Date | Cell used | Label | Staining Condition | Laser Settings | Objective | Detection (nm) | Pixel Time (µs) | Pixel size (nm) | FOV (µm) |
|---|---|---|---|---|---|---|---|---|---|---|
| 1 | 102325(0001) | HeLa MagLOV | TMRE | 10nM; 15 min; DMEM | λex561nm (0.3%)<br>λex488nm (1%) | 63x | 565-630<br>500-550 | 1.01 | 102 | 768×768 |
| 2 | 102325(0002) | HeLa MagLOV | TMRE | 10nM; 15 min; DMEM | λex561nm (0.3%)<br>λex488nm (1%) | 63x | 565-630<br>500-550 | 0.85 | 85 | 917×917 |
| 3 | 102325(0004) | HeLa Wild Type | TMRE | 10nM; 15 min; DMEM | λex561nm (0.3%)<br>λex488nm (1%) | 63x | 565-630<br>500-550 | 1.01 | 102 | 768×768 |
| 4 | 102325(0005) | HeLa Wild Type | TMRE | 10nM; 15 min; DMEM | λex561nm (0.3%)<br>λex488nm (1%) | 63x | 565-630<br>500-550 | 0.85 | 85 | 917×917 |
| 5 | 102225(0001, 0004) | HeLa MagLOV | TMRE | 10nM; 15 min; DMEM | λex561nm (0.3%)<br>λex488nm (1%) | 63x | 565-630<br>500-550 | 1.01 | 102 | 768×768 |
| 6 | 102225(0002, 0005) | HeLa Ma gLOV | TMRE | 10nM; 15 min; DMEM | λex561nm (0.3%)<br>λex488nm (1%) | 63x | 565-630<br>500-550 | 0.85 | 85 | 917×917 |
| 7 | 102225(0007-0008) | HeLa MagLOV | TMRE | 10nM; 15 min; DMEM | λex561nm (0.3%)<br>λex488nm (1%) | 63x | 565-630<br>500-550 | 1.01 | 102 | 768×768 |
| 8 | 102225(0009-0016) | HeLa MagLOV | TMRE | 10nM; 15 min; DMEM | λex561nm (0.3%)<br>λex488nm (1%) | 63x | 565-630<br>500-550 | 1.01 | 39 | 768×524 |
| 9 | 102225(0017-0019) | HeLa MagLOV | TMRE | 10nM; 15 min; DMEM | λex561nm (0.3%)<br>λex488nm (1%) | 63x | 565-630<br>500-550 | 0.85 | 51 | 512×424 |
| 10 | 102225(0020) | HeLa Wild Type | TMRE | 10nM; 15 min; DMEM | λex561nm (0.3%)<br>λex488nm (1%) | 63x | 565-630<br>500-550 | 1.01 | 102 | 768×768 |
| 11 | 102225(0021) | HeLa Wild Type | TMRE | 10nM; 15 min; DMEM | λex561nm (0.3%)<br>λex488nm (1%) | 63x | 565-630<br>500-550 | 0.85 | 85 | 917×917 |
| 12 | 102225(0023, 0024) | HeLa MagLOV | TMRE | 10nM; 15 min; DMEM | λex561nm (0.3%)<br>λex488nm (1%) | 63x | 565-630<br>500-550 | 1.01 | 102 | 768×768 |
| 13 | 102125(0001,0005-0007, 0016, 0020, 0024, 0025, 0032, 0033, 0042) | HeLa MagLOV | TMRE | 10nM; 15 min; DMEM | λex561nm (0.3%)<br>λex488nm (1%) | 63x | 565-630<br>500-550 | 1.01 | 102 | 768×768 |
| 14 | 102125(0002-0004) | HeLa MagLOV | TMRE | 10nM; 15 min; DMEM | λex561nm (0.3%)<br>λex488nm (1%) | 63x | 565-630<br>500-550 | 1.01 | 24 | 768×768 |
| 15 | 102125(0008) | HeLa MagLOV | TMRE | 10nM; 15 min; DMEM | λex561nm (0.3%)<br>λex488nm (1%) | 63x | 565-630<br>500-550 | 1.01 | 97 | 768×404 |
| 16 | 102125(0009) | HeLa MagLOV | TMRE | 10nM; 15 min; DMEM | λex561nm (0.3%)<br>λex488nm (1%) | 63x | 565-630<br>500-550 | 1.01 | 46 | 768×768 |
| 17 | 102125(0010,0011) | HeLa MagLOV | TMRE | 10nM; 15 min; DMEM | λex561nm (0.3%)<br>λex488nm (1%) | 63x | 565-630<br>500-550 | 1.01 | 23 | 768×768 |
| 18 | 102125(0012-0014) | HeLa MagLOV | TMRE | 10nM; 15 min; DMEM | λex561nm (0.3%)<br>λex488nm (1%) | 63x | 565-630<br>500-550 | 0.85 | 76 | 512×370 |
| 19 | 102125(0015) | HeLa MagLOV | TMRE | 10nM; 15 min; DMEM | λex561nm (0.3%)<br>λex488nm (1%) | 63x | 565-630<br>500-550 | 0.85 | 57 | 512×512 |

| | | | | | | | | | | |
|---|---|---|---|---|---|---|---|---|---|---|
| 20 | 102125(0017-0019) | HeLa MagLOV | TMRE | 10nM; 15 min; DMEM | λex561nm (0.3%)<br>λex488nm (1%) | 63x | 565-630<br>500-550 | 1.01 | 51 | 768×768 |
| 21 | 102125(0021) | HeLa MagLOV | TMRE | 10nM; 15 min; DMEM | λex561nm (0.3%)<br>λex488nm (1%) | 63x | 565-630<br>500-550 | 1.01 | 90 | 768×426 |
| 22 | 102125(0022, 0023) | HeLa MagLOV | TMRE | 10nM; 15 min; DMEM | λex561nm (0.3%)<br>λex488nm (1%) | 63x | 565-630<br>500-550 | 1.01 | 38 | 768×768 |
| 23 | 102125(0026) | HeLa MagLOV | TMRE | 10nM; 15 min; DMEM | λex561nm (0.3%)<br>λex488nm (1%) | 63x | 565-630<br>500-550 | 1.01 | 99 | 768×354 |
| 24 | 102125(0027) | HeLa MagLOV | TMRE | 10nM; 15 min; DMEM | λex561nm (0.3%)<br>λex488nm (1%) | 63x | 565-630<br>500-550 | 0.85 | 141 | 512×269 |
| 25 | 102125(0028-0031, 0035-0039) | HeLa MagLOV | TMRE | 10nM; 15 min; DMEM | λex561nm (0.3%)<br>λex488nm (1%) | 63x | 565-630<br>500-550 | 1.01 | 42 | 768×768 |
| 26 | 102125(0034) | HeLa MagLOV | TMRE | 10nM; 15 min; DMEM | λex561nm (0.3%)<br>λex488nm (1%) | 63x | 565-630<br>500-550 | 1.12 | 42 | 692×768 |
| 27 | 102125(0040) | HeLa MagLOV | TMRE | 10nM; 15 min; DMEM | λex561nm (0.3%)<br>λex488nm (1%) | 63x | 565-630<br>500-550 | 0.85 | 125 | 512×201 |
| 28 | 102125(0041) | HeLa MagLOV | TMRE | 10nM; 15 min; DMEM | λex561nm (0.3%)<br>λex488nm (1%) | 63x | 565-630<br>500-550 | 1.01 | 84 | 768×768 |
| 29 | 102125(0043) | HeLa MagLOV | TMRE | 10nM; 15 min; DMEM | λex561nm (0.3%)<br>λex488nm (1%) | 63x | 565-630<br>500-550 | 0.85 | 85 | 917×917 |
| 30 | 101725(0001, 0004) | HeLa Wild Type | - | - | λex488nm (1%) | 63x | 500-550 | 1.01 | 102 | 768×768 |
| 31 | 101725(0002, 0005) | HeLa Wild Type | - | - | λex488nm (1%) | 63x | 500-550 | 0.85 | 85 | 917×917 |
| 32 | 101725(0007) | HeLa Wild Type | TMRE | 10nM; 15 min; DMEM | λex561nm (0.3%)<br>λex488nm (1%) | 63x | 565-630<br>500-550 | 1.01 | 102 | 768×768 |
| 33 | 101725(0008) | HeLa Wild Type | TMRE | 10nM; 15 min; DMEM | λex561nm (0.3%)<br>λex488nm (1%) | 63x | 565-630<br>500-550 | 0.85 | 85 | 917×917 |
| 34 | 101725(0010, 0013) | HeLa MagLOV | - | - | λex488nm (1%) | 63x | 500-550 | 1.01 | 102 | 768×768 |
| 35 | 101725(0011, 0014) | HeLa MagLOV | - | - | λex488nm (1%) | 63x | 500-550 | 0.85 | 85 | 917×917 |
| 36 | 101725(0016) | HeLa Wild Type | TMRE | 10nM; 15 min; DMEM | λex561nm (0.3%)<br>λex488nm (1%) | 63x | 565-630<br>500-550 | 1.01 | 102 | 768×768 |
| 37 | 101725(0017) | HeLa Wild Type | TMRE | 10nM; 15 min; DMEM | λex561nm (0.3%)<br>λex488nm (1%) | 63x | 565-630<br>500-550 | 0.85 | 85 | 917×917 |
| 38 | 101725(0019, 0022) | HeLa MagLOV | TMRE | 10nM; 15 min; DMEM | λex561nm (0.3%)<br>λex488nm (1%) | 63x | 565-630<br>500-550 | 1.01 | 102 | 768×768 |
| 39 | 101725(0020, 0023) | HeLa MagLOV | TMRE | 10nM; 15 min; DMEM | λex561nm (0.3%)<br>λex488nm (1%) | 63x | 565-630<br>500-550 | 0.85 | 85 | 917×917 |
| 40 | 101625(0001, 0004) | HeLa Wild Type | TMRE | 10nM; 15 min; DMEM | λex561nm (0.3%)<br>λex488nm (1%) | 63x | 565-630<br>500-550 | 1.01 | 102 | 768×768 |
| 41 | 101625(0002, 0005) | HeLa Wild Type | TMRE | 10nM; 15 min; DMEM | λex561nm (0.3%)<br>λex488nm (1%) | 63x | 565-630<br>500-550 | 0.85 | 85 | 917×917 |
| 42 | 101625(0007, 0010, 0013, 0016) | HeLa MagLOV | TMRE | 10nM; 15 min; DMEM | λex561nm (0.3%)<br>λex488nm (1%) | 63x | 565-630<br>500-550 | 1.01 | 102 | 768×768 |
| 43 | 101625(0008, 0011, 0014, 0017) | HeLa MagLOV | TMRE | 10nM; 15 min; DMEM | λex561nm (0.3%)<br>λex488nm (1%) | 63x | 565-630<br>500-550 | 0.85 | 85 | 917×917 |
| 44 | 101625(0019) | HeLa Wild Type | TMRE | 10nM; 15 min; DMEM | λex561nm (0.3%)<br>λex488nm (1%) | 63x | 565-630<br>500-550 | 1.01 | 102 | 768×768 |
| 45 | 101625(0020) | HeLa Wild Type | TMRE | 10nM; 15 min; DMEM | λex561nm (0.3%)<br>λex488nm (1%) | 63x | 565-630<br>500-550 | 0.85 | 85 | 917×917 |
| 46 | 101525(0001, 0004, 0007) | HeLa MagLOV | TMRE | 10nM; 15 min; DMEM | λex561nm (0.3%)<br>λex488nm (1%) | 63x | 565-630<br>500-550 | 1.01 | 102 | 768×768 |

| | | | | | | | | | | |
|---|---|---|---|---|---|---|---|---|---|---|
| 47 | 101525(0002, 0005, 0008) | HeLa MagLOV | TMRE | 10nM; 15 min; DMEM | λex561nm (0.3%)<br>λex488nm (1%) | 63x | 565-630<br>500-550 | 0.85 | 85 | 917×917 |
| 48 | 101425(0001) | HeLa MagLOV | TMRE | 10nM; 15 min; DMEM | λex561nm (0.3%)<br>λex488nm (1%) | 63x | 565-630<br>500-550 | 1.01 | 102 | 768×768 |
| 49 | 101425(0002) | HeLa MagLOV | TMRE | 10nM; 15 min; DMEM | λex561nm (0.3%)<br>λex488nm (1%) | 63x | 565-630<br>500-550 | 0.85 | 85 | 917×917 |
| 50 | 101425(0004, 0007) | HeLa MagLOV | - | - | λex488nm (1%) | 63x | 500-550 | 1.01 | 102 | 768×768 |
| 51 | 101425(0005, 0008) | HeLa MagLOV | - | - | λex488nm (1%) | 63x | 500-550 | 0.85 | 85 | 917×917 |
| 52 | 101325(0001) | HeLa MagLOV | TMRE | 10nM; 15 min; DMEM | λex561nm (0.3%)<br>λex488nm (1%) | 63x | 565-630<br>500-550 | 1.01 | 102 | 768×768 |
| 53 | 101325(0002) | HeLa MagLOV | TMRE | 10nM; 15 min; DMEM | λex561nm (0.3%)<br>λex488nm (1%) | 63x | 565-630<br>500-550 | 0.85 | 85 | 917×917 |
| 54 | 100725(0001, 0004) | HeLa MagLOV | MTRed | 10nM; 15 min; DMEM | λex561nm (0.3%)<br>λex488nm (1%) | 63x | 565-630<br>500-550 | 1.01 | 102 | 768×768 |
| 55 | 100725(0002, 0005) | HeLa MagLOV | MTRed | 10nM; 15 min; DMEM | λex561nm (0.3%)<br>λex488nm (1%) | 63x | 565-630<br>500-550 | 0.85 | 85 | 917×917 |
| 56 | 100725(0007) | HeLa MagLOV | TMRE | 10nM; 15 min; DMEM | λex561nm (0.3%)<br>λex488nm (1%) | 63x | 565-630<br>500-550 | 1.01 | 102 | 768×768 |
| 57 | 100725(0008) | HeLa MagLOV | TMRE | 10nM; 15 min; DMEM | λex561nm (0.3%)<br>λex488nm (1%) | 63x | 565-630<br>500-550 | 0.85 | 85 | 917×917 |
| 58 | 100725(0010) | HeLa Wild Type | MTRed | 10nM; 15 min; DMEM | λex561nm (0.3%)<br>λex488nm (1%) | 63x | 565-630<br>500-550 | 1.01 | 102 | 768×768 |
| 59 | 100725(0011) | HeLa Wild Type | MTRed | 10nM; 15 min; DMEM | λex561nm (0.3%)<br>λex488nm (1%) | 63x | 565-630<br>500-550 | 0.85 | 85 | 917×917 |
| 60 | 100725(0013, 0016) | HeLa Wild Type | TMRE | 10nM; 15 min; DMEM | λex561nm (0.3%)<br>λex488nm (1%) | 63x | 565-630<br>500-550 | 1.01 | 102 | 768×768 |
| 61 | 100725(0014, 0017) | HeLa Wild Type | TMRE | 10nM; 15 min; DMEM | λex561nm (0.3%)<br>λex488nm (1%) | 63x | 565-630<br>500-550 | 0.85 | 85 | 917×917 |
| 62 | 100325(0003) | HeLa MagLOV | TMRE | 10nM; 15 min; DMEM | λex561nm (0.3%)<br>λex488nm (1%) | 63x | 565-630<br>500-550 | 1.01 | 102 | 768×768 |
| 63 | 100325(0004) | HeLa MagLOV | TMRE | 10nM; 15 min; DMEM | λex561nm (0.3%)<br>λex488nm (1%) | 63x | 565-630<br>500-550 | 0.85 | 85 | 917×917 |
| 64 | 100225(0001, 0004) | HeLa Wild Type | MTRed | 10nM; 15 min; DMEM | λex561nm (0.3%)<br>λex488nm (1%) | 63x | 565-630<br>500-550 | 1.01 | 102 | 768×768 |
| 65 | 100225(0002, 0005) | HeLa Wild Type | MTRed | 10nM; 15 min; DMEM | λex561nm (0.3%)<br>λex488nm (1%) | 63x | 565-630<br>500-550 | 0.85 | 85 | 917×917 |
| 66 | 100225(0007, 0010) | HeLa MagLOV | MTRed | 10nM; 15 min; DMEM | λex561nm (0.3%)<br>λex488nm (1%) | 63x | 565-630<br>500-550 | 1.01 | 102 | 768×768 |
| 67 | 100225(0008, 0011) | HeLa MagLOV | MTRed | 10nM; 15 min; DMEM | λex561nm (0.3%)<br>λex488nm (1%) | 63x | 565-630<br>500-550 | 0.85 | 85 | 917×917 |
| 68 | 100125(0010) | HeLa MagLOV | TMRE | 10nM; 15 min; DMEM | λex561nm (0.3%)<br>λex488nm (1%) | 63x | 565-630<br>500-550 | 1.01 | 102 | 768×768 |
| 69 | 100125(0011) | HeLa MagLOV | TMRE | 10nM; 15 min; DMEM | λex561nm (0.3%)<br>λex488nm (1%) | 63x | 565-630<br>500-550 | 0.85 | 85 | 917×917 |
| 70 | 100125(0013) | HeLa Wild Type | TMRE | 10nM; 15 min; DMEM | λex561nm (0.3%)<br>λex488nm (1%) | 63x | 565-630<br>500-550 | 1.01 | 102 | 768×768 |
| 71 | 100125(0014) | HeLa Wild Type | TMRE | 10nM; 15 min; DMEM | λex561nm (0.3%)<br>λex488nm (1%) | 63x | 565-630<br>500-550 | 0.85 | 85 | 917×917 |
| 72 | 100125(0016) | HeLa Wild Type | MTRed | 10nM; 15 min; DMEM | λex561nm (0.3%)<br>λex488nm (1%) | 63x | 565-630<br>500-550 | 1.01 | 102 | 768×768 |

| | | | | | | | | | | |
|---|---|---|---|---|---|---|---|---|---|---|
| 73 | 100125(0017) | HeLa Wild Type | MTRed | 10nM; 15 min; DMEM | λex561nm (0.3%)<br>λex488nm (1%) | 63x | 565-630<br>500-550 | 0.85 | 85 | 917×917 |
| 74 | 093025(0001) | HeLa MagLOV | - | - | λex561nm (0.3%)<br>λex488nm (1%) | 63x | 565-630<br>500-550 | 1.01 | 102 | 768×768 |
| 75 | 093025(0002) | HeLa MagLOV | - | - | λex561nm (0.3%)<br>λex488nm (1%) | 63x | 565-630<br>500-550 | 0.85 | 85 | 917×917 |
| 76 | 093025(0004, 0008) | HeLa MagLOV | TMRE | 10nM; 15 min; DMEM | λex561nm (0.3%)<br>λex488nm (1%) | 63x | 565-630<br>500-550 | 1.01 | 102 | 768×768 |
| 77 | 093025(0009) | HeLa MagLOV | TMRE | 10nM; 15 min; DMEM | λex561nm (0.3%)<br>λex488nm (1%) | 63x | 565-630<br>500-550 | 0.85 | 85 | 917×917 |
| 78 | 093025(0011) | HeLa Wild Type | TMRE | 10nM; 15 min; DMEM | λex561nm (0.3%)<br>λex488nm (1%) | 63x | 565-630<br>500-550 | 1.01 | 102 | 768×768 |
| 79 | 093025(0012) | HeLa Wild Type | TMRE | 10nM; 15 min; DMEM | λex561nm (0.3%)<br>λex488nm (1%) | 63x | 565-630<br>500-550 | 0.85 | 85 | 917×917 |
| 80 | 093025(0018, 0021) | HeLa MagLOV | TMRE | 10nM; 15 min; DMEM | λex561nm (0.3%)<br>λex488nm (1%) | 63x | 565-630<br>500-550 | 1.01 | 102 | 768×768 |
| 81 | 093025(0020, 0023) | HeLa MagLOV | TMRE | 10nM; 15 min; DMEM | λex488nm (1%) | 63x | 500-550 | 1.01 | 102 | 768×768 |
| 82 | 092725(0001, 0004, 0006) | HeLa MagLOV | - | - | λex488nm (1%) | 63x | 500-550 | 1.01 | 102 | 768×768 |
| 83 | 092725(0003) | HeLa MagLOV | - | - | λex488nm (1%) | 63x | 500-550 | 0.85 | 85 | 917×917 |
| 84 | 092625(0009) | HeLa Wild Type | - | - | λex561nm (0.3%)<br>λex488nm (1%) | 63x | 565-630<br>500-550 | 1.01 | 102 | 768×768 |
| 85 | 092525(0001) | HeLa Wild Type | TMRE | 10nM; 15 min; DMEM | λex561nm (0.3%)<br>λex488nm (1%) | 63x | 565-630<br>500-550 | 0.85 | 85 | 917×917 |
| 86 | 092325(0001) | HeLa MagLOV | TMRE | 10nM; 15 min; DMEM | λex561nm (0.3%)<br>λex488nm (1%) | 63x | 565-630<br>500-550 | 1.01 | 102 | 768×768 |
| 87 | 092325(0002) | HeLa MagLOV | TMRE | 10nM; 15 min; DMEM | λex561nm (0.3%)<br>λex488nm (1%) | 63x | 565-630<br>500-550 | 0.85 | 85 | 917×917 |
| 88 | 092325(0005) | HeLa MagLOV | TMRE | 10nM; 15 min; DMEM | λex488nm (1%) | 63x | 500-550 | 1.01 | 102 | 768×768 |
| 89 | 092225(0001, 0004, 0007, 0010, 0013, 0015) | HeLa MagLOV | - | - | λex488nm (1%) | 63x | 500-550 | 1.01 | 102 | 768×768 |
| 90 | 092225(0002, 0005, 0008, 0011, 0014, 0016) | HeLa MagLOV | - | - | λex488nm (1%) | 63x | 500-550 | 0.85 | 85 | 917×917 |

#### Endogenous vs. MagLOV fluorescence

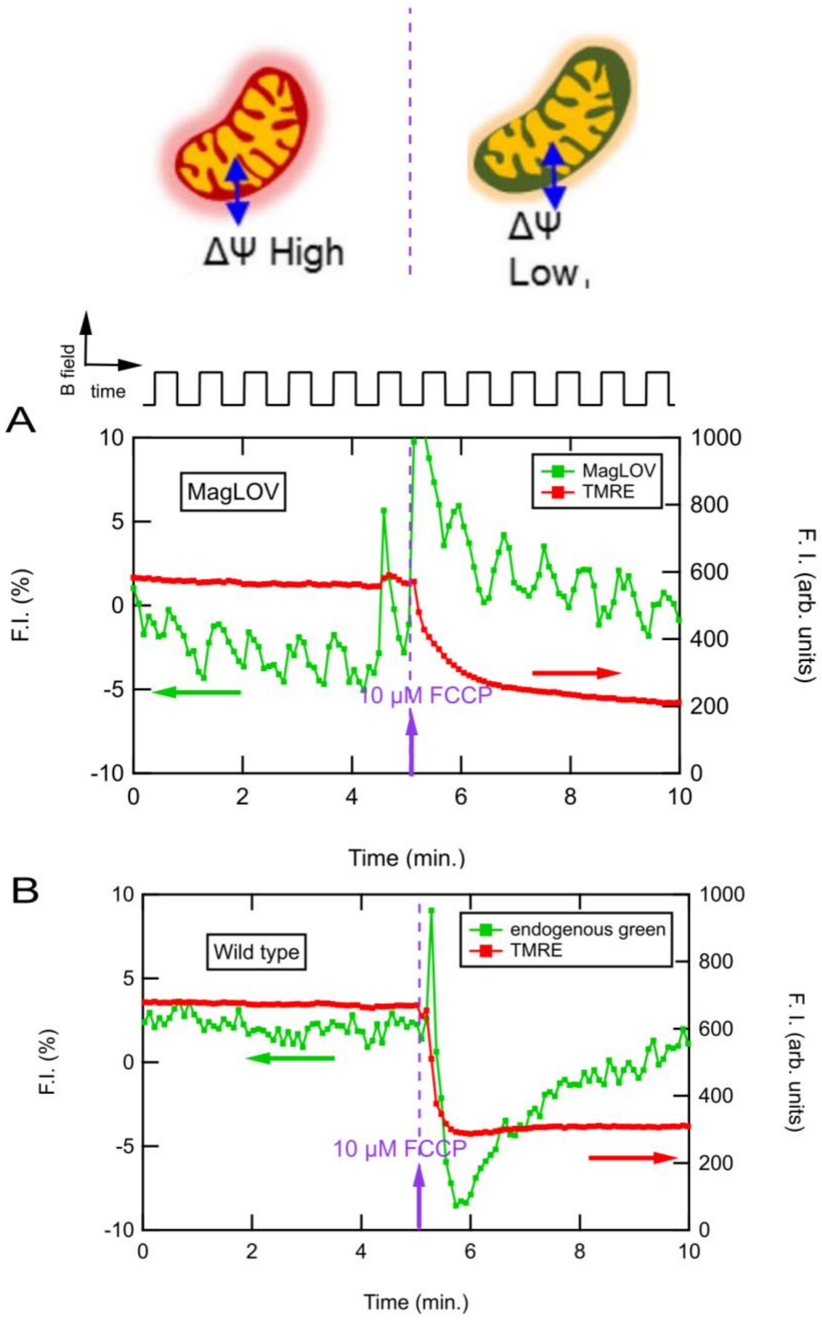


**Figure S12. Effect of FCCP on magnetic-field response.** (A) In mtMagLOV2-expressing cells, addition of FCCP induces a rapid drop in TMRE fluorescence (shown schematically at top of the Figure S14), confirming complete mitochondrial depolarization. Despite this, mtMagLOV2 fluorescence retains its characteristic ~10% magnetic-field modulation (shown schematically as a square wave above the Figures), indicating that mtMagLOV2 remains functional as a quantum sensor after loss of membrane potential. (B) Wild type cells show the expected FCCP-induced TMRE collapse but exhibit no measurable magnetic-field–dependent fluorescence; endogenous flavin autofluorescence is substantially lower than mtMagLOV2 fluorescence and its magnetic-field response is below the noise threshold.

#### Supplementary MATLAB code for MFE quantification

The following MATLAB code was used to quantify MFE from fluorescence time-series data exported as CSV files.

```
clear; clc; close all;
%% -------- 1. Load CSV --------
[fname, fpath] = uigetfile('*.csv', 'Select your CSV file');
if isequal(fname, 0)
    disp('No file selected. Exiting.');
    return;
end
[~, fname_base, ~] = fileparts(fname);
data = readtable(fullfile(fpath, fname), 'VariableNamingRule', 'preserve');
TimeW  = data{:,1};
TMRE   = data{:,2};
MagLOV = data{:,3};
fprintf('Loaded %d rows | Time: %.2f - %.2f min\n', ...
        height(data), min(TimeW), max(TimeW));
%% -------- 2. Find upper and lower fluorescence envelopes --------
[~, idx_top] = findpeaks( MagLOV, 'MinPeakDistance', 1);
[~, idx_bot] = findpeaks(-MagLOV, 'MinPeakDistance', 1);
fprintf('Maxima found: %d\n', length(idx_top));
fprintf('Minima found: %d\n', length(idx_bot));
env_top = interp1(TimeW(idx_top), MagLOV(idx_top), TimeW, 'spline', 'extrap');
env_bot = interp1(TimeW(idx_bot), MagLOV(idx_bot), TimeW, 'spline', 'extrap');
% Continuous magnetic-field modulation from the envelope.
% This is ΔF/F, not the final MFE change.
dFF_cont = (env_top - env_bot) ./ env_bot * 100;
%% -------- 3. Figure for manual window selection --------
fig1 = figure('Name', 'Click to define windows', ...
              'Position', [80 200 1000 450], ...
              'Visible', 'on');
plot(TimeW, MagLOV, 'g-', 'LineWidth', 1.0, 'DisplayName', 'MagLOV');
hold on;
yyaxis right
plot(TimeW, TMRE, 'r-', 'LineWidth', 1.5, 'DisplayName', 'TMRE');
```

```
yyaxis left
xlabel('Time (min.)', 'FontSize', 11);
title(sprintf('Click 4 times: [pre start] [injection] [stable] [post end] | %s', fname), ...
    'Interpreter', 'none', 'FontSize', 10);
legend('Location', 'best', 'FontSize', 9);
set(gca, 'FontSize', 10);
grid on; box on;
%% -------- 4. Four clicks --------
disp('>>> Click 1: start of pre-treatment window');
[t_pre_start, ~] = ginput(1);
fprintf('Pre start:   %.2f min\n', t_pre_start);
yyaxis left
xline(t_pre_start, '--', 'Color', [0.2 0.6 0.2], 'LineWidth', 1.8, ...
    'Label', sprintf('Pre: %.2f', t_pre_start), ...
    'LabelVerticalAlignment', 'top');
drawnow;
disp('>>> Click 2: injection time / start of dead zone');
[inj_time, ~] = ginput(1);
fprintf('Injection:   %.2f min\n', inj_time);
xline(inj_time, '--', 'Color', [0.5 0 0.8], 'LineWidth', 1.8, ...
    'Label', sprintf('Inj: %.2f', inj_time), ...
    'LabelVerticalAlignment', 'top');
drawnow;
disp('>>> Click 3: end of dead zone / signal stabilized');
[stab_time, ~] = ginput(1);
fprintf('Stabilized:  %.2f min\n', stab_time);
xline(stab_time, '--', 'Color', [1 0.5 0], 'LineWidth', 1.8, ...
    'Label', sprintf('Stable: %.2f', stab_time), ...
    'LabelVerticalAlignment', 'top');
ylims_raw = ylim;
patch([inj_time stab_time stab_time inj_time], ...
    [ylims_raw(1) ylims_raw(1) ylims_raw(2) ylims_raw(2)], ...
    [0.8 0.8 0.8], ...
    'FaceAlpha', 0.3, ...
    'EdgeColor', 'none', ...
    'HandleVisibility', 'off');
drawnow;
disp('>>> Click 4: end of post-treatment window');
[t_post_end, ~] = ginput(1);
fprintf('Post end:    %.2f min\n', t_post_end);
xline(t_post_end, '--', 'Color', [0.1 0.4 0.8], 'LineWidth', 1.8, ...
    'Label', sprintf('Post: %.2f', t_post_end), ...
    'LabelVerticalAlignment', 'top');
drawnow;
set(fig1, 'Visible', 'off');
%% -------- 5. Define analysis windows --------
mask_pre  = TimeW >= t_pre_start & TimeW <  inj_time;
mask_dead = TimeW >= inj_time    & TimeW <  stab_time;
mask_post = TimeW >= stab_time   & TimeW <= t_post_end;
mask_win  = TimeW >= t_pre_start & TimeW <= t_post_end;
fprintf('\nWindows:\n');
```

```
fprintf('  Pre:       %.2f - %.2f min  (%d points)\n', ...
        t_pre_start, inj_time, sum(mask_pre));
fprintf('  Dead zone: %.2f - %.2f min  (%d points) <- ignored\n', ...
        inj_time, stab_time, sum(mask_dead));
fprintf('  Post:      %.2f - %.2f min  (%d points)\n', ...
        stab_time, t_post_end, sum(mask_post));
%% -------- 6. Normalize MagLOV and keep TMRE raw --------
% MagLOV is normalized to the field-OFF baseline.
% In this envelope-based analysis, the lower envelope approximates
% the field-OFF fluorescence baseline.
F0_MagLOV = env_bot;
MagLOV_norm = (MagLOV ./ F0_MagLOV - 1) * 100;
% TMRE is kept as raw fluorescence intensity in arbitrary units.
% TMRE is plotted only for visualization and is not used for MFE calculation.
TMRE_raw = TMRE;
trend_pre_MagLOV  = mean(MagLOV_norm(mask_pre));
trend_post_MagLOV = mean(MagLOV_norm(mask_post));
trend_pre_TMRE    = mean(TMRE_raw(mask_pre));
trend_post_TMRE   = mean(TMRE_raw(mask_post));
%% -------- 7. dF/F and MFE per window --------
mean_top_pre  = mean(env_top(mask_pre));
mean_bot_pre  = mean(env_bot(mask_pre));
mean_top_post = mean(env_top(mask_post));
mean_bot_post = mean(env_bot(mask_post));
dFF_pre  = (mean_top_pre  - mean_bot_pre)  / mean_bot_pre  * 100;
dFF_post = (mean_top_post - mean_bot_post) / mean_bot_post * 100;
% Magnetic Fluorescence Effect: relative percent change after treatment
% compared with before treatment.
MFE_change = (dFF_post - dFF_pre) / dFF_pre * 100;
%% -------- 8. Plot normalized MagLOV + raw TMRE + dF/F --------
fig2 = figure('Name', 'Normalized MagLOV + Raw TMRE + Magnetic Modulation', ...
              'Position', [120 120 1100 550]);
subplot(2,1,1);
yyaxis left
plot(TimeW(mask_win), MagLOV_norm(mask_win), 'g-', ...
    'LineWidth', 1.3, ...
    'DisplayName', 'MagLOV norm.');
hold on;
yline(0, 'k--', 'LineWidth', 0.8, 'HandleVisibility', 'off');
ylims1 = ylim;
patch([inj_time stab_time stab_time inj_time], ...
     [ylims1(1) ylims1(1) ylims1(2) ylims1(2)], ...
     [0.8 0.8 0.8], ...
     'FaceAlpha', 0.3, ...
     'EdgeColor', 'none', ...
     'DisplayName', 'Dead zone');
ylabel('MagLOV \DeltaF/F (%)', 'FontSize', 11);
ax2 = gca;
ax2.YColor = [0.1 0.5 0.1];
yyaxis right
plot(TimeW(mask_win), TMRE_raw(mask_win), 'r-', ...
```

```
        'LineWidth', 1.5, ...
        'DisplayName', 'TMRE');
    ylabel('TMRE F.I. (arb. units)', 'FontSize', 11);
    ax2.YColor = [0.8 0.1 0.1];
    xline(t_pre_start, '--', 'Color', [0.2 0.6 0.2], ...
          'LineWidth', 1.2, 'HandleVisibility', 'off');
    xline(inj_time, '--', 'Color', [0.5 0 0.8], ...
          'LineWidth', 1.5, 'HandleVisibility', 'off');
    xline(stab_time, '--', 'Color', [1 0.5 0], ...
          'LineWidth', 1.5, 'HandleVisibility', 'off');
    xline(t_post_end, '--', 'Color', [0.1 0.4 0.8], ...
          'LineWidth', 1.2, 'HandleVisibility', 'off');
    xlabel('Time (min.)', 'FontSize', 11);
    title('MagLOV normalized to field-OFF baseline; TMRE raw intensity', 'FontSize', 11);
    legend({'MagLOV norm.', 'Dead zone', 'TMRE'}, ...
           'Location', 'northeast', 'FontSize', 10);
    set(gca, 'FontSize', 10);
    grid on; box on;
    hold off;
    subplot(2,1,2);
    plot(TimeW(mask_win), dFF_cont(mask_win), 'b-', 'LineWidth', 1.3);
    hold on;
    ylims2 = ylim;
    patch([inj_time stab_time stab_time inj_time], ...
          [ylims2(1) ylims2(1) ylims2(2) ylims2(2)], ...
          [0.8 0.8 0.8], ...
          'FaceAlpha', 0.3, ...
          'EdgeColor', 'none');
    yline(dFF_pre, 'g--', 'LineWidth', 1.5, ...
          'Label', sprintf('\\DeltaF/F pre = %.2f%%', dFF_pre), ...
          'LabelHorizontalAlignment', 'left');
    yline(dFF_post, 'b--', 'LineWidth', 1.5, ...
          'Label', sprintf('\\DeltaF/F post = %.2f%%', dFF_post), ...
          'LabelHorizontalAlignment', 'left');
    yline(0, 'k--', 'LineWidth', 0.8);
    xline(t_pre_start, '--', 'Color', [0.2 0.6 0.2], 'LineWidth', 1.2);
    xline(inj_time, '--', 'Color', [0.5 0 0.8], 'LineWidth', 1.5);
    xline(stab_time, '--', 'Color', [1 0.5 0], 'LineWidth', 1.5);
    xline(t_post_end, '--', 'Color', [0.1 0.4 0.8], 'LineWidth', 1.2);
    xlabel('Time (min.)', 'FontSize', 11);
    ylabel('\DeltaF/F (%)', 'FontSize', 11);
    title('Continuous magnetic-field modulation from envelope', 'FontSize', 11);
    set(gca, 'FontSize', 10);
    grid on; box on;
    hold off;
    %% -------- 9. Results figure --------
    fig3 = figure('Name', 'Results Summary', ...
                  'Position', [900 300 500 460]);
    axis off;
    results_str = {
       sprintf('File: %s', fname),                                  ...
```

```
    ' ',                                                 ...
    sprintf('Pre start:   %.2f min', t_pre_start),                    ...
    sprintf('Injection:   %.2f min', inj_time),                       ...
    sprintf('Stabilized:  %.2f min', stab_time),                      ...
    sprintf('Post end:    %.2f min', t_post_end),                     ...
    sprintf('Dead zone:   %.2f min', stab_time - inj_time),           ...
    ' ',                                                 ...
    '--- Normalized signal (MagLOV) ---',                             ...
    sprintf('Pre  mean: %+.2f%%', trend_pre_MagLOV),                  ...
    sprintf('Post mean: %+.2f%%', trend_post_MagLOV),                 ...
    sprintf('Change:    %+.2f%%', trend_post_MagLOV - trend_pre_MagLOV), ...
    ' ',                                                 ...
    '--- TMRE raw fluorescence ---',                                  ...
    sprintf('Pre  mean: %.2f a.u.', trend_pre_TMRE),                  ...
    sprintf('Post mean: %.2f a.u.', trend_post_TMRE),                 ...
    sprintf('Change:    %.2f a.u.', trend_post_TMRE - trend_pre_TMRE),   ...
    ' ',                                                 ...
    '--- Magnetic-field modulation ---',                              ...
    'Formula: dF/F = (mean_top - mean_bot) / mean_bot * 100',         ...
    sprintf('dF/F pre:    %+.2f%%', dFF_pre),                         ...
    sprintf('dF/F post:   %+.2f%%', dFF_post),                        ...
    sprintf('MFE change:  %+.2f%%', MFE_change),                      ...
};
for i = 1:length(results_str)
    is_header = startsWith(results_str{i}, '---');
    text(0.05, 1 - i*0.043, results_str{i}, ...
        'Units', 'normalized', ...
        'FontName', 'Courier', ...
        'FontSize', 9.2, ...
        'FontWeight', iif(is_header, 'bold', 'normal'), ...
        'Color', iif(is_header, [0 0 0.7], [0 0 0]));
end
title('Results Summary', 'FontSize', 12, 'FontWeight', 'bold');
%% -------- 10. Print to Command Window --------
fprintf('\n');
fprintf('==============================================\n');
fprintf('  RESULTS\n');
fprintf('==============================================\n');
fprintf('File:        %s\n', fname);
fprintf('Pre start:   %.2f min\n', t_pre_start);
fprintf('Injection:   %.2f min\n', inj_time);
fprintf('Stabilized:  %.2f min\n', stab_time);
fprintf('Post end:    %.2f min\n', t_post_end);
fprintf('Dead zone:   %.2f min\n', stab_time - inj_time);
fprintf('\n--- 1. Normalized signal (MagLOV) ---\n');
fprintf('Pre  mean: %+.2f%%\n', trend_pre_MagLOV);
fprintf('Post mean: %+.2f%%\n', trend_post_MagLOV);
fprintf('Change:    %+.2f%%\n', trend_post_MagLOV - trend_pre_MagLOV);
fprintf('\n--- 2. TMRE raw fluorescence ---\n');
fprintf('Pre  mean: %.2f a.u.\n', trend_pre_TMRE);
fprintf('Post mean: %.2f a.u.\n', trend_post_TMRE);
```

```
fprintf('Change:    %.2f a.u.\n', trend_post_TMRE - trend_pre_TMRE);
fprintf('\n--- 3. Magnetic-field modulation ---\n');
fprintf('Formula: dF/F = (mean_top - mean_bot) / mean_bot x 100\n');
fprintf('dF/F_pre    = %+.2f%%\n', dFF_pre);
fprintf('dF/F_post   = %+.2f%%\n', dFF_post);
fprintf('MFE_change  = %+.2f%%\n', MFE_change);
fprintf('========================================\n');
%% -------- 11. Save figures --------
fig2_name = fullfile(fpath, [fname_base '_signals.pdf']);
fig3_name = fullfile(fpath, [fname_base '_results.pdf']);
exportgraphics(fig2, fig2_name, 'ContentType', 'vector');
exportgraphics(fig3, fig3_name, 'ContentType', 'vector');
fprintf('Figure 2 saved: %s\n', fig2_name);
fprintf('Figure 3 saved: %s\n', fig3_name);
%% -------- 12. Save to Excel --------
xlsx_name = fullfile(fpath, [fname_base '_results.xlsx']);
results_table = table( ...
   {fname_base}, ...
   t_pre_start, inj_time, stab_time, t_post_end, ...
   stab_time - inj_time, ...
   trend_pre_MagLOV, trend_post_MagLOV, trend_post_MagLOV - trend_pre_MagLOV, ...
   trend_pre_TMRE, trend_post_TMRE, trend_post_TMRE - trend_pre_TMRE, ...
   dFF_pre, dFF_post, MFE_change, ...
   'VariableNames', { ...
      'File', ...
      'Pre_start_min', 'Injection_min', 'Stabilized_min', 'Post_end_min', ...
      'Dead_zone_min', ...
      'MagLOV_pre_mean_pct', 'MagLOV_post_mean_pct', 'MagLOV_change_pct', ...
      'TMRE_pre_mean_au', 'TMRE_post_mean_au', 'TMRE_change_au', ...
      'dFF_pre_pct', 'dFF_post_pct', 'MFE_change_pct' ...
   });
if isfile(xlsx_name)
   existing = readtable(xlsx_name, 'VariableNamingRule', 'preserve');
   combined = [existing; results_table];
   writetable(combined, xlsx_name);
   fprintf('Row appended to existing file: %s\n', xlsx_name);
else
   writetable(results_table, xlsx_name);
   fprintf('New Excel file created: %s\n', xlsx_name);
end
%% -------- Helper function --------
function out = iif(cond, a, b)
   if cond
      out = a;
   else
      out = b;
   end
end
```

### Epifluorescence experiments

The remaining portion of this Supplementary Information provides a detailed report of epifluorescence experiments. In contrast to the main text, which reports confocal measurements, epifluorescence is essentially a steady state measurement compared to the timescale of the confocal. Therefore, the two experimental conditions are distinct. Despite these differences, the technique also demonstrates clear sensitivity of mtMagLOV2 magnetic field dependent fluorescence in response to pharmacological manipulation of the metabolic state of the mitochondria.

**Modulation of mtMagLOV2 Fluorescence by Magnetic Fields (Epi)**

Upon application of the magnetic field, we observed distinct oscillations in the fluorescence intensity of the mtMagLOV2 sensor. The mtMagLOV2 domain exhibited periodic "dimming and brightening" synchronized with the 5-second magnetic pulses (Figure S13).

Quantitative analysis of the mtMagLOV2 domain's response revealed significant fluctuations in the normalized fluorescence change, calculated as:

$$\frac{\Delta F}{F}(\%) = \frac{F - F_0}{F_0} \times 100$$

where $F$ represents the intensity at time $t$ and $F_0$ is the baseline intensity. The $\Delta F/F$ values fluctuated in direct response to the magnetic field cycles, confirming the sensor's sensitivity to magnetic stimuli in live HeLa-mtMagLOV2 cells [1].

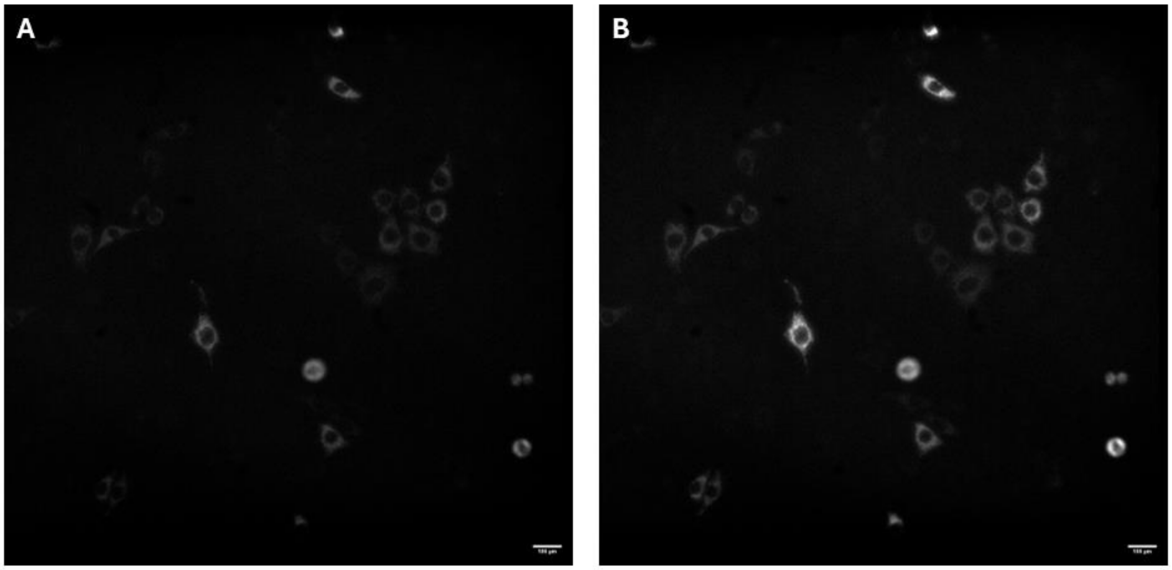


**Figure S13. HeLa-mtMagLOV2 Fluorescence Modulation.** (A) Coil ON: Application of the magnetic field induces fluorescence dimming (quenching). (B) Coil OFF: Removal of the field results in brightening as the signal returns to baseline. The observed oscillations are synchronized with the 5-second magnetic pulses.

**Demonstration of quantum sensing of metabolic state through pharmacological manipulation**

To evaluate the real-time monitoring capabilities of the mtMagLOV2 sensor under dynamic metabolic conditions, we performed a sequential mitochondrial stress test analogous to a Seahorse XF assay. HeLa-mtMagLOV2 cells were subjected to three successive pharmacological perturbations during a single imaging session: oligomycin (1 µM), FCCP (10 µM), and rotenone (10 µM) mixture (Figure S14). Each reagent was added at approximately 7-minute intervals to allow for signal stabilization and the capture of metabolic transition kinetics.

To quantify the impact of pharmacological perturbation on magnetic responsiveness, the Magnetic Field Effect (MFE) was calculated using the following equation: $MFE =$

$\left(\frac{(\Delta F/F)_{after} - (\Delta F/F)_{before}}{(\Delta F/F)_{before}}\right) \times 100$ where $((\Delta F/F)_{before})$ represents the baseline responsiveness before treatment and $((\Delta F/F)_{after})$ represents the responsiveness at each 7-minute interval post-treatment.

Initial treatment with oligomycin, an ATP synthase inhibitor, resulted in a modest increase in magnetic field effect (MFE = 36%), likely reflecting a slight hyperpolarization of the mitochondrial membrane potential ($\Delta\Psi_m$) as proton flow through the ATP synthase ($F_0F_1$) complex was restricted.

The most dramatic shift occurred following the addition of the uncoupler FCCP. This stage induced rapid signal quenching, reaching an MFE of 56%. This pronounced attenuation correlates with the maximal depolarization of the mitochondrial inner membrane and the collapse of the proton gradient, proving that mtMagLOV2 is acutely sensitive to extreme shifts in mitochondrial functional state.

Finally, rotenone was added, and the MFE shifted to 49%. Rather than a novel effect from rotenone itself, this slight decrease from the FCCP peak represents the time-dependent decay of the initial inhibition and a slow recovery phase as the membrane adapts. Together, these results demonstrate that mtMagLOV2 provides a robust, real-time readout capable of resolving multiple metabolic phases within the same cells.

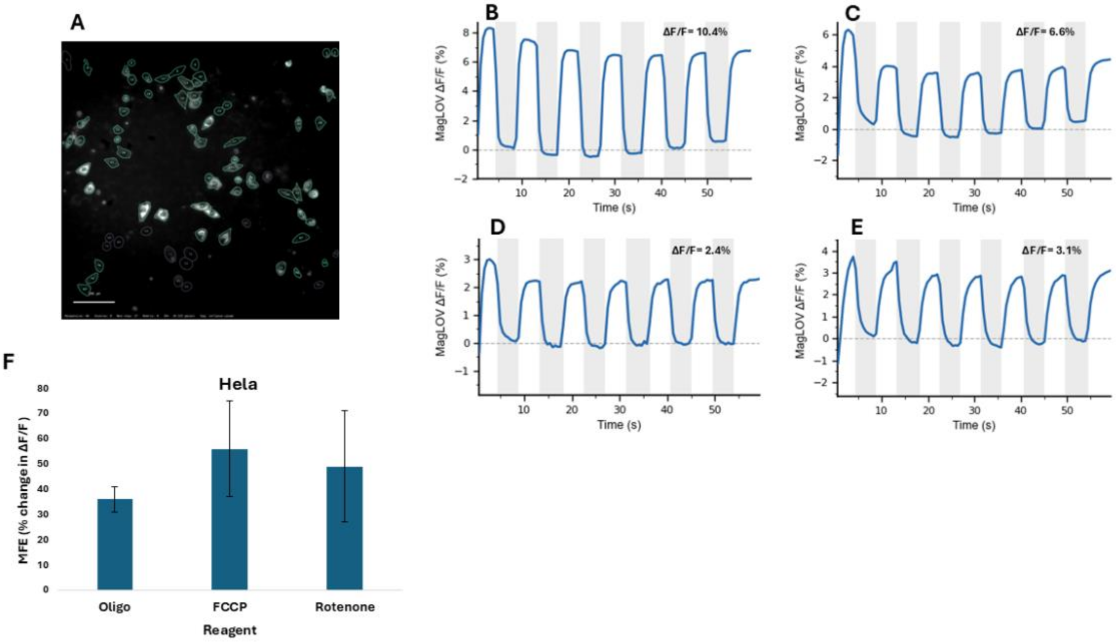


**Figure S14. Sequential mitochondrial stress test in HeLa-mtMagLOV2 cells. The bar chart and time-course profiles illustrate the responses following the sequential addition of respiratory modulators at 7-minute intervals.** (A) Segmentation of cells responsive to magnetic stimulation ($\Delta F/F$). (B) Baseline: Initial magnetic responsiveness recorded prior to treatment. (C) Hyperpolarization: Positive shift in the magneto-optical response following the addition of oligomycin (1 µM). (D) Depolarization: Significant signal quenching and collapse of the mitochondrial membrane potential $\Delta\Psi_m$ after the addition of FCCP (10 µM). (E) Recovery Phase: Slow, time-dependent recovery of the response after adding rotenone (10 µM), confirming full suppression of the electron transport chain (ETC). (F) Statistical summary of the sequential mitochondrial stress test showing the percentage change in MFE across treated HeLa-mtMagLOV2 cells. Values represent the mean calculated from a total of 25 responsive cells. The error bars represent the standard deviation (SD) from n=3 independent experiments.
